# Temporal control of the integrated stress response by a stochastic molecular switch

**DOI:** 10.1101/2022.01.01.474691

**Authors:** Philipp Klein, Stefan M. Kallenberger, Hanna Roth, Karsten Roth, Thi Bach Nga Ly-Hartig, Vera Magg, Janez Aleš, Soheil Rastgou Talemi, Yu Qiang, Steffen Wolf, Olga Oleksiuk, Roma Kurilov, Barbara Di Ventura, Ralf Bartenschlager, Roland Eils, Karl Rohr, Fred A. Hamprecht, Thomas Höfer, Oliver T. Fackler, Georg Stoecklin, Alessia Ruggieri

## Abstract

Stress granules (SGs) are formed in the cytosol as an acute response to environmental cues and activation of the integrated stress response (ISR), a central signaling pathway controlling protein synthesis. Using chronic virus infection as stress model, we previously uncovered a unique temporal control of the ISR resulting in recurrent phases of SG assembly and disassembly. Here, we elucidate the molecular network generating this fluctuating stress response, by integrating quantitative experiments with mathematical modeling, and find that the ISR operates as a stochastic switch. Key elements controlling this switch are the cooperative activation of the stress-sensing kinase PKR, the ultrasensitive response of SG formation to the phosphorylation of the translation initiation factor eIF2α, and negative feedback via GADD34, a stress-induced subunit of protein phosphatase 1. We identify GADD34 mRNA levels as the molecular memory of the ISR that plays a central role in cell adaptation to acute and chronic stress.

## Introduction

Mammalian cells maintain cellular homeostasis and promote survival by integrating a multitude of extrinsic and intrinsic signals into a spatially and temporally regulated response. Adverse conditions such as oxidative stress, endoplasmic reticulum (ER) stress and virus infections are detected by four specialized cytosolic sentinels that belong to the eukaryotic translation initiation factor 2-alpha (eIF2α) kinase family of serine/threonine kinases. They initiate the integrated stress response (ISR) by immediately phosphorylating eIF2α (1), interfering with formation of the eIF2-GTP-tRNAiMet ternary complex and thus inhibiting translation initiation (2). As a consequence, polysomes disassemble and non-translating messenger RNAs (mRNAs) phase separate together with RNA-binding proteins into membraneless biomolecular condensates called stress granules (SGs) (3). Among the eIF2α-kinases, protein kinase R (PKR) is an interferon (IFN) induced kinase (4, 5) that mediates translation suppression in response to replication of many RNA viruses. PKR activation was initially described to result from binding to double-stranded (ds) RNA of diverse viral origin such as viral replication intermediates or transcripts containing stem loop structures generated during infection (6). However, other sources of cellular dsRNAs, including mitochondrial dsRNA, circular RNAs and small nucleolar RNAs, also control its activation (7). PKR dimerization, which is required for its activation, leads to autophosphorylation of the PKR kinase domain, most notably at threonine 446, and to structural rearrangements that facilitate binding to eIF2α (8–11).

Translation shutdown feeds back into the regulation of eIF2α phosphorylation by upregulating Growth Arrest and DNA-Damage-inducible 34 (GADD34), a stress-induced regulatory subunit of protein phosphatase 1 (PP1) (12) that acts as antagonist of the eIF2α-kinases and mediates eIF2α dephosphorylation. This system allows the cell to integrate different cues and adjust the degree of translational suppression during the course of stress responses. We previously discovered an extreme case of such temporal control, whereby hepatitis C virus (HCV), a major human pathogen causing chronic liver infection, induces a sustained cellular stress response characterized by recurrent alternating translational off- and on-states that are, respectively, accompanied by the assembly and disassembly of SGs. Notably, the duration of off- and on-states was highly variable in individual cells and not synchronous between cells. At the molecular level, the fluctuating SG response was dependent on PKR and GADD34, and strongly enhanced by treatment of HCV-infected cells with IFN-α (13). However, the molecular mechanisms underlying the apparently oscillatory nature of this response have not been elucidated.

Oscillations are observed in numerous biological systems including circadian clock, metabolism, signaling, cell division and development (14, 15). Oscillating systems are characterized by signal amplitude and periodicity covering a large range of time scales from seconds in the case of calcium signaling (16) to hours, e.g. for bursts of tumor suppressor p53 expression upon DNA damage (17) or for the circadian clock (18). However, not every dynamic biochemical system characterized by recurring bursts in signals necessarily constitutes an oscillator. Biochemical processes are subjected to biological noise that impacts the kinetics of cellular reactions, particularly involving factors of low abundance, and influences the behavior of the response, thus generating cell-to-cell variability (19–23).

Here, we set out to model the recurrent alternating stress response, using HCV infection as a model for chronic stress response, with the aim to understand the molecular mechanisms that govern its establishment and maintenance over long periods of time. To this end, we developed a quantitative deterministic mathematical model that describes the components and reactions involved in the ISR to oxidative stress, ER stress and more specifically to the dsRNA-induced SG response. To approximate our cell system as closely as possible, we experimentally determined a large set of key species concentration ranges using bulk and single-cell methods. While our observations first suggested that the ISR signaling network shares common features of an oscillator, we found that HCV-induced SG fluctuations result from repetitive stochastic transitions between On- and Off-states that are regulated by cell-to-cell variability in PKR and dsRNA concentrations. Importantly, our results identified GADD34 mRNA levels as the molecular memory of the ISR, which plays a crucial role in adaptation to acute and chronic stress.

## Results

### The fluctuating SG response to HCV infection exhibits a stochastic switch behavior with memory

To accurately characterize the dynamics of the cellular stress response to HCV infection, we monitored SG formation using long-term live cell microscopy with high time resolution. Human hepatocarcinoma-derived Huh7 cells stably expressing the SG protein T-cell-restricted intracellular antigen-1 (TIA-1) fused to the yellow fluorescent protein (YFP) were infected with HCV-like particles (HCV_TCP_), which encode the fluorescent protein mCherry fused to the viral non-structural protein NS5A and Renilla luciferase for the measurement of viral replication (fig. S1A). As replication reached its maximum, i.e. 48 hours post infection (fig S1B), SG response dynamics was monitored for 72 hours in infected cells that were either left untreated or treated with IFN-α (fig. S1C). Images were acquired with 15-minute intervals to ensure the detection of short SG phases (24) while limiting photo-toxicity (Figure 1A, Movie S1 and S2). Automated single-cell tracking in such long-term time-lapses is challenging due to the relatively long interval between acquisition frames, cell division events, and the increasing cell density towards the end of the acquisition period (25, 26). We therefore developed a semi-automated image processing pipeline primarily based on the machine-learning toolkit ilastik (27) and analyzed the appearance and disappearance of SGs (here defined as SG-On and SG-Off phases, respectively) in single cells over time (Fig. 1, B and C, and fig. S1D). As previously observed (13), SGs were extremely rare in uninfected cells (fig. S2A). In HCV-infected cells that were not exposed to IFN-α, the majority of cells displayed 0.5 to 1.5 SG-On phases per day (Fig. 1D left graph, fig. S2A). This number increased upon addition of IFN-α, with the majority of cells having 1 to 2.5 SG-On phases per day (Fig. 1D right graph). Importantly, SG phases did not exhaust over time (Fig. 1C). The comparative analysis of SG properties and infection level – as measured by mCherry-NS5A fluorescence intensity – revealed a slight positive correlation between the infection level and the total number of SGs as well as the number of SG phases, and a negative correlation with the SG integral (total duration of SG presence) and duration (mean phase duration) (fig. S2B), suggesting a possible role of SGs in host defense as reported for other viral infections (7).

**Fig. 1.**
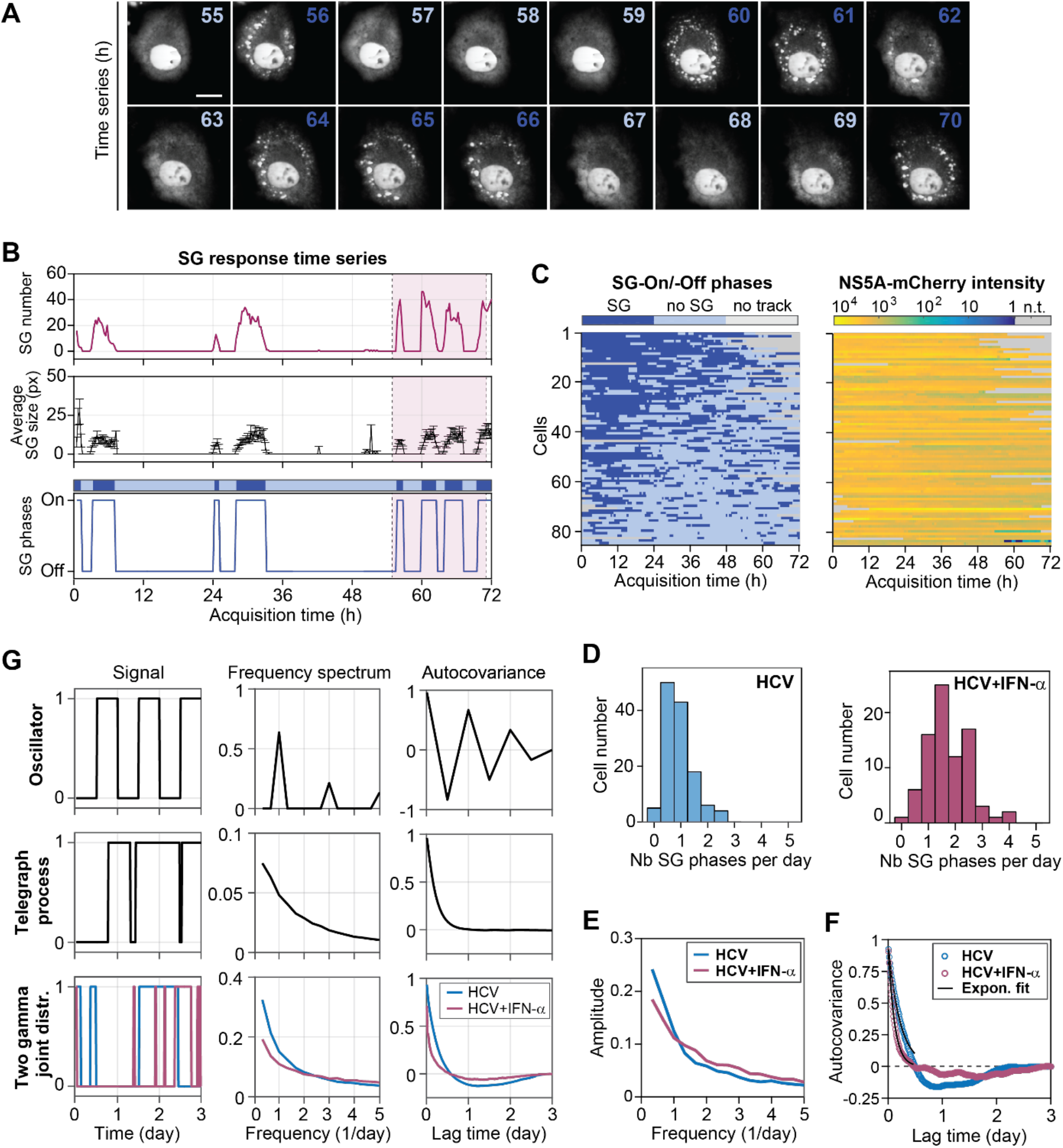
Time-resolved analysis of HCV-induced SG phases. Huh7 YFP-TIA1 cells were infected with HCV_TCP_ for 48 h and SG dynamics monitored for 72 h using live-cell time-lapse microscopy. (**A**) Still images of a cropped section of the time-lapse movie of HCV-infected cells treated with IFN-α (YFP-channel, movie S1). A representative cell is shown with 1-h interval for the time period 55-70 h after the start of the acquisition. The color of the time label indicates SG-On phases in dark blue and SG-Off phases by light blue. Scale bar, 25 μm. (**B**) Example of time-lapse analysis output for the cell shown in (A). The number of SGs (upper panel) and the average SG size in pixel (middle panel) were analyzed for each frame (15-min interval) and allowed defining SG-On and SG-Off phases (lower panel). The pink shaded area corresponds the time period 55-70 h shown in (A). A schematic of the SG response time series is shown in the top with dark blue regions representing SG-On phases and light blue regions SG-Off phases. (**C**) Analysis of multiple single cell SG response time series. Cells used for analysis were tracked for at least 48 h. Left panel, SG-On and SG-Off phases (YFP-TIA1 channel, n=85). Right panel, infection levels as measured by NS5A-mCherry signal intensity of the corresponding cells. Single-cell SG response time series were sorted based on the total stress duration, with cell 1 on the top having more stress than the cells on the bottom of the panel. (**D**) Quantification of the number of SG phases per day in the absence (left panel, HCV) or presence of IFN-α (right panel, HCV + IFN-α). (**E**) Average frequency spectra (Fourier transforms) of experimental single-cell SG response time series. (**F**) Autocorrelation functions of experimental single-cell SG response time series. (**G**) To evaluate the presence of possible periodicity in the experimental single-cell SG response time series, two scenarios, an oscillator and a random telegraph process, were simulated. Panels on the left represent the type of signal response, in the middle the frequency spectrum and on the right the autocovariance function (frequency spectrum and autocovariance of the telegraph process were calculated as averages of n=500 realizations). The observed frequency spectra and autocovariance functions in HCV-infected as well as HCV-infected and IFN-α-treated cells can be well reproduced by sampling from joint gamma distributions assuming a random process with short and long SG-On and SG-Off phase lengths.

SG fluctuations could result from an oscillator mechanism, which has a fixed period possibly disturbed by noise effects, or from the random switching between SG-On and SG-Off states, a so-called random telegraph process, which has no fixed period because the moments in which cells switch from one state to the other are not influenced by the time already elapsed. To get more insight into the quality of the SG fluctuations, we simulated these models computationally and compared the results to experimental single cell SG response time series (Fig. 2G, Supplementary Text section 1). We analyzed the degree of SG phase periodicity from each time series using the frequency spectrum (obtained via the Fourier transform) and the autocovariance function. In contrast to a simulated oscillator distinguished by distinct peaks in the frequency spectrum and an autocovariance function alternating between positive and negative values (Fig. 2G, upper), SG response time series revealed the absence of a defined frequency, both in presence and absence of IFN-α (Fig. 1E). The autocovariance function showed a continuous decline similar to that of the telegraph process (Fig. 2G, middle). However, unlike the telegraph process, it even reached slightly negative values, particularly for untreated cells (Fig. 1F), suggesting a negative correlation between SG-On and SG-Off phase length. The analysis of the experimental SG phase length distribution in HCV-infected cells indicated preferred phase lengths and validated this correlation, whose distribution was well described by two joint gamma distributions (fig. S2C). Simulations of such a gamma process (Fig. 2G, bottom) reliably recapitulated the frequency spectra and autocovariance functions of the experimental observations (Fig. 1, E and F). In addition, calculated autocovariance exponential decay times were reduced upon IFN-α treatment (from four to two hours, fig. S2D), reflecting the occurrence of shorter SG-On and SG-Off phases under this condition. Taken together, our results suggest that SG fluctuations are described neither by an oscillator nor by a simple telegraph process, but rather by a stochastic switching process with memory. Specifically, the moment at which cells switch SG phase is negatively correlated with the duration of preceding SG phase.

**Fig. 2.**
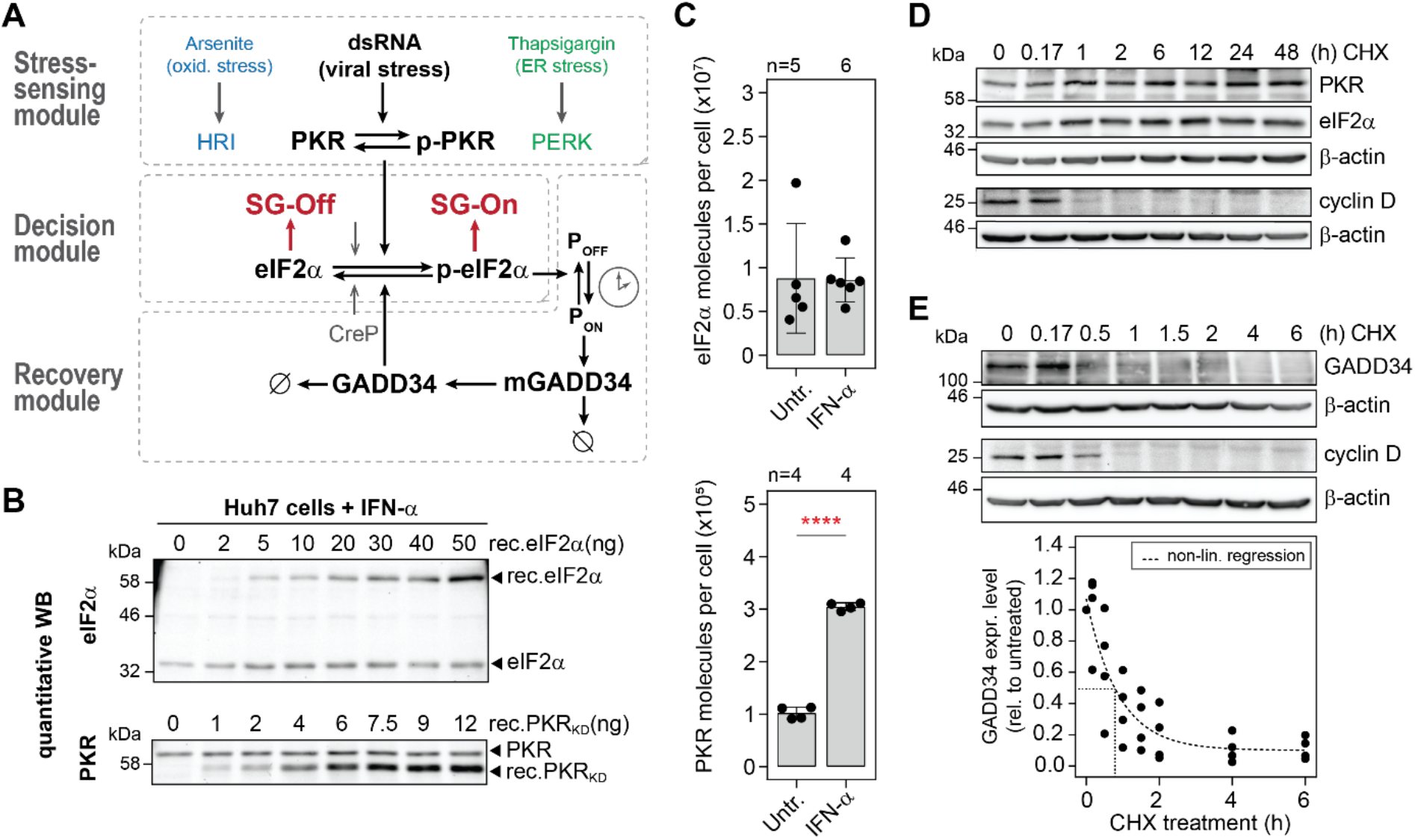
Deterministic mathematical model of the cellular ISR. (**A**) Schematic representation of the parameters and reactions included in the mathematical model. The stress-sensing module represents the activation of stress kinases. PKR is activated by binding to dsRNA (p-PKR) (viral stress), HRI by arsenite treatment (oxidative stress) and PERK by thapsigargin (ER stress). Active stress kinases signal to the decision module, where upon crossing a certain threshold p-eIF2α levels will trigger SG formation (SG-On) in response to the translational attenuation. Elevated p-eIF2α levels activate the recovery module consisting of the GADD34 negative feedback loop i.e. *ppp15R1a* promoter (P_OFF_ to P_ON_), GADD34 transcription (mGADD34) and protein synthesis (GADD34) with time delay (clock symbol). In turn, GADD34 dephosphorylates eIF2α and thereby resumes translation (SG-Off). Grey arrows indicate a basal eIF2α dephosphorylation by CReP, the constitutive regulatory subunit of PP1. Ø, degradation rates. (B and C) Absolute quantification of eIF2α and PKR mean molecule numbers in Huh7 YFP-TIA1 cells, in absence and presence of IFN-α. (**B**) Representative quantitative Western blot analysis of eIF2α and PKR. Cell lysates of a determined cell number were spiked with varying amounts of recombinant GST-tagged eIF2α (rec. eIF2α) or GST-tagged PKR kinase domain (rec. PKR_KD_). (**C**) Estimated eIF2α and PKR mean molecule number per cell (± SD). Number of repeats (n) and statistical significance compared to untreated cells (untr.) are indicated; ****p<0.0001. (**D and E**) Determination of protein half-lives in CHX pulse experiments. Levels were normalized to β-actin expression. Shown are representative Western blot analyses. Cyclin D expression was used as control of a short-lived protein. PKR and eIF2α expression was measured in Huh7 cells treated for 48 h with CHX (n=3). For quantifications see fig. S3E (D). GADD34 turnover was measured in Huh7 GADD34 cells treated with CHX for 6 h (n=4) (E). GADD34 expression is shown relative to untreated cells. The black line depicts best non-linear fit (one-phase decay).

### Network topology of the elF2α-mediated signaling pathway

We previously uncovered that SG fluctuations during chronic HCV infection result from antagonistic activities of PKR and GADD34-PP1 on eIF2α phosphorylation (13). To understand how these main regulators shape this peculiar SG response dynamics in single cells, we developed a deterministic mathematical model based on ordinary differential equations (ODEs) that describes the eIF2α-dependent signaling pathway in response to stress, particularly to dsRNA, assuming a homogenous and synchronized cell population. The model was structured into three independent but connected modules (Fig. 2A). The upper stress-sensing module described the activation of the stress kinase PKR by dsRNA. Of note, in this module PKR is interchangeable with two other eIF2α-kinases, heme-regulated inhibitor (HRI) and PKR-like (ER) kinase (PERK), that act upstream of eIF2α phosphorylation. The core decision module was composed of eIF2α whose phosphorylation leads to translation repression and SG formation. The downstream recovery module described the transcriptional and translational activation of GADD34, which represents a negative feedback loop that promotes SG disassembly, translation re-initiation and return to homeostasis through dephosphorylation of eIF2α.

To constrain the mathematical model for making quantitative predictions, we experimentally determined several of its parameters. We first measured the mean abundance per cell of the key proteins i.e. PKR, eIF2α and GADD34. Protein amounts were measured by quantitative Western blot analysis using standard dilutions of recombinant GST-tagged eIF2α and PKR as reference (Fig. 2B and fig S3A). As expected for a constitutively expressed protein, the levels of eIF2α did not vary upon addition of IFN-α. However, the levels of PKR increased by three-fold (Fig. 2C). By measuring the mean volume of Huh7 cells (fig. S3B), we deduced the corresponding mean concentrations of approx. 2.24 mM for eIF2α, 25.24 nM for PKR in untreated cells and 75.48 nM for PKR in IFN-α treated cells, respectively. As no basal expression of GADD34 was detected in unstressed cells, we measured the amount of GADD34 that could be induced upon treatment with thapsigargin, a chemical inducer of ER stress. Since endogenous, overexpressed and even recombinant GADD34 are extremely sensitive to degradation (fig. S3C), we used an indirect approach and first quantified GADD34 levels in Huh7 cells stably expressing GADD34-eGFP using a titration of recombinant eGFP. These calibrated lysates served as reference for GADD34 measurements in Huh7 cells treated with thapsigargin. Thereby, we estimated that the mean concentration of stress-induced GADD34 typically reaches 8×10^3^ molecules per cell (equivalent to 2 nM) (fig. S3D).

Lastly, degradation rates of species involved in feedback loops are linked to time delays and thereby facilitate oscillations (28). We hence determined protein turnover by cycloheximide (CHX) chase experiments using the short-lived protein cyclin D as a control. PKR and eIF2α were found to be long-lived proteins (Fig. 2D, fig. S3E) whose degradation rates are therefore negligible for the model. In turn, stably overexpressed GADD34 was labile (Fig. 2E) in agreement with previous reports (29).

Comparison of two possible degradation models, Michaelis-Menten kinetics and exponential decay, indicated that GADD34 degradation is described more accurately by exponential decay - implying that GADD34 degradation was not saturated - with an estimated half-life of approx. 37 minutes (Supplementary Text section 3).

Together, this first set of quantitative measurements yielded mean concentrations of the main model species that will serve as boundary conditions of the deterministic mathematical model. In addition, we found PKR and eIF2α turnover to be negligible parameters for the model, whereas GADD34 is short-lived, a feature that might facilitate the generation of oscillations.

### Dose-response analyses reveal the switch-like behavior of SG formation

Following the quantification of these key parameters, we sought to characterize the central decision module of the model and determined the level of p-eIF2α at which cells transition from a SG-Off to a SG-On phase. To this end, we performed dose-response analyses and measured the percentage of SG-positive cells at the population level as a function of dsRNA concentration. Since the precise nature and length of dsRNA intermediates produced during HCV infection are unknown, we synthesized *in vitro* dsRNAs of variable lengths (100, 200 and 400 bp; fig. S4A) and compared their potential to activate PKR upon transfection into cells (fig. S4B). PKR activity was assessed by measuring (i) eIF2α and PKR phosphorylation levels using Western blot analysis (Fig. 3, A and B, and fig. S4C), (ii) the percentage of p-eIF2α relative to total eIF2α using Phos-tag gel analysis (Fig. 3, A and B), and (iii) the percentage of SG-positive cells by immunofluorescence analysis (Fig. 3C). Of note, although 400-bp dsRNA triggered the highest levels of PKR activation, it formed additional secondary structures under non-reducing conditions (fig. S4A). We thus chose 200-bp dsRNA for the following dose-response experiments. At best, dsRNA transfection induced 25% eIF2α phosphorylation (Fig. 3B) and 40% SG-positive cells (Fig. 3C). PKR activation followed a bell-shape curve as reflected by a significant decrease in the levels of phosphorylated PKR (p-PKR), p-eIF2α, and in the number of SG-positive cells, at higher dsRNA concentrations. This is consistent with earlier reports (30) and suggests inhibition of PKR at high levels of dsRNA.

**Fig. 3.**
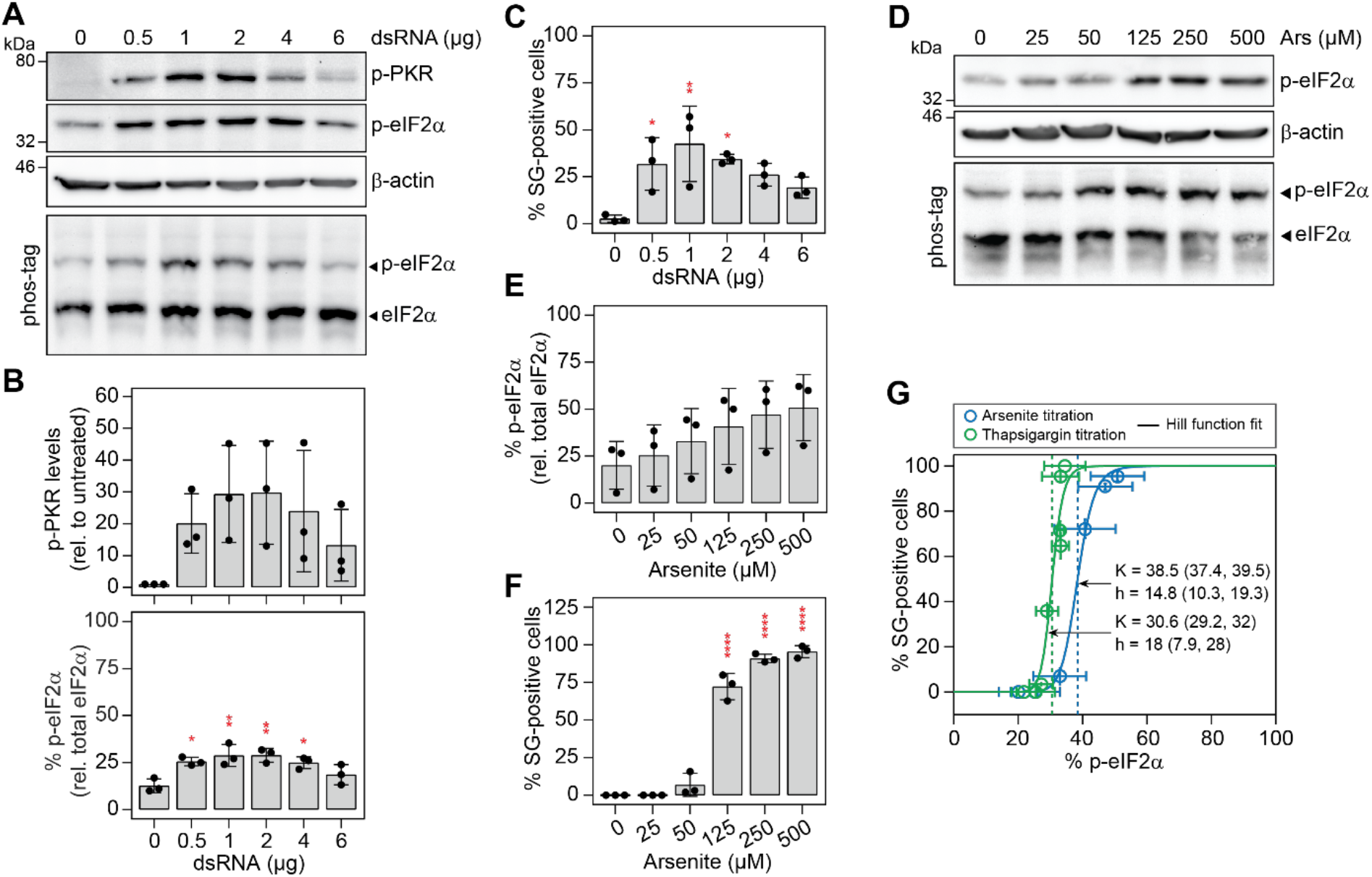
SG formation is a switch-like process. (**A to C**) Activation of PKR in Huh7 YFP-TIA1 cells transfected with increasing amounts of 200-bp dsRNA (n=3). (A) Representative Western blot analysis of p-PKR and p-eIF2α expression levels. Expression levels of β-actin served as loading control. The percentage of p-eIF2α was analyzed by Phos-tag polyacrylamide gel. (B) Shown are the quantifications of mean p-PKR expression levels (± SD) normalized to the loading control and relative to untreated cells (upper panel) and quantifications of the mean p-eIF2α percentage relative to the total eIF2α (± SD). Statistical significance is indicated compared to untreated cells. (C) The presence of SGs in transfected cells was analyzed by fluorescence microscopy. For each condition, more than 100 cells were analyzed for the presence of SGs using the YFP-TIA1 signal. Shown are mean percentages ± SD. Statistical significance is indicated compared to untreated cells. *p<0.05, **p<0.01. (**D to F**) Induction of oxidative stress in Huh7 YFP-TIA1 cells by treatment with increasing concentrations of arsenite for 45 min (n=3). (D) Representative Western blot and Phos-tag analyses of p-eIF2α levels. Shown are mean percentages ± SD of p-eIF2α (E) and SG-positive cells (F). For each condition, more than 100 cells were analyzed in fluorescence microscopy. Statistical significance is indicated compared to untreated cells; ****p<0.0001. (**G**) Dose-response analysis and determination of the p-eIF2α level that results in formation of SGs in 50% of cells upon treatment with arsenite (related to panel F; n=3) and thapsigargin (related to fig. S4, E to H; n=3). Fifty percent of cells form SGs when in mean 30.6 and 38.5 % eIF2α is phosphorylated in response to arsenite and thapsigargin treatment, respectively. Hill-coefficients (h=14.8 for arsenite, h=18.0 for thapsigargin).

These results combined with the possibility that the transfection approach does not homogeneously deliver dsRNA to all cells represented a significant technical limitation to determine the half-maximal dose of p-eIF2α causing SG formation. Therefore, we turned to chemical stress inducers and determined p-eIF2α thresholds upon treatment with sodium arsenite (Fig.3, D and E, fig. S4D) and thapsigargin (fig S4, E to G), which trigger HRI-dependent oxidative stress and PERK-dependent ER stress, respectively (31), and synchronously induced SG formation in all cells at higher concentrations (Fig. 3F, fig. S4H). When the percentage of SG-positive cells was plotted against the percentage of p-eIF2α (Fig. 3G), we observed an ultrasensitive sigmoidal SG response curve with a steep transition over a narrow range of chemical stress inducer concentrations. The half-maximal SG response occurred when approximately 38,5 % and 30,6 % eIF2α was phosphorylated upon arsenite and thapsigargin treatment, respectively, defining the level at which cells switched from a SG-Off to a SG-On phase. The steepness of the sigmoidal curve estimated by the Hill function (32) revealed strikingly high coefficients of 14.8 and 18 for arsenite and thapsigargin treatment, respectively, indicating a high degree of cooperativity.

Altogether, the characterization of the core decision module identified a threshold level of p-eIF2α at which cells transition from a SG-Off to SG-On phase. The steep SG response curve revealed the “switch-like” behavior of SG formation when cells were exposed to different chemical stress inducers, with different thresholds for each of the eIF2α-kinases. Of these, PKR emerged as the only eIF2α-kinase whose activation is dampened at higher dsRNA concentrations.

### PKR activation results from its cooperative recruitment to dsRNA

We next focused on characterizing the parameters and reactions involved in the stress-sensing module upstream of eIF2α. To understand PKR activation in more detail, we established an *in vitro* kinase assay using recombinant full-length His-tagged PKR and His-tagged eIF2α, and quantitatively assessed the impact of increasing dsRNA length and molarity on PKR activation. Addition of both 100-bp and 200-bp dsRNAs led to a bell-shaped activation of PKR, which reached its maximum at 10 nM dsRNA, irrespective of the dsRNA length (Fig. 4A, fig. S5A). Under these experimental conditions, i.e. in the absence of phosphatase, full eIF2α phosphorylation was reached at slightly lower dsRNA concentrations (fig. S5B). In contrast to 100-bp and 200-bp-dsRNAs, 40-bp dsRNA was much less potent in activating PKR, leading to a maximum of only 25% of eIF2α being phosphorylated (fig. S5, B and C).

**Fig. 4.**
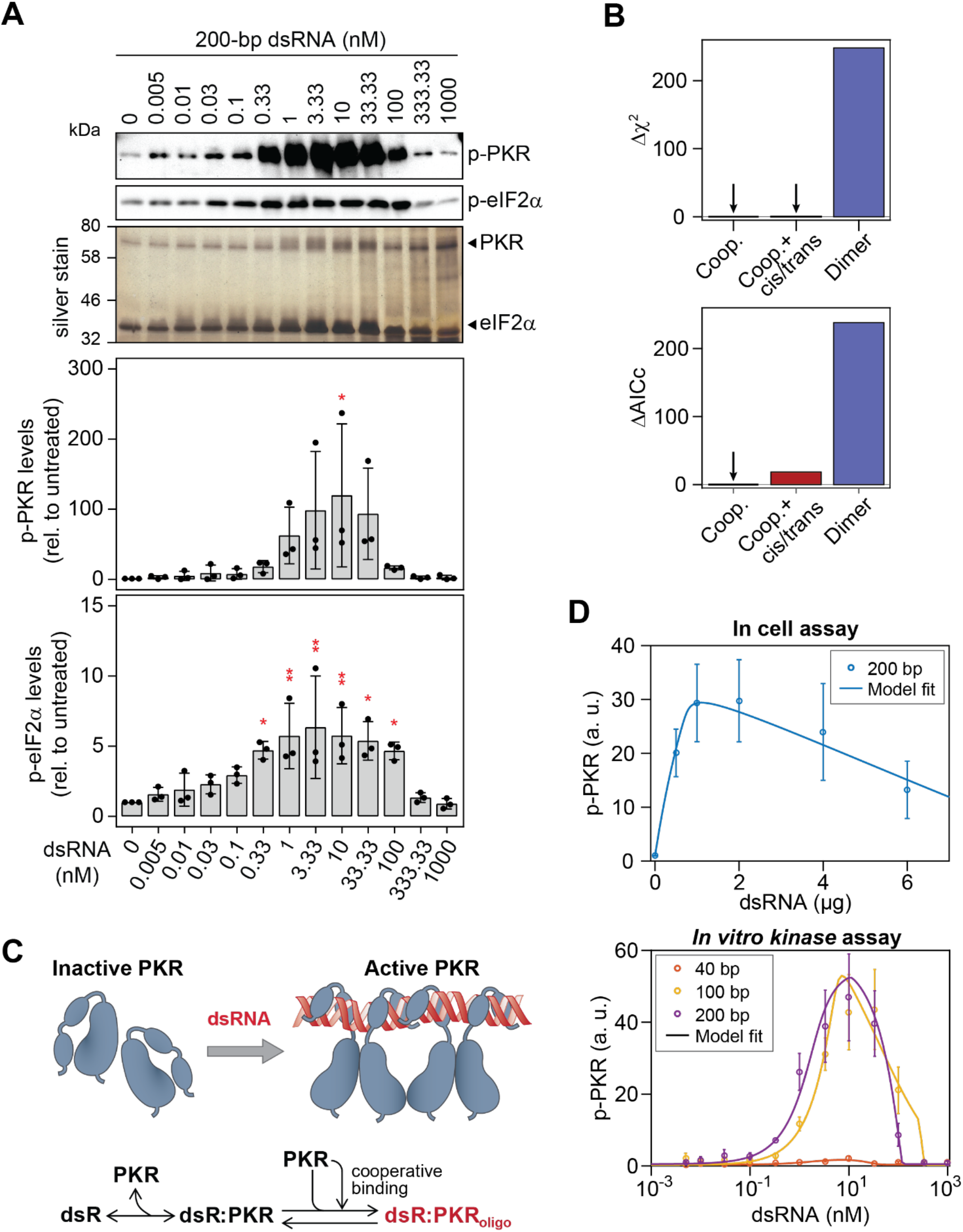
Activation of PKR by dsRNA. (**A**) *In vitro* PKR kinase assay. His-tagged PKR and His-tagged eIF2α were incubated with increasing molarities of 200-bp dsRNA (n=3). The upper panels show representative Western blot analyses of p-PKR and p-eIF2α levels. Silver staining of proteins in the gel served as loading control. Quantifications of mean levels relative to untreated control ± SD are shown in the lower panels. Statistical significance is indicated compared to untreated; *p<0.05, **p<0.01. (**B to D**) Computational prediction of PKR activation by dsRNA. (B) Differences in the chi-square (Δχ^2^) and corrected Akaike information criterion (ΔAICc) to the optimal PKR activation model variant (variant 3, see fig. S6 and S7). The model describing PKR dimerization upon binding to dsRNA was significantly improved by considering PKR cooperative binding to dsRNA (ΔAICc>200). *Cis* and *trans* reactions did not improve the cooperativity model (ΔAICc=18.4). (C) Overview of the optimal model variant. PKR monomers reversibly bind to PKR on dsRNA in a cooperative manner and form active PKR oligomers (dsR:PKRoligo). (D) Best model fits of p-PKR levels to in cell (related to Fig 3B.) and *in vitro* kinase assays (n=500 multi-start optimization runs).

We used this information to formulate a mathematical model of ODEs that describes the sequence of events leading to PKR activation (Supplementary Text section 2, Supplementary Table S1). Starting with a simple model describing PKR dimerization upon binding to dsRNA (fig. S6A and S7, variant 1), several models with stepwise increasing complexity were developed and calibrated with the experimental datasets obtained in cell transfection and *in vitro* kinase assays (fig. S7, Supplementary Text and Supplementary Table S1). Models were fitted at steady state assuming that binding and phosphorylation reactions were fast. The corrected Akaike information criterion (AICc) was used for model selection (Supplementary Text section 2, fig. S6, B and C). Testing the contribution of previously described PKR *cis* (intramolecular) or *trans* (intermolecular) interactions (9, 33, 34) did not significantly improve model fits (Fig. 4B, fig. S7 variant 4 to 5.1). Altogether, the model that most closely recapitulated the experimental results (Fig. 4, C and D, fig. S7 variant 3) revealed three important features of PKR activation: (i) different affinities for PKR to dsRNA of varying lengths, (ii) the formation of PKR oligomers rather than dimers on dsRNA of varying lengths, and (iii) the cooperative recruitment to dsRNA and oligomerization of PKR.

### Quantitative characterization of the GADD34 negative feedback loop

We next explored the parameters involved in the recovery module (Fig. 2A) consisting of the GADD34 negative feedback loop (35). Stress and phosphorylation of eIF2α trigger the translation of the activating transcription factor 4 (ATF4) (36). In turn, ATF4 activates the C/EBP homologous protein (CHOP) and ultimately, in complex with CHOP, the promoter of the ppp1r15a gene encoding GADD34 (36, 37). However, in IFN-competent cells, GADD34 transcription is additionally regulated by the IFN regulatory factor (IRF) 3 and IRF7, downstream of the RNA sensing innate immune pathway (38, 39). We used fluorescence in situ hybridization (FISH) to investigate GADD34 transcriptional regulation in HCV-infected Huh7 cells in the absence and presence of IFN-α treatment (Fig. 5A). This single-cell approach accounted for the low number of infected cells and avoided underestimation in case of bulk measurement. Additionally, cells were co-stained with specific fluorescent probes hybridizing to HCV positive sense single-stranded (+)ss RNA genome and polyadenylated (polyA) mRNA, whereby GADD34 mRNA and HCV infection levels could be directly correlated in SG-negative and SG-positive cells (Fig. 5A, fig. S8, A and B). Fluorescent probes directed against *Bacillus subtilis* dihydrodipicolinate reductase (dapB), a non-eukaryotic transcript, served as control for unspecific binding (neg. ctrl). To assess the contribution of the IRF3/IRF7 pathway in GADD34 transcriptional activation, we circumvented activation of the p-eIF2α-ATF4-CHOP pathway by using Huh7 PKR knockout (PKRKO) cells. As expected, the absence of PKR abolished SG formation in response to HCV infection. Since neither infection nor treatment with IFN-α elicited GADD34 transcription in PKRKO cells (fig. S8, C and D), we concluded that IRF3/IRF7-dependent GADD34 activation is negligible in Huh7 cells. This allowed us to develop a simpler mathematical model that specifically described the stress-induced GADD34 negative feedback loop (Fig. 2A). In naïve Huh7 cells, stress induced by HCV infection in presence of IFN-α resulted in a 2.5-fold increase in the mean number of GADD34 transcripts in SG-positive cells. However, the cell-to-cell variability was more substantial, almost 20-fold (Fig. 5A).

**Fig. 5.**
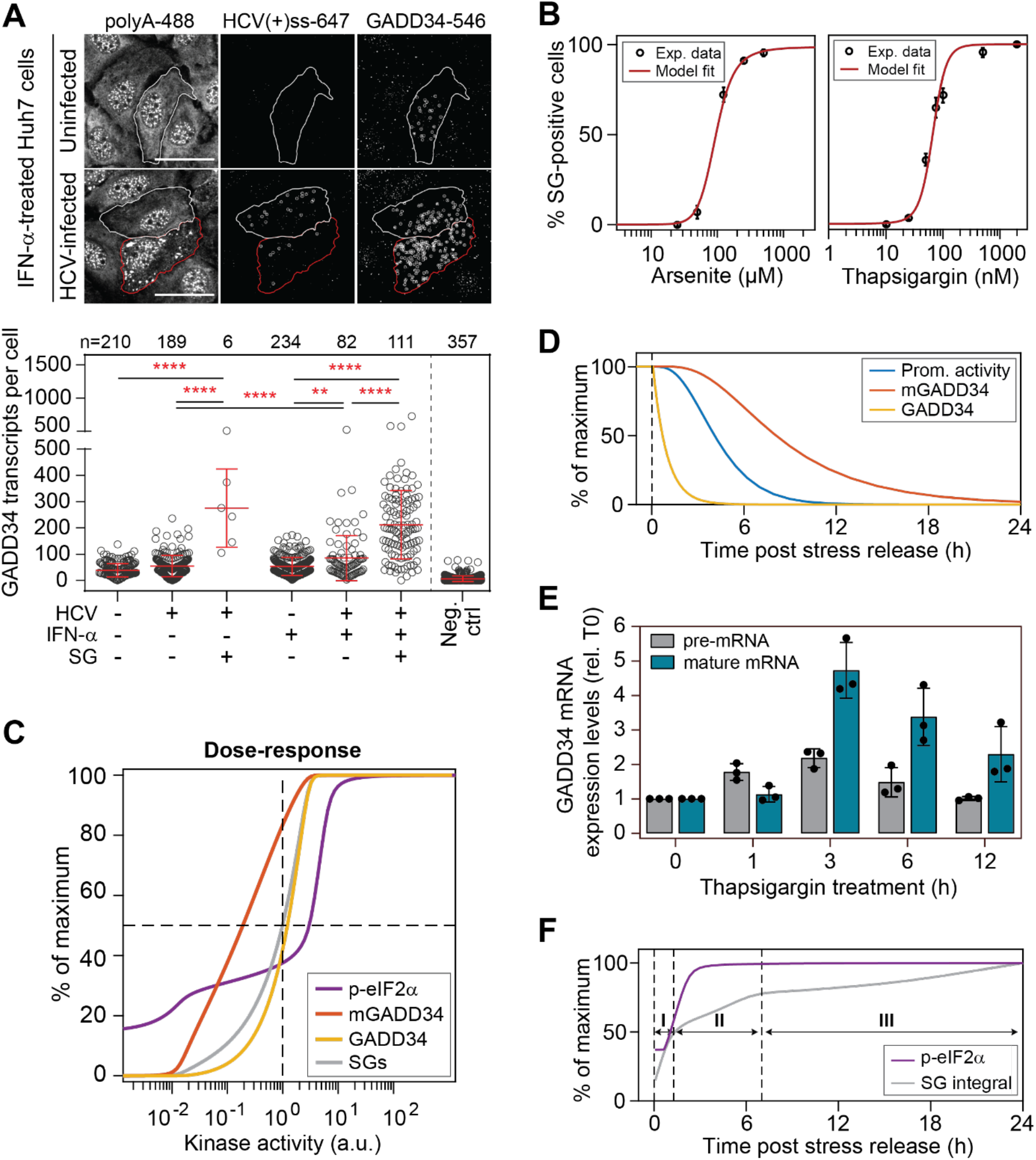
Analysis of GADD34 negative feedback loop. (**A**) Quantification of the mean number of GADD34 transcripts ± SD in Huh7 cells using FISH. The upper panel shows representative still images of uninfected and HCV-infected cells treated with IFN-α for 24 h. HCV (+) ssRNA genomes, GADD34 transcripts and total polyA-tailed mRNAs were simultaneously detected using fluorescent probes. Staining of polyA-tailed mRNAs allowed for the visualization of SGs and cell boundaries. Outlined in red is a representative SG-positive cell. Outlined in white, a representative unstressed cell. White circles indicate single particles detected after background subtraction. Scale bar, 20 μm. The bottom panel shows the scatter plot of GADD34 mean transcript levels ± SD per cell are shown on the bottom. Statistical significance and the number of analyzed cells (n) cells from two independent biological repeats are indicated in the top of the graph; **p<0.001, ****p<0.0001. (**B**) Deterministic model best fits (n=2,500 multi-start optimization runs, see fig. S11 for the results of multi-experiment fitting) to the percentage of SG-positive cells experimentally measured in cells treated with increasing concentrations of arsenite or thapsigargin. (**C**) Computational simulations of dose-response curves for p-eIF2α, GADD34 mRNA and protein and SG-positive cells in the population at steady state. Shown are percentages of maximal values as a function of different kinase activities. The reference kinase activity (100) results in 50% SG-positive cells. Kinase activities below 10^-1^ reflect low to moderate stress. Activities around the reference reflect an intermediate stress. Activities higher than 2 are considered as high stress. (**D**) Model prediction: behavior of the GADD34 negative feedback loop parameters (promoter activity, mRNA and protein) after stress release. Shown are computational simulations of the percentage of their maximum response over time. Estimated decay processes [t1/2, Prom. ≈ 256 min; t1/2,mRNA ≈ 200 min (from (53)); t1/2, GADD34 ≈ 37 min]. (E) Huh7 cells were treated with 2 μM thapsigargin for 12 hours and harvested at the indicated time points. Shown are relative mean expression levels ± SD of GADD34 pre-mRNA and mature mRNA compared to untreated cells (n=3). (F) Model prediction: behavior of the stress response over time, as represented by the levels of p-eIF2α and SG-positive cells after a second one-hour stress pulse applied at different times after stress release. Levels of cell protection against the second stress pulse can be divided in three phases (I to III, indicated by arrows).

Of note, this cell-to-cell variability was significantly smaller for transcriptional induction of PKR by IFN-α (fig. S9, A and B). In addition, GADD34 transcript levels only weakly correlated with the number of HCV genome copies per cell (fig. S9C). This quantitative information was then used to assess the SG response at the population level in a dose-response manner based on GADD34 mRNA levels (Supplement Text section 4.7, fig. S10A).

Time delay is an important parameter of negative feedback loops that exists in numerous mammalian signaling pathways in response to external stimuli (40) and may result in network oscillations (41). For GADD34, this time delay (depicted as a clock in Fig. 2A) includes the time required for *ppp1r15a* promoter activation and upstream open reading frame (uORF)-mediated translation of GADD34 (42). We therefore analyzed p-eIF2α and GADD34 induction kinetics in cells treated with 2 μM thapsigargin, a dose at which all cells formed SGs. Levels of p-eIF2α rapidly increased, reached a maximum after one hour and returned to basal levels approx. four hours post treatment. GADD34 started to visibly increase at 1.5 hours post treatment and reached a maximum at five hours (fig. S11, A and B).

Lastly, we sought to determine the levels at which GADD34 antagonizes the activity of eIF2α-kinases, thereby allowing cells to resume translation and switch from a SG-On to a SG-Off phase. To this end, we ectopically expressed increasing levels of GADD34 using lentiviral transduction and challenged cells with thapsigargin for one hour. At this time point, endogenous GADD34 was not yet detectable (see fig. S11, A and B) and all cells transduced with a control lentivirus formed SGs (fig. S11C). GADD34 expression reduced the level of p-eIF2α and SG formation already at the lowest concentration. We found about 11.6 nM GADD34 protein per cell to antagonize SG formation in 50% of the cells and about 4-fold more was required to antagonize SG formation in all cells (fig. S11D).

Altogether, these results identified Huh7 cells as a unique cell system that allows studying GADD34 stress-induced transcriptional regulation independently of the induced innate immunity pathway. The GADD34 negative feedback loop presents several features that can act as sources of oscillations, e.g. a time-delay caused by the need to transcribe and translate GADD34 mRNA, and non-linear degradation of GADD34. In addition, FISH experiments pointed to an important stochasticity in the expression of GADD34 in this cell system.

### Behavior of the ISR components upon stresses of varying intensities

Based on this quantitative information, we developed an integrative deterministic model describing the ISR. We combined eight model parts, used the measured mean concentrations for eIF2α, PKR and GADD34 protein as well as mRNA as initial values and simultaneously fitted the experimental datasets (Fig. 5B, fig. S10 A to H, Supplementary Text section 4). Since the direct measurement of HCV dsRNA concentrations in infected cells was technically not possible, we extracted starting values from the live-cell imaging experiments assuming that these are proportional to the mean signal intensity of NS5A-mCherry (fig. S10I). Missing mathematical model parameters were estimated by model fitting to experimental data. Model equations, definitions of observables, parameter estimates, and parameter confidence intervals determined by profile likelihood estimation are provided in Supplementary Tables S2–S4.

Using this parametrized ISR model, we studied the behavior of the system over a broad range of stress intensities. To this end, we predicted steady state levels of the key species in response to a one-hour stress pulse of different intensities, here reflected by different eIF2α-kinase activity levels. We defined a kinase activation level of 1 as an arbitrary reference at which 50% of cells form SGs (intermediate stress) and analyzed the response to a range of activation levels, from 100-fold lower (mild stress) to 100-fold higher (acute stress) than the reference. As shown in Figure 5C, the simulations recapitulated the ultrasensitive switch-like behavior of SG formation and GADD34 protein expression within a narrow range of kinase activity around the reference, reaching maximum levels for a 2-fold increase of kinase activity. In contrast, GADD34 mRNA levels were predicted to increase steadily with the kinase activity. Interestingly, p-eIF2α followed a biphasic dose-response curve with a largely hyposensitive phase at mild and moderate stress, during which levels increased only moderately due to the GADD34 negative feedback (between 100-fold less and 2-fold more than the reference), and a hypersensitive phase at higher stress levels (between 2- and 10-fold more than the reference) because of insufficient GADD34. Importantly, at mild to moderate stress levels, the observed shift between the GADD34 mRNA and GADD34 protein dose-response curves suggested that the accumulation of mRNA allows for rapid GADD34 translation in response to moderate p-eIF2α increases, which could be regarded as the “memory” of the previous ISR activation, possibly conferring transient adaptation to stress.

### GADD34 negative feedback loop determines cell adaptation to acute and chronic stress

Adaptation in biological processes refers to the ability of a system to return to or overshoot its initial state after a transient response to environmental changes (43, 44). To investigate this possibility, we simulated the levels of all components involved in the GADD34 negative feedback loop over time following a one-hour stress pulse that leads to a maximal stress response. The promoter activity of *ppp1r15a* and the levels of GADD34 mRNA and protein, all returned to basal levels upon stress release, yet with different kinetics (Fig. 5D). According to its determined half-life, the level of GADD34 protein declined first, followed by the levels of *ppp1r15a* promoter activity, and finally of GADD34 mRNA. We experimentally confirmed this prediction by measuring GADD34 pre-mature mRNA levels, as a proxy for promoter activity, and GADD34 mature mRNA levels in cells treated with thapsigargin over 12 hours. GADD34 pre-mRNA peaked at three hours and returned to basal levels at six hours while mature mRNA remained elevated for up to 12 hours (Fig. 5E). Based on this result, we delineated three time windows after stress release during which cell responsiveness to a second stress pulse increases over time according to the sequential decay of the different GADD34 species (Fig. 5F; phase I, II and III); with phase I corresponding to a short refractory state in which responsiveness to stress is weak due to remaining GADD34 protein, and full responsiveness recovered after 20 hours.

To address the hypothesis of stress adaptation, we simulated consecutive acute stress pulses of varying intensity interspaced by a five-hour recovery period. Since the duration of the kinase activity is known to influence cellular response outcome (45), we modeled different half-lives of kinase activity lasting up to four hours (Fig. 6A, fig. S12). In all scenarios, the second stress pulse resulted in highly attenuated p-eIF2α and SG responses due to the lasting *ppp1r15a* promoter activity and continued presence of high GADD34 mRNA levels (Fig. 6A). This suggests that GADD34 can be directly translated after re-stimulation, leading to rapid dephosphorylation of eIF2α and SG disassembly. To experimentally test this hypothesis we subjected cells to consecutive stress pulses. Because of its rapidly reversible nature, we chose to heat shock cells for one hour at 42°C, and allowed them to recover for five hours at 37°C before challenging them with a second heat pulse. As shown in Figure 6B, the first heat pulse resulted in about 50% of p-eIF2α. Mirroring the appearance of GADD34 protein, the percentage of p-eIF2α decreased over the next five hours of recovery, however, without reaching its basal level. As predicted, the second heat shock did not increase p-eIF2α levels, confirming the lack of cell responsiveness to a second acute stress in this short time window. Altogether, using the calibrated model, we predicted and experimentally validated that the sustained presence of GADD34 mRNA and GADD34 protein limits or prevents the response to the second stress.

**Fig. 6.**
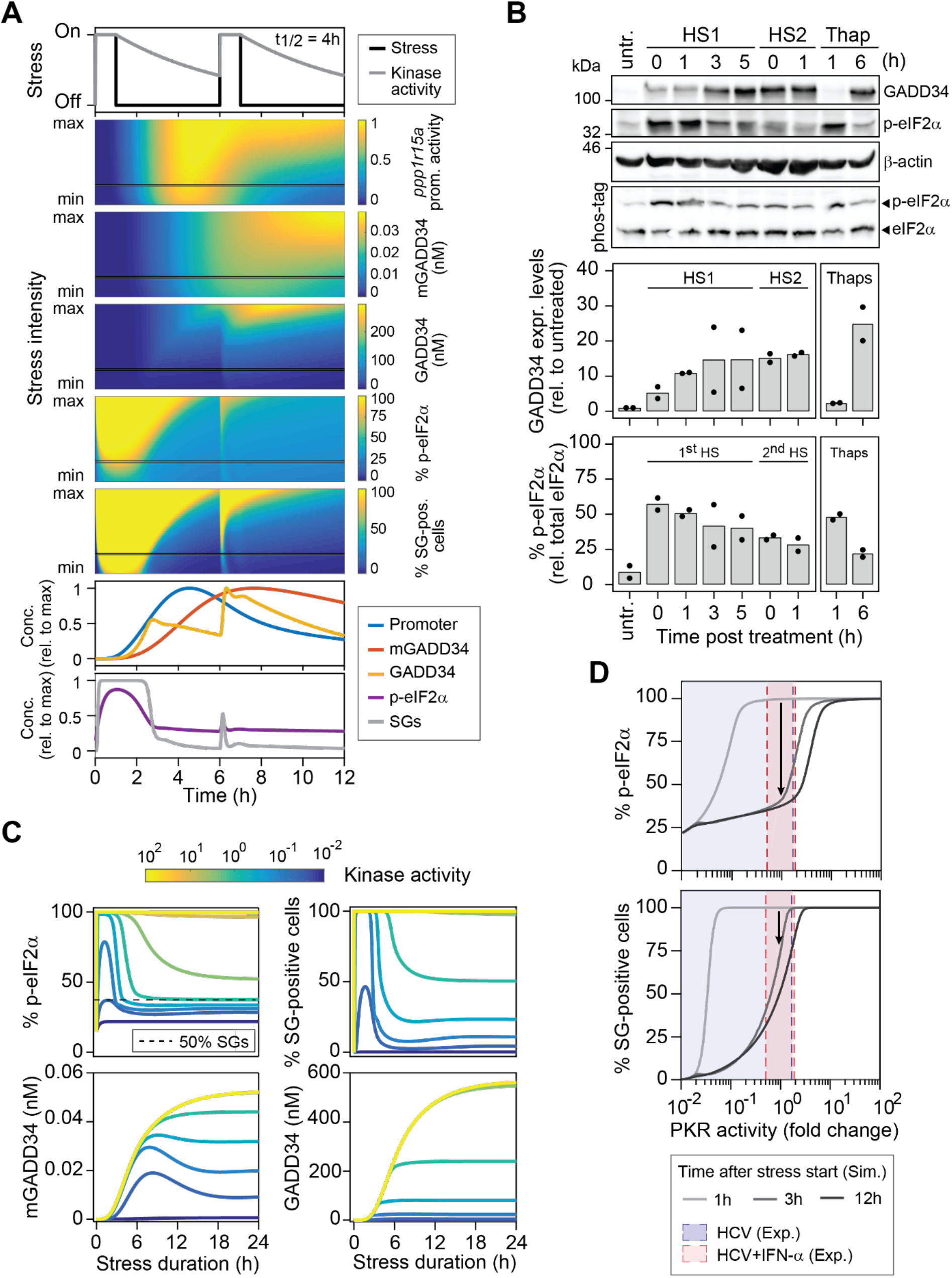
Cell adaptation to repeated and continuous stress. (**A**) Computational simulations of two consecutive one-hour stress pulses interspaced by a five-hour recovery period. Shown is a range of stress kinase activities (stress intensity) varying between 10-fold lower (min) and 10-fold higher (max) than the reference kinase activity leading to 50% SG-positive cells. Color plots show the behavior of *ppp1r15a* promoter activity, concentrations of GADD34 mRNA and protein, percentages of p-eIF2α and SG-positive cells. Graphs at the bottom reflect the behavior of the above mentioned components for one chosen stress intensity (black line, moderate stress). (**B**) Experimental validation of the predictions shown in (A). Huh7 cells were subjected with a first heat-shock (HS1) at 42°C for 1 h and immediately transferred at 37°C for recovery. Cells were harvested at the indicated time points after the first or the second heat-shock (HS2) (n=2). Cells treated with 2 μM thapsigargin for one and six hour served as reference. Shown are representative Western blot and Phos-tag gel analyses. Bottom panels show the quantification of GADD34 expression levels normalized to the loading control and relative to untreated cells as well as the percentage of p-eIF2α relative to total eIF2α. (**C**) Model prediction: behavior of the SG response and GADD34 negative feedback loop components over a 24-hour time period upon continuous stress. The color of the curves reflects different kinase activity levels. (**D**) During the adaptation to a stress stimulus, depending on PKR activity, the expression of GADD34 will result in dose-response curve shifts (hysteresis) in the percentage of p-eIF2α (upper panel) and of SG-positive cells (bottom panel). The shift of the dose-response curve depends on the stress duration and intensity and is reverted after stress relief. Blue and red shared areas indicate 1σ-confidence intervals of estimated PKR activities in HCV and HCV+IFN-α experiments.

Different from arsenite, thapsigargin or heat shock, stress induced by chronic HCV infection is continuous and probably of milder intensity. In addition, by the time cells are treated with IFN-α (48 hours post infection), viral replication has reached a steady state (fig S1A). We thus predicted the behavior of the SG response under continuous stress of different intensities over a period of 24 hours (Fig. 6C). Moderate to intermediate stress resulted in a single burst of eIF2α phosphorylation and SG assembly, and the response declined as a function of GADD34 mRNA and protein induction. In the following, the GADD34 mRNA levels showed a slight decrease after 8 to 10 hours for the milder stress, but remained at a higher plateau compared to the initial level in all cases. This was similar for the level of p-eIF2α and the fraction of SG-positive cells in the population. In fact, these predictions explain our previous results, where we observed that the percentage of HCV-infected cells displaying SGs reached a plateau of 30 to 40% within 12 hours of IFN-α treatment, accompanied by the expression of high GADD34 mRNA levels when measured in bulk (13). Altogether, this result strongly suggests that under infection-induced stress conditions, sustained levels of GADD34 mRNA and protein promote long-term cell adaptation to stress.

To test more accurately if such an adaptation scenario occurs during HCV infection, we simulated the levels of p-eIF2α and SG-positive cells at one, three and 12 hours post stress using the range of PKR activities determined experimentally in untreated and IFN-α-treated Huh7 cells (Fig. 6D). At the low to moderate levels of PKR activation achieved in untreated HCV-infected cells, cells were predicted to rapidly adapt within three hours of the onset of stress, going from 100% SG-positive cells within one hour to less than 25% within three hours, an amount that did not change after 12 hours. In contrast, at intermediate levels of PKR activation in the presence of IFN-α, adaptation was expected to be slower, with approximately 75% of SG-positive cells after 3 hours and 50% after 12 hours before reaching a steady state. Indeed, the analysis of experimental single cell SG response time series supported this hypothesis and revealed the rapid appearance of SGs a in a majority of cells after IFN-α addition. This first SG-On phase was long (approx. 12 hours) followed by shorter phases (see Fig. 1C). This suggested that in our system HCV infection can be considered, amplified by the addition of IFN-α that upregulates PKR, after which cells slowly adapt to stress and exhibit shorter SG phases.

In conclusion, our quantitative deterministic model comprehensively describes the main components of the cellular ISR and temporal behavior of the SG response at steady state. Moreover, our results revealed an unexpected function of GADD34 mRNA levels as a molecular “memory” of the system, which promotes adaptation of cells under mild stress conditions after an acute stress.

### A fine-tuned balance between PKR and dsRNA levels determines SG response dynamics

Although the ISR signaling network could theoretically support sustained oscillations due to the existence of a time delay resulting from GADD34 transcriptional and translational activation (14, 28, 35), the deterministic approach described above was not sufficient to explain the experimentally observed SG response dynamics with recurrent cycles of SG formation. This implied that additional stochastic processes regulate HCV-induced SG response dynamics. We hypothesized that the heterogeneity observed in the SG time series could be influenced by additional parameters, including (i) the inherent stochasticity in GADD34 and PKR expression whose initial protein abundance is low (20, 21, 46, 47), (ii) the inherent stochasticity of the infection process, as the number of particles entering a cell depends on a probabilistic process (48), and (iii) the presence of IFN-α, which induces anti-viral proteins including PKR and represses HCV replication over time (see fig. S1B). To assess these possibilities, we translated the calibrated ODEs into a corresponding stochastic model describing reactions downstream of PKR activation. To this end, kinetic parameters were transformed into reaction propensities. Firstly, to account for the heterogeneity of our cell system as closely as possible, we analyzed the cell-to-cell variability of eIF2α and PKR by single-cell Western blot analysis (49) (fig. S13, A to D). Individual Huh7 cells were loaded into the micro-wells of array slides patterned in a photoactive polyacrylamide gel. After chemical lysis, electrophoresis and UV fixation, proteins were stained with primary and fluorescently-labeled secondary antibodies. The presence of cells in wells was confirmed by staining with GAPDH. For PKR detection, Huh7 PKRKO cells were used as background control. Within the population of Huh7 cells, signal intensities varied by more than 5-fold for eIF2α (fig. S13, A and B), 3-fold for PKR and by more than 6-fold for PKR in cells treated with IFN-α (fig. S13, C and D). We combined these results with the previously determined mean concentrations and thereby determined the log-normal distribution of PKR and eIF2α initial values to be implemented in the model (Supplementary Text section 5). In addition, parameters for the log-normal distribution of NS5A-mCherry intensity were calculated from the live-cell imaging data (fig. S13E, Supplementary Text section 5).

To simulate the ISR in a heterogeneous cell population, concentrations of PKR, eIF2α and infection levels were sampled and used as initial conditions for stochastic simulations of single cell SG response time series (Fig. 7A, bottom graphs). The time series as well as the distribution in SG phase number (Fig. 7B) and stress duration (integral of stress) per day (Fig. 7C) closely reflected the experimental results obtained in live-cell imaging, especially for IFN-α treated cells.

**Fig. 7.**
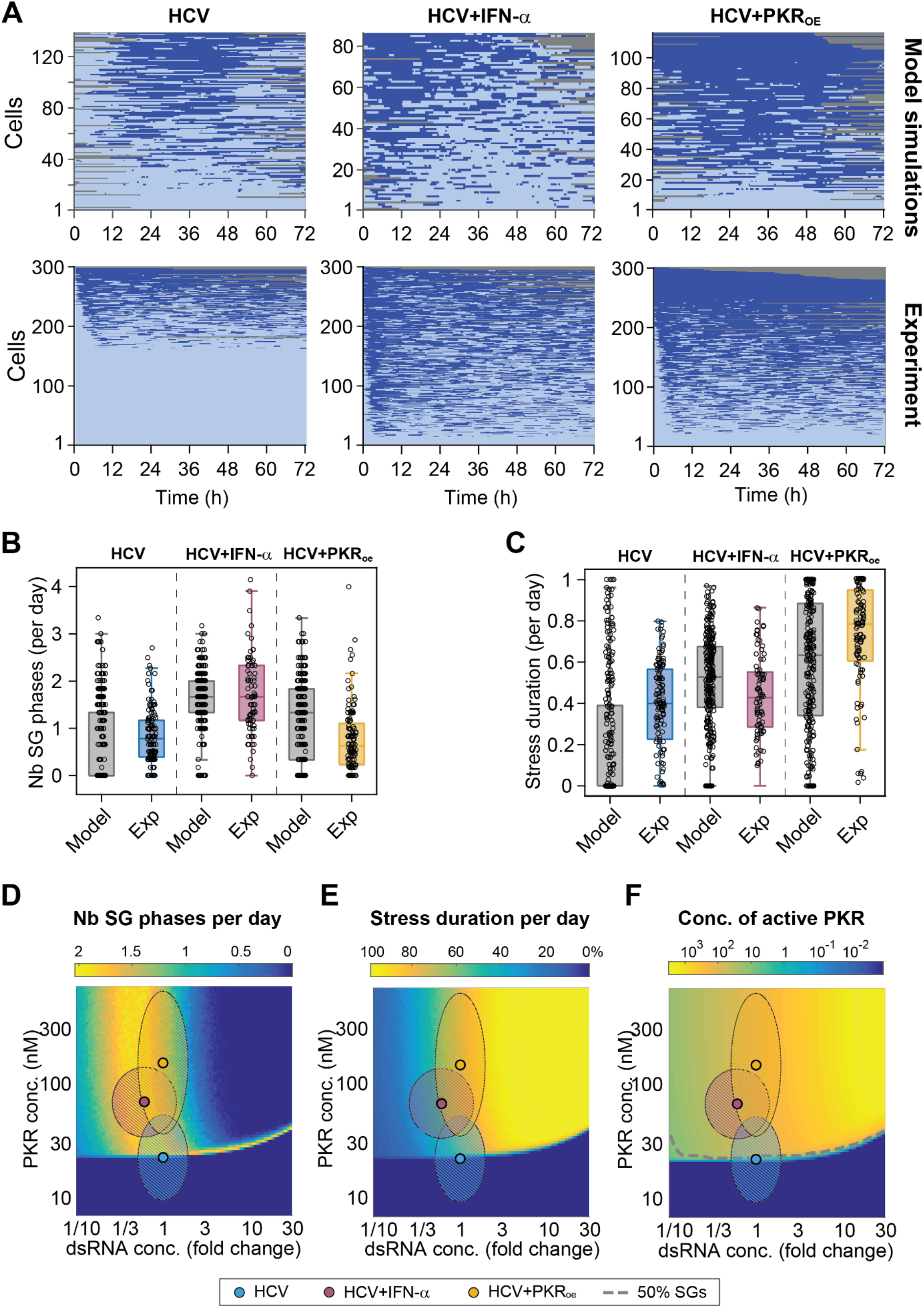
Stochastic mathematical model of the ISR recapitulates HCV-induced SG response dynamics. (**A to C**) Comparison of experimental and computational simulations of three-day single-cell SG response time series using the parameters estimated in HCV-infected cells in presence and absence of IFN-α, and in HCV-infected Huh7 PKR_OE_ cells (bottom panels, n=300) (A). Corresponding number of SG phases per day (B) and stress duration per day (C) were compared. Boxes indicate 25% and 75% percentiles around the median. (D to F) Computational simulations of average SG phases per day (E), stress duration per day (F) and concentration of active PKR (F) for varying PKR and dsRNA concentrations. DsRNA concentrations are expressed as fold changes relative to conditions of the HCV experiment (simulations of 500 single-cell trajectories per combination; hatched ellipsoid areas: 95% confidence intervals for HCV and HCV+IFN-α experiments).

Next, we used the stochastic model to systematically characterize the behavior of the system over a broad range of PKR and dsRNA concentrations. The dsRNA concentration, serving as input for stochastic simulations, was scaled relative to the level analyzed experimentally in HCV-infected cells in absence of IFN-α treatment, corresponding to maximum HCV levels. Color plots in Figure 7 indicate the number of SG phases and stress duration per day (Fig. 7, D and E), as well as the levels of active PKR (Fig. 7F) obtained from simulations of 500 SG response time series per condition. Results revealed that highly fluctuating SG responses occurred only in a narrow range of dsRNA concentrations of about one third of the reference level (Fig. 7D), consistent with the concentration experimentally determined in HCV-infected cells treated with IFN-α (FIG. 7D, pink dot and surrounding area). Interestingly, this prediction also postulated a critical PKR concentration of approximately 25 nM as a threshold below which PKR is not activated in Huh7 cells, regardless of the dsRNA concentration. In agreement with this, we experimentally observed that about half of the Huh7 cells in the population had PKR concentrations below this level, thus accounting for the reduced responsiveness of the population and reduced number of SG phases per day observed in HCV-infected cells. The PKR concentration increases upon treatment with IFN-α, bringing the cells into a state that allows for dynamic SG responses. Furthermore, the simulations revealed that concentrations of dsRNA 3-fold higher than the reference would result in a loss of SG response dynamics, as reflected by the decrease of the number of SG phases per day and an increase in the stress duration, suggesting that SG-On phases would become longer up to reaching a plateau of permanent stress response (Fig. 7E).

Finally, we decided to perturb the system and predict the impact of higher PKR levels on SG response dynamics in HCV-infected cells. Key SG component concentrations in Huh7 cells stably overexpressing PKR (PKR_OE_) (13) were experimentally determined as previously done for naïve Huh7 cells (fig. S14, A and B). Huh7 PKR_OE_ cells expressed mean PKR levels approximately 8.3-fold higher than Huh7 cells and 2.7-fold higher than Huh7 cells treated with IFN-α. Ectopic PKR expression did not affect eIF2α expression levels. In addition, cell-to-cell variability was determined by single-cell Western blot (fig. S14, C and D). Finally, cell-to-cell variability in GADD34 transcripts was analyzed by FISH (fig. S14D). These experimentally determined distributions were used as input for the stochastic model to simulate the SG response when PKR is overexpressed. The resulting time series predicted distinct SG response dynamics with longer SG-On phases that were only rarely interspersed with SG-Off phases (Fig. 7A, model simulations). To experimentally test this prediction, Huh7 PKR_OE_ cells were infected with HCVTCP for 48 hours and SG formation was monitored using live-cell imaging for an additional 72 hours (Fig. 7A, Exp. HCV + PKR_OE_, Movie S3). The analysis of the experimental single-cell SG response time series showed that, as predicted by the model, cells displayed on average longer SG-On phases and a higher stress duration per day (Fig. 7, C and E, HCV + PKR_OE_). Consistently, this was paralleled by a reduced number of SG phases per day (Fig. 7, B and D, HCV + PKR_OE_).

Collectively, our results revealed that the dynamic SG response to HCV infection, and generally to dsRNA, consists of repetitive stochastic transitions between On- and Off-states that are regulated by PKR and dsRNA concentrations in individual cells. The stochastic version of the deterministic model that was calibrated using an extensive experimental dataset could successfully predict characteristics of single cell SG responses in heterogeneous cell populations, and was independently validated for the case of PKR overexpression. In addition, our modeling approach points towards elevated GADD34 mRNA expression levels that act as a memory of previous ISR activation and enable long-term adaptation of chronically infected cells to the continuous presence of virus.

## Discussion

The ISR is a pivotal process that allows cells to adapt to environmental changes by rapidly repressing host cell translation, thereby preventing damage to nascent polypeptides and reallocating resources to restore homeostasis. Its regulation has been the subject of many studies involving mathematical modeling (39, 50–55). However, these studies have mainly focused on the unfolded protein response network via PERK-eIF2α signaling. Possible periodic oscillations of the stress response components were predicted in only two of the studies under certain parameter conditions that were not determined quantitatively (39, 55). While these studies provided valuable information on the possible behavior of the stress response over time, they could not explain the peculiar SG fluctuations observed in response to chronic HCV infection.

By combining long-term live-cell imaging, population and single-cell analyses, and ODE modeling, we here provide a detailed quantitative insight into the individual regulatory steps of the ISR, including SG formation. In particular, we captured the dynamics and role of the different components of the stress response to HCV infection. In the quantitative model we developed, eIF2α is the central reaction component of the system whose phosphorylation mediated by the different eIF2α-kinases can be reversed by PP1 in complex with GADD34 in a negative feedback manner. Although the topology of this stress-kinase-eIF2α-GADD34 signaling network displays the hallmarks of a biological oscillator (14) and despite our previous intuitive description of HCV-induced SG On- and Off-phases as “oscillations”, the Fourier transform analysis of hundreds of single-cell SG response time series suggested that the observed SG phases are rather bursts controlled by a stochastic process with memory. While our mathematical analysis focused on finding distinct oscillation frequencies in a heterogeneous cell population, we cannot exclude the existence of individual cells whose parameters may lie in the range that permits oscillation. An algorithm developed by Phillips and colleagues represents an attractive alternative that allows finding single oscillators of varying frequencies in a cell population, thereby differentiating between signal fluctuations and noise (56). However, the binary SG response time series generated by our analysis pipeline are not suitable for this type of analysis, which requires a continuous signal as well as a signal amplitude as input information.

Dose-response experiments with arsenite and thapsigargin treatments provided evidence for the remarkable switch-like behavior of SG formation. This was reflected by a steep sigmoidal response curve and high Hill coefficients and suggested that SG assembly is a highly cooperative process, consistent with current knowledge about SG assembly by liquid-liquid phase separation (57). Thus, we determined a threshold level of p-eIF2α of 38% (arsenite) and 30% (thapsigargin), which is critical for Huh7 cells to switch to an SG-On phase. These values depend on the concentration of the respective kinase, and are in the range of what other studies described (58–60).

Some aspects of the mechanism of PKR activation are still unclear, including how monomers are recruited to dsRNA. By combing the quantitative information on the stoichiometry of PKR activation obtained *in vitro* and mathematical modelling, we observed cooperative binding of PKR molecules with 100-bp and 200-bp dsRNA, unlike others have seen with shorter dsRNAs (10). Regardless of dsRNA length, maximal PKR activation and eIF2α phosphorylation were achieved at 10 nM dsRNA, a concentration lower than previously reported (10, 61), which might be explained by lower concentrations of PKR and salt in our assay. Unlike what is observed at higher concentrations (62), our results did not provide evidence that at physiological level PKR can be activated in the absence of dsRNA. Due to the absence of time-resolved measurements, the previously hypothesized contribution of *cis* (intramolecular) and *trans* (intermolecular) interactions in PKR activation was not evident from our dataset (33, 34, 62–64).

An important finding of our deterministic model was that the ISR allows different levels of adaptation. Adaptive responses have been reported for various biological systems, e.g. during osmotic shock in yeast (65), perturbations of calcium homeostasis in mammalian cells (66), and chemotaxis in E. coli (67). Adaptation is defined, from a biochemical point of view, as the ability of a system to respond to a stimulus and return to the pre-stimulus state or to a different steady state. In this manner, cells can maintain homeostasis, especially in the presence of persistent perturbations (68). The topology of our ISR model exhibits strong similarity with other theoretical minimal networks, e.g. the negative feedback loop with a buffer node (43), previously described to allow biochemical adaptation. Although theoretically possible, our deterministic model of the ISR revealed the absence of oscillations (for p-eIF2α level and SG response) within the experimentally determined range of the different parameters. Using repeated heat pulses, we confirmed that after a first acute stress, long-lasting GADD34 mRNA levels protect cells from responding a second stress pulse. For a continuous stress of intermediate intensity, as triggered by HCV infection in presence of IFN-α, our analyses confirmed that adaptation occurs within 12 hours with GADD34 mRNA levels remaining a high steady state level over time (13). Based on this result, we propose that GADD34 transcriptional activation and mRNA serve as a “memory” of the system, allowing for rapid GADD34 translation and limiting the SG phase duration in the case of both acute and continuous stress. Our results are in agreement with several studies that have either addressed the impact of repeated stress pulses theoretically (50, 51), or explored the response of chronic chemical stress experimentally (52, 54, 69).

Finally, our work highlights the importance of cell-to-cell variability and stochasticity in shaping cellular responses to stress. Translating the calibrated model into a corresponding stochastic model, by taking into account random bursts of transcription and cell-to-cell variability, allowed us to faithfully model HCV-induced SG response fluctuations observed in long-term live-cell time lapses. To our surprise, the results indicated that recurrent SG phases occur only in a relative narrow range of PKR and dsRNA concentrations. By combining the information obtained from the calibrated deterministic and stochastic models, we propose a scenario in which the stress response to HCV, in presence of IFN-α, infection reaches a steady state level within 12 hours, determined by the concentration of active PKR and dsRNA at the single cell level, allowing for an adaptation to this type of viral chronic stress. In this system, recurring SG-On and SG-Off phases are caused by stochastic events that perturb the steady state level of eIF2α phosphorylation and evoke bursts of GADD34 transcription.

The fact that eIF2α-kinases are interchangeable in our quantitative hybrid model of the ISR opens up several possibilities for future investigations. For instance, it will be interesting to address the impact of other chronic treatments with low doses chemical stress inducers. In addition, expanding the model to quantitative data sets from IFN-competent cell types and other RNA viruses (39) would further shed light on the role of the IFN-mediated regulation of GADD34 expression in the ISR to virus infection. Altogether, the hybrid model developed here will help to further understand the detailed mechanisms and kinetics underlying cellular adaptation to stress and stress recovery.

## Materials and Methods

### Cell lines and cell culture reagents

All cell lines were cultured in Dulbeccos’s modified Eagle’s medium (DMEM) supplemented with 2 mM L-glutamine, 1x non-essential amino acids, 100 U/ml penicillin, 100 μg/ml streptomycin (all from GIBCO, Life Technologies) and 10 % fetal calf serum (Capricorn), hereafter referred to as DMEM complete. HEK 293T cells were used for lentivirus particle production. Huh7 PKRKO clones were described elsewhere (70). Huh7 YFP-TIA1 Neo cells (13) were supplemented with 1 mg/ml G418 (Invitrogen, Life Technologies). Huh7 YFP-TIA1 Neo PKR Blr cells and Huh7 PKR Blr cells (PKR_OE_)(13) were additionally supplemented with 5 μg/ml blasticidine (Life Technologies). Huh7 GADD34 Puro cells (13) were supplemented with 3 μg/ml puromycin (Sigma-Aldrich). Huh7.5 cells (71) were used for production of HCV (Jc1) virus stock. Huh7.5 [CE1][E2p7NS2] Blr cells (72) were supplemented with 5 μg/ml blasticidine and used for the production of trans-complemented HCV (HCVTCP) particles. For live-cell microscopy, cells were maintained in phenol red-free DMEM containing 1 mM HEPES and supplemented with penicillin/streptomycin and 10% fetal calf serum (microscopy medium).

### Plasmids

A plasmid encoding HCV subgenomic replicon harboring *Renilla* luciferase (RLuc) gene for monitoring replication levels and mCherry fused to HCV NS5A to allow identification of infected cells in live cell imaging was generated by PmeI and AgeI digest of pFKI389neoNS3-3’dg_JFH-1_NS5A-aa383_mCherry K1402Q plasmid (13). RLuc coding sequence flanked by AgeI and PmeI restriction sites was amplified from the vector pFK i389 JcR2a dg (73) by PCR using Phusion High-Fidelity DNA polymerase (New England Biolabs) using the following primers: AgeI-RLuc_Forward: 5′-CGG AAC CGG TGA GTA CAC - 3′and RLuc-PmeI_Reverse: 5’ - AGG CGT TTA AAC TTA TTC ATT TTT GAG AAC TCG - 3′. This replicon (pFKI389RLuc_NS3-3’dg_JFH-1_NS5A-aa383_mCherry K1402Q, designated ‘HCV RLuc-mCherry replicon’) was used to generate HCV_TCP_ particles for live-cell imaging.

To generate plasmids used for bacterial expression of recombinant His-tagged proteins, sequences of PKR (*EIF2AK2*) and eIF2α (*EIF2S1*) were amplified by PCR using the Phusion Flash High-Fidelity PCR Master Mix (Thermo Scientific), pWPI PKR Blr or pET-MCN eIF2α (kindly provided by G. Stoecklin, Mannheim) as template and primers introducing sequences C- and N-terminally overlapping with the pET-His 1a vector (PKR_For: 5’ – TTA TTT TCA GGG CGC CAT GGC TGG TGA TCT TTC AG – 3’; PKR_Rev: 5’ – CGA ATT CGG ATC CGG TAC CCT AAC ATG TGT GTC GTT C – 3’; eIF2α_For: 5’ – TTA TTT TCA GGG CGC CAT GCC GGG TCT AAG TTG TAG – 3’; eIF2α_Rev: 5’ – CGA ATT CGG ATC CGG TAC CTT AAT CTT CAG CTT TGG CTT C – 3’). Full pET-His 1a vector (kindly provided by I. Sinning, Heidelberg) was amplified by PCR (pET-His_1a_For: 5’ – GGT ACC GGA TCC GAA TTC G – 3’ and pET-His_1a_Rev: 5’ GGC GCC CTG AAA ATA AAG – 3’). The following PCR conditions were employed: 98 °C for 10 sec and 30 cycles of 98 °C for 1 sec, 55 °C for 5 sec and 72 °C for varying periods (90 sec for PKR, 15 sec for eIF2α, 20 sec for pET-His 1a). A final extension at 72 °C for 60 sec was performed. Template plasmid was digested using DpnI FastDigest (Thermo Fisher) for 1 h at 37°C. PCR products were purified from agarose gels using the Nucleospin Gel and PCR Clean-Up kit (Macherey-Nagel). *EIF2AK2* and *EIF2S1* sequences were inserted into pET-His 1a vector using the Gibson Assembly Cloning Kit (New England Biolabs) as described by the manufacturer.

pWPI GADD34-(G4S)4-eGFP Puro used for lysate calibration for absolute quantification of GADD34 molecules per cell was generated by amplifying ppp1r15a sequence from its cDNA clone (13) by PCR introducing XhoI and HindIII restriction sites flanking the PCR product using the following primers: XhoI_GADD34_For: 5’ – AAC TTC CTC GAG ATG GCC CCA GGC CAA GCA CCC CAT C – 3’ and HindIII_GADD34_Rev: 5’ – GGA TCG AAG CTT GCC ACG CCT CCC ACT GAG GTC CAG G – 3’. First, the PCR product and vector pTurboGFP-N (Evrogen) were digested with XhoI and HindIII purified and ligated to generate pGADD34-TurboGFP. Second, the sequence of a (G4S)4-linker flanked by HindIII and AgeI restriction sites was generated by hybridizing 100 μM of each of the following oligonucleotide (HindIII_(G4S)4_AgeI_For: 5’ – AGC TTG GTG GAG GCG GGT CTG GGG GCG GAG GTT CAG GCG GGG GTG GTT CCG GTG GCG GTG GCT CGG GA – 3’ and AgeI_(G4S)4_HindIII_Rev: 5’ – CCG GTC CCG AGC CAC CGC CAC CGG AAC CAC CCC CGC CTG AAC CTC CGC CCC CAG ACC CGC CTC CAC CA – 3’) in annealing buffer (25 mM Hepes pH7.4, 50 mM NaCl) at 95 °C for 5 min followed by a gradual decrease in temperature to 25 °C at 5 °C per minute. Hybridized oligonucleotides were phosphorylated with 5 U T4 polynucleotide kinase in T4 Ligation buffer (New England Biolabs) for 30 min at 37 °C. Finally, pGADD34-TurboGFP was digested by HindIII and AgeI and pGADD34-(G4S)4-TurboGFP was generated by ligation with the hybridized oligonucleotide. Of note, TurboGFP was exchanged for eGFP by amplifying the eGFP sequence by PCR from peGFP-N1 (Clontech) using the following primers to introduce AgeI and NotI restriction sites: AgeI_eGFP_For: 5’ – TTA TTA GAC CGG TCA TGG TGA GCA AGG GCG AGG A – 3’ and NotI_eGFP_Rev: 5’ – TAA TTG CGG CCG CTT ACT TGT ACA GCT CGT CCA – 3’. PCR product and vector were digested with AgeI and NotI and the digested vector was purified by gel extraction. pGADD34-(G4S)4-eGFP was generated by ligation. GADD34-(G4S)4-eGFP sequence was finally cloned into a lentiviral vector pWPI Puro. To this end, pGADD34-(G4S)4-eGFP was digested by XhoI and NotI. Sticky ends were blunted using 10 U of DNA Polymerase I Large (Klenow) Fragment (New England Biolabs). pWPI Puro was digested by PmeI and dephosphorylated using 20 U of calf alkaline phosphatase (New England Biolabs) for 1 h at 37°C. The lentiviral vector pWPI GADD34-(G4S)4-eGFP Puro was generated by ligation.

### *In vitro* transcription

*In vitro* transcription reaction was performed as described earlier (13). Ten micrograms of plasmid template were linearized by restriction with MluI (New England Biolabs) and purified using the Nucleospin Gel and PCR Clean-Up kit (Macherey-Nagel). *In vitro* transcription reaction was carried out in a final volume of 100 μl transcription mix containing 80 mM HEPES (pH 7.5), 12 mM MgCl2, 2 mM spermidine, 40 mM dithiothreitol, 3.125 mM of each nucleoside triphosphate (Roche), 100 U of RNasin ribonuclease inhibitor (Promega), and 80 U T7 RNA polymerase (Promega). After 2 h incubation at 37°C, 40 U of T7 RNA polymerase were added to the reaction mix and incubated for an additional 2 h at 37°C. Transcription was terminated by addition of 10 U RQ1 RNase-free DNase (Promega) and incubation at 37°C for 30 min. RNA was extracted with acidic phenol and chloroform, precipitated with isopropanol, and dissolved in RNase-free water. RNA integrity was determined by using non-denaturing agarose gel electrophoresis and RNA concentration was determined by measuring absorption at 260 nm.

### Analysis of HCV replication kinetics

To characterize replication kinetics of HCV RLuc-mCherry replicon, *in vitro* transcripts were introduced into Huh7.5 cells by electroporation. To this end, single-cell suspensions were generated by trypsinization, washed twice with phosphate-buffered saline (PBS) and resuspended in cytomix solution (74) freshly supplemented with 5 mM glutathion and 2 mM ATP to a final concentration of 1.5×10^7^ cells per ml. 400 μl of cell suspension was mixed with 10 μg of *in vitro* transcript, transferred in a 0.4 cm electroporation cuvette (Bio-Rad) and pulsed at 975 μF and 270 V using a Gene Pulser system (Bio-Rad). Electroporated cells were immediately resuspended in 15 ml DMEM complete and seeded into one 6-well plate in duplicates. At 4 h, 24 h, 48 h, 72 h and 96 h, cells were washed once with PBS and lysed using 250 μl of ice-cold Luciferase Lysis Buffer (25 mM Glycyl-Glycine pH 7.8, 15 mM MgSO_4_, 15 mM K_2_PO_4_, 4 mM EGTA, 10 % (v/v) glycerol, 0.1 % Triton X-100) freshly supplemented with 1 mM dithiothreitol. Plates were stored at −80°C. For Renilla luciferase activity measurement, 20 μl lysates thawn on ice were mixed with 100 μl Luciferase Assay Buffer (25 mM Glycyl-Glycine pH 7.8, 15 mM MgSO_4_, 15 mM K_2_PO_4_, 4 mM EGTA). Lysates were measured in duplicates per well using a tube luminometer (Berthold Technologies). Relative light unit (RLU) values from the 4 h time point post electroporation served as input control for the normalization of RNA levels.

### Virus production and titration

For production of RLuc-mCherry HCV_TCP_ used to observe SG response dynamics by live-cell imaging, Huh7.5 [CE1][E2p7NS2] Blr cells were electroporated with 10 μg of HCV RLuc-mCherry replicon *in vitro* transcript as described above, immediately resuspended into 6.5 ml DMEM complete and seeded into a 10 cm-tissue culture dish. Virus supernatants were collected at 24, 48, 72 and 96 h post electroporation and filtered using filter units (Merck, 0.45 μm membrane pore size). Infectious titers of virus stocks were determined by limiting dilution assay (TCID_50_/ml) using Huh7 cells as described in (75).

Full-length HCV Jc1 was employed to analyze levels of various proteins and transcripts per cell during virus infection. To generate a virus stock, Huh7.5 cells were electroporated with *in vitro* transcripts from pFK-J6/C3 (76) and seeded as described above. Cell supernatants were collected and filtered as described above. Virus particles were concentrated by precipitation using 8 % (w/v) polyethylene glycol-8000 in PBS for 72 h at 4 °C followed by 2 h centrifugation at 8,000 x g. The virus pellet was resuspended in DMEM complete and stored at −80 °C. Infectious titers were determined by limiting dilution assay as described in (75).

### Analysis of HCV_TCP_ replication kinetics upon IFN-α treatment

To measure replication levels of HCV RLuc-mCherry replicon in response to IFN-α treatment, 3×104 Huh7 cells were seeded in 12-well plates and infected with HCV_TCP_ at a MOI of 2 TCID_50_ per cell in duplicate wells for 48 h. Medium was replaced with 2 ml of DMEM complete with and without 100 IU/ml IFN-α (PBL International). Cells were lysed at 24 h, 48 h or 72 h after treatment. *Renilla* luciferase activity analyzed as described above. Relative light unit (RLU) values were normalized to the time point of IFN-α addition (T0).

### Long-term live-cell imaging of HCV-infected cells and microscope equipment

1.8 × 10^4^ Huh7 YFP-TIA1 Neo cells with or without stable overexpression of PKR were seeded 24 h prior to infection in 12-well plates with glass bottom (thickness 0.16-0.19 mm) (Cellvis). Cells were infected with HCV_TCP_ at a MOI of 1.5 TCID_50_ per cell. Medium was replenished 24 h after infection. 48 h post infection, culture medium was replenished with phenol red-free microscopy medium supplemented with 100 IU/ml IFN-α (PBL International) and transferred to the heating chamber of the microscope (Okolab). Image acquisition was performed using a Nikon Eclipse Ti2/Andor Revolution CSU-W1 spinning disc confocal microscope equipped with a 20x air objective CFI Plan Apo lambda (NA=0.75), Nikon Perfect Focus System (PFS), Andor lasers 514 nm (40 mW) and 561 nm (50 mW), triple line dichroic beamsplitter 445/514/561, emission filters for YFP (540/30) and mCherry (600/50), Nikon motorized stage with linear encoder, EM CCD Camera iXon DU-888 - 13 x 13 μm pixel size (Andor), NIS Elements AR software (Nikon). Images (signals of YFP-TIA1 and NS5A-mCherry) were acquired in 15-min intervals for 72 h, starting four to five hours after treatment with IFN-α. Typically 15-30 fields of view were manually selected using the NS5A-mCherry signal and acquired simultaneously.

### Image analysis of SG single cell time series

Images were exported as one 16-bit hyperstack (TIF format) per field of view using NIS Elements AR software. Using the ilastik (Linux v1.3) (27) Image Conversion Tool, the images were further converted into 8-bit .h5 format. ilastik Pixel Classification Tool was used for automatic detection of single cells, stress granules and virus protein. Random frames were used for software training by manually creating separate labels for pixels representing nuclei, SGs, background pixels (all in the YFP-TIA1 channel) and virus protein (NS5A-mCherry channel). Using the Segmentation Tool, segmentation masks were created for each label. Single cell tracks were generated with the Ilastik Tracking Toolkit using the nuclei segmentation masks. To assign density and count as well as virus protein levels to individual cell tracks at any given point in time, we utilized a Voronoi partitioning of each timeframe based on the nuclei segmentation center of mass. For each cell track and frame, the respective Voronoi cells were overlaid with SG and infection marker masks to generate time-resolved single-cell trajectories of total number of SG pixels, SGs and NS5A-mCherry pixels. To exclude faulty cell tracks and SG detection, each track was manually curated using a Python/Kivy-based graphical interphase iterating every timeframe of interface long tracks (min. 48 h length) and comparing time-lapse data with assigned center of mass of tracks and detected SG pixel number. Only tracks with correctly assigned presence of SGs (SG pixel threshold >15) with maximum aberrations of 1 frame false positive/negative SG detection were accepted. In a further downstream data filter, all stress events/stress disruption events of 1 frame length were smoothed out.

### Estimation of Huh7 cell volume

10^5^ Huh7 YFP-TIA1 were seeded in a 35-mm dish with 20 mm cover glass #1.5 (MatTek). Confocal Z-stacks (200 nm Z-spacing) of whole cells were acquired using the YFP-TIA1 signal (514 nm excitation) on a Nikon Ti Eclipse using a Apo TIRF 60x oil (NA 1.49) objective (Nikon) with a digital image using the Orca Flash 4 v2 camera - 6,5 x 6,5 μm pixel size - (Hamamatsu). Substacks with 1 μm Z-spacing including the whole cell volume were chosen and manually circled to measure area per stack. Total cell volume was calculated by multiplication of total area volume with 1 μm stack thickness.

### Western blot analysis

For Western blot analysis, cells were detached by trypsinization. Cell pellets were lysed with ice-cold protein lysis buffer (50 mM Tris-HCl pH 7.3, 150 mM NaCl, 1 % Triton-X100) supplemented with EDTA-free protease inhibitor cocktail (c0mplete Mini EDTA-free, Roche) and phosphate inhibitors (60 mM β-glycerophosphate, 15 mM 4-nitrophenylphosphate, 1 mM sodium orthovanadate, 1 mM sodium fluoride) on ice for 30 min. Cell debris were pelleted by centrifugation for 30 min at 13,000 rpm at 4°C. Total protein concentration was determined by colorimetric measurement using Protein Assay (Bio-Rad). Bovine serum albumin (Thermo Fisher Scientific) dilutions served as protein standard for calibration. Equal amounts of total protein (20 - 100 μg, depending on experiment) were resuspended in 1x Laemmli sample buffer (62.5 mM Tris-HCl pH 6.8, 10 % glycerol, 1.5 % SDS, 1.5 %β-mercaptoethanol, 0.01 % bromophenol blue), denatured for 5 min at 95°C, separated by SDS-PAGE and transferred to a PVDF membrane (Millipore). Membranes were blocked by incubation with Tris-buffered saline (TBS) containing 0.1 % Tween 20 and 5 % (w/v) skim milk powder or 5 % (w/v) BSA (Roth) depending on the antibody requirements for 1 h,. Immunostaining was performed in the corresponding buffer supplemented with the respective primary and secondary antibodies. Proteins were detected using Western Lightning ECL Plus (Perkin Elmer) according to the instructions of the manufacturer. Chemiluminescence signal was detected using Advance ECL Chemocam Imager (Intas Science Imaging). Band intensities were quantified using the LabImage 1D Software (v 4.1; Intas Science Imaging).

The following primary antibodies were used: rabbit polyclonal anti-phospho-PKR (T446) (Abcam; BSA; 1:500), rabbit polyclonal anti-rabbit-phospho-eIF2α (S51) (Cell signaling; BSA; 1:500), rabbit polyclonal anti-PKR (K-17) (Santa Cruz; milk; 1:1,000) for GST-PKR titration experiment, rabbit polyclonal anti-PKR (Proteintech; BSA; 1:1,000) for detection in other experiments, rabbit polyclonal anti-eIF2α (Cell signaling; BSA; 1:1,000), rabbit polyclonal anti-GADD34 (Proteintech; milk; 1:1,000), rabbit polyclonal anti-cyclin D1 (H-295) (Santa Cruz; BSA; 1:1,000), mouse monoclonal anti-β-actin (Sigma-Aldrich; milk; 1:5,000), mouse monoclonal anti-GAPDH (G-9) (Santa Cruz; milk; 1:10,000) and mouse monoclonal anti-GFP (Clontech; milk; 1:2,000).

### Analysis of protein half-lives

Huh7 cells or Huh7 GADD34 Puro cells were incubated with 100 μg/ml cycloheximide (Sigma-Aldrich) and harvested at different times after treatment. Protein expression levels were measured by Western blot and band intensities quantified using LabImage1D as described above. Protein half-life was determined by normalization to loading control (GAPDH or β-actin) and compared to levels in untreated cells. Cyclin D was used as control for short-lived protein.

### Absolute quantification of eIF2α and PKR protein in cell lysates

2 x 10^6^ Huh7 YFP-TIA1 Neo with or without stable overexpression of PKR were seeded on 15 cm cell culture dishes. One day after seeding, cells were incubated with DMEM complete supplemented with 100 IU/ml IFN-α or left untreated. 24 h after treatment, cells were detached by trypsinization, counted by FACS-sorting and lysed in protein lysis buffer as described above. To ensure thorough lysis, lysates were subjected to three freeze/thaw cycles at −80 °C before centrifugation. Total protein concentration was measured as described in the previous paragraph. Lysates (n=8) containing 20 μg of total protein were spiked with different amounts of recombinant GST-tagged eIF2α (Abnova; 5/10/15/22/33/50/75 ng), recombinant GST-tagged PKR kinase domain (Abcam; 1/2/3/4.4/6.7/10/12.5 ng) or protein lysis buffer. Samples were supplemented with Laemmli sample buffer, boiled for 5 min at 95 °C and subjected to SDS-PAGE and Western blotting as described above. Membranes were incubated with antibodies targeting epitopes present in endogenous and recombinant protein as described above. Band intensities were quantified as described above and recombinant protein titration intensities were used as standard curves to quantify average endogenous protein molecule numbers per cell.

### Absolute quantification of GADD34 protein in cell lysates

Huh7 cells were reverse transduced with lentivirus of pWPI GADD34-eGFP Puro for 30 h as previously described. To determine GADD34-eGFP molecule number per cell, lysate of 1.75 x 10^5^ cells (ca. 100 μg total protein as determined by protein assay) was analyzed in triplicates by Western blotting. A dilution series (0.5/1/2.5/5/10 ng) of recombinant eGFP protein was used to generate a standard curve. Whole-lane signal of membranes probed with eGFP-specific antibody was used to determine the total molecule number of GADD34-eGFP expressed in the standard lysate, including degradation products. The average molecule number per cell was used to calculate levels of GADD34 in various experiments by analyzing the calibrated lysate on the same Western blot membranes. To this end, band intensities of full-length GADD34 and GADD34-eGFP were compared. GAPDH signal intensity was used as a loading control to account for different cell numbers in the lysate analyzed.

### Single-cell transcript quantification using RNA fluorescence in situ hybridization (FISH)

To analyze cell-to-cell variability of transcript levels and to discriminate between infection and stress status, we performed single-cell transcript quantification using the ViewRNA ISH Cell Assay Kit (Affymetrix) consisting of target specific RNA FISH probe sets and branch DNA signal amplification for detection of specific signal. 1.5 x 105 Huh7 cells were seeded in a 6-well plate and infected with HCV (MOI = 10 TCID_50_ per cell) or left uninfected. 24 h post infection, cells were treated with 100 IU/ml IFN-α (PBL) or left untreated. 48 h after infection, cells were fixed for 15 min in 4 % PFA in PBS. RNA FISH was performed as follows: cells were permeabilized for 45 min with 100 % ethanol at 4 °C, followed by three washes with PBS. RNA-binding proteins were digested with the Protease QS diluted 1:12,000 in PBS for 5 min at RT. Cells were washed three times with PBS. Cells were incubated for 30 min at 40 °C with the fluorescent probes diluted 1:25 with pre-warmed Probe Set Diluent QF. Cells were washed three times for 10 min with Wash Buffer. Cells were incubated for 30 min at 40 °C with the Label Probe Mix (diluted 1:25 with pre-warmed Probe Set Diluent QF). Cells were washed three times for 10 min with PBS and once with distilled water before mounting coverlips with ProLong Gold Antifade Mountant (Molecular Probes). All fluorescently-labeled probes were purchased from Affymetrix: human EIF2S1 (eIF2α) Type 1 (Alexa Fluor 546, #VA1-20426, diluted 1:50); human PPP1R15A (GADD34) Type 1 (Alexa Fluor 546, # VA1-15768, diluted 1:50); Human PRKR (PKR) Type 4 (Alexa Fluor 488, # VA4-18296, diluted 1:50); HCV (+)ssRNA Type 6 (Alexa Fluor 647, #VF6-13516, diluted 1:100). Probes targeting Bacillus subtilis (dapB) transcripts were used as negative control (Type 1 Alexa Fluor 546 #VF1-11712 and Type 4 Alexa Fluor 488 # VF-4-10408), all diluted 1:50. Oligo(dT)50 probes targeting poly(A) sequences, labeled at the 5’ end with Alexa488 (Invitrogen) or Alexa555 (MWG-Biotech, Eurofins) were used at 50 nM to visualize cell outlines and SGs.

Images of whole cell volume (200 nm Z-spacing) were acquired on a Nikon Ti Eclipse microscope equipped with a Perkin Elmer UltraView Vox spinning disc CSU-X1, using the CFI Apo TIRF 60x (NA 1.49) Oil objective (Nikon). Images were corrected for uneven illumination by division with an illumination bias mask and multiplying each pixel intensity by factor 2000 using the Fiji package software (http://fiji.sc/wiki/index.php/Fiji) (77). Individual masks were generated for each channel by acquiring an image of autofluorescent plastic slides (Chromas) and consequently processing applying the Gaussian blur function in Fiji (radius 50).

To determine transcript levels on a single-cell level, a software (based on Matlab) was developed to manually circle cells of interest and to assign individual phenotype (infected/non-infected and stressed/non-stressed). High-throughput data analysis was performed using Matlab. For each dataset and fluorescent channel, negative control samples (uninfected cells hybridized with probes targeting prokaryotic dapB transcripts and HCV genome) were used to identify fixed size threshold (ST) and intensity thresholds (IT): GADD34+HCV FISH - GADD34 [IT 750/1500, PT 25/15] and HCV [IT 750/1500, PT 10/15]; PKR+HCV FISH – PKR [IT 220/400, PT 10/6] and HCV [IT 500/1800, PT 15/10]; GADD34+HCV FISH in PKROE - GADD34 [IT 6000, PT 10] and HCV [IT 1500, PT 10/]. Applying these values, total transcript levels were calculated on a single-cell, multi-layered phenotype level. Signals exceeding the intensity and size threshold (determined in cells hybridized by negative control probes) are circled in white.

### Single-cell Western blot analysis

10^5^ Huh7 cells (naïve, stably overexpressing PKR or PKRKO clone 2#3) were seeded in 6-well plates. 24 h post seeding, cells were treated with 100 IU/ml IFN-α or left untreated. 24 h after treatment, cells were detached using Accutase (Capricorn). Single cell Western blotting was performed using the Standard scWestern Kit (Proteinsimple) as recommended by the manufacturer. In short, 2 ml of cell suspension (4×105 cells) was loaded on dry slides, allowed to settle for 30 min and carefully washed with suspension buffer. For PKR protein detection, 500 μl of Huh7 PKRKO cell suspension was loaded on a separate section of the microslide as a background control. Well occupation rate (max. 30 %) and duplicates (max 2 % of occupied wells) were monitored manually by light microscopy. Microslides were transferred to the MILO single cell Western system (Proteinsimple), subjected to 10 sec lysis, 90 sec electrophoresis at 240 V and 240 s UV capture. Subsequently, slides were washed three times 15 min in wash buffer and stained in primary antibodies (dilution 1:10 in Antibody Diluent 2) for 2 h, followed by three washes of 15 min. Fluorescent secondary antibodies (dilution 1:20 in Antibody Diluent 2) were incubated for 90 min, followed by three washes of 15 min. For complete salt removal, slides were washed in double-distilled water twice for at least 2 h. Finally, slides were transferred to 50 ml reaction tubes and subjected to dry centrifugation for 1 h at 1,000 × g. Images were acquired using the InnoScan 710 microarray scanner (Innopsys) and the acquisition software Mapix (Innopsys, version 8.1.1). Correction for staining intensity variation was performed using Fiji by division of each image by a blurred mask of itself and subsequent pixel intensity multiplication by 2,000. The masks were generated applying Gaussian blur function (radius 200). Each well was manually curated by filtering out damaged or soiled regions of the chip using Scout 2.1 software (Proteinsimple). To determine total signal intensity per lane, a script was developed to automatically detect centers of wells and to segment total lane areas on each chip. Background intensity was determined using empty wells for eIF2α quantification or PKRKO cells for PKR quantification. The following primary and secondary antibodies were used: mouse monoclonal anti-GAPDH (G-9) (Santa Cruz), rabbit polyclonal anti-eIF2α (Cell Signaling), rabbit monoclonal anti-PKR (Proteintech), goat anti-mouse-Alexa532 (Invitrogen), goat anti-rabbit-Alexa635 (Invitrogen).

### *In vitro* synthesis of single-stranded RNA of positive and negative polarity, hybridization of dsRNA and purification

The sequence of the prokaryotic ampicillin resistance gene was used as template for synthesis of dsRNA (100, 200 and 400 bp). Primers used included the T7 polymerase promoter sequence: Plus_For (5’ - TAA TAC GAC TCA CTA TAG GGA GTA TTC AAC ATT TCC GTG TCG CCC TTA T - 3’); Plus_100bp_Rev (5’ – CAT CTT TTA CTT TCA CCA GCG TTT CTG GGT – 3’); Plus_200bp_Rev (5’ – ATC ATT GGA AAA CGT TCT TCG GGG CGA AAA – 3’); Minus_100bp_For (5’ – TAA TAC GAC TCA CTA TAG GTC TTT TAC TTT CAC CAG CGT TTC TGG GTG A – 3’); Minus_200bp_For (5’ – TAA TAC GAC TCA CTA TAG GCA TTG GAA AAC GTT CTT CGG GGC GAA AAC T – 3’) and Minus_Rev (5’ – ATG AGT ATT CAA CAT TTC CGT GTC GCC CTT – 3’). 40bp-dsRNA including the first 40 bases of ampicillin resistance gene was synthesized by Metabion (5’ – AUG AGU AUU CAA CAU UUC CGU GUC GCC CUU AUU CCC UUU U – 3’). PCR products were purified using the Nucleospin Gel and PCR Clean-Up kit (Macherey-Nagel). Single-stranded RNA of positive and negative polarity was synthesized by *in vitro* transcription and purified as described above, using 3 μg of PCR product as template in a 100 μl reaction. For hybridization, equimolar amounts of respective positive and negative ssRNA were incubated at 85 °C for 10 min in hybridization buffer (25 mM HEPES, 50 mM NaCl) and gradually cooled to room temperature within 1 h. Double-stranded RNA was precipitated by adding 10 % (v/v) 2 M sodium acetate (pH 4.5) and 75 % (v/v) isopropanol and incubation on ice for 2 h. Precipitated dsRNA was pelleted at 20,000 x g for 45 min. Pellets were washed with 70 % (v/v) ethanol and resuspended in RNase-free water. RNA concentration was determined by measuring optical density at 260 nm. RNA integrity and secondary structures were analyzed on denaturing agarose gels.

### Induction of SGs by dsRNA and chemical stressors

1.5 x 10^5^ Huh7 cells or Huh7 YFP-TIA1 cells were seeded in 6-well plates containing glass coverslips. After 24 h, medium was replaced with fresh medium containing varying concentrations of sodium arsenite (Sigma-Aldrich), thapsigargin (Biotrend) or DMSO (Merck Millipore). Samples were analyzed by Western blot, Phos-tag gel or by immunofluorescence analyses. To determine p-PKR and p-eIF2α expression levels and visualize SG formations, cells were treated for 45 min with arsenite or 1 h with thapsigargin. To determine GADD34 expression levels, cells were treated for 8 h with DMSO or thapsigargin. To measure the response to dsRNA, cells were transfected with varying amounts of 200-bp dsRNA using Lipofectamine 2000 (Invitrogen) with a RNA:transfectant ratio of 1:2, according to the manufacturer instructions and harvested after 16 h.

### Quantification of SGs by immunofluorescence assay

To quantify percentage of SG-positive cells after drug treatment, Huh7 cells were fixed for 15 min with 4 % (w/v) paraformaldehyde in PBS, permeabilized by incubation in PBS containing 0.5 % Triton-X100 for 5 min and incubated in blocking buffer (5 % horse serum (C.c.pro), 5 % sucrose in PBS) for 30 min. SGs were visualized using polyclonal rabbit anti-eIF3B (Bethyl Laboratories; 1:1,000) diluted in blocking buffer for 1 h, washed three times with PBS for 5 min and incubated with donkey anti-rabbit secondary antibody coupled to Alexa Fluor 488 (Invitrogen; 1:2,000). Coverslips were washed in PBS as above and mounted on glass slides using Fluoromount G Reagent (Southern Biotech). For Huh7 YFP-TIA1 cells, coverslips were directly mounted on glass slides after fixation. Images were acquired with a Nikon Ti Eclipse fluorescence microscope using a CFI Plan Apo Lambda 20x objective (NA 0.75) (Nikon). Fluorescent signal was captured using an EM-CCD camera C9100 (Hamamatsu) the NIS-Elements AR software package (Nikon, version 4.30). Percentage of SG-positive cells was determined manually using the Cell Counter plugin of the Fiji software package. SG formation in Huh7 YFP-TIA1 cells was determined using the YFP signal.

### Phos-tag polyacrylamide gel analysis

Phos-tag gel analysis was performed to detect mobility shift of eIF2α phosphorylated form as described previously (70). A 10 % resolving polyacrylamide gel was supplemented with 70 μM Phos-tag acrylamide (Fujifilm Wako Chemicals) and 140 μM Mn2+ as recommended by the manufacturer. Before blotting, gels were incubated in transfer buffer (25 mM Tris-HCl pH 8.3, 150 mM glycine, 20 % methanol) supplemented with 1 mM EDTA for 10 min, followed by 10 min washing step in transfer buffer in absence of EDTA. Both basal and phosphorylated forms of eIF2α were visualized using an anti-eIF2α antibody. Signal was detected using the Advance ECL Chemocam Imager (Intas Science Imaging) and band intensity was quantified using the Lab-Image 1D Software (Intas Science Imaging). The percentage of p-eIF2α was determined by dividing intensity of the slowly migrating band (p-eIF2α) with the sum of both band intensities (p-eIF2α + eIF2α).

### Inhibition of thapsigargin-induced SG formation by GADD34 overexpression

For GADD34 lentivirus production, 5×10^6^ HEK 293T cells were seeded into 10cm cell culture dishes. One hour prior to transfection, medium was replenished. For transfection, 6.4 μg pWPI GADD34 Puro or pWPI Puro (control), 6.4 μg packaging plasmid (pCMVΔ8.91) and 2.16 μg of the vector expressing the VSV-G envelope glycoprotein (pMD2.G) were mixed with OptiMEM (Gibco) to a final volume of 400 μl. Polyethylenimine (PEI) (Sigma-Aldrich) was diluted to 112.5 μg/ml in OptiMEM to a final volume of 400 μl. Solutions of plasmid and PEI were mixed, vortexed rigorously and incubated for 20 min at room temperature. Transfection mix was added drop-wise onto producer cells. Medium was replenished after six hours. Lentivirus supernatant was harvested at 48 and 72 h after transfection and filtered with a 0.45 μm pore size membrane. For transient overexpression of GADD34, Huh7 cells were detached by trypsinization, and 105 cells were resuspended in lentivirus supernatant diluted with DMEM complete to a final volume of 2 ml and seeded in 6-well plates. Thirty hours post transduction, cells were treated with 2 μM thapsigargin for 1 h and subsequently harvested for Western blot and immunofluorescence analyses. Lentiviruses expressing the antibiotic resistance gene only (pWPI Puro) were used as control.

### Bacterial expression and purification of His-PKR and His-eIF2α

His-PKR was expressed from pET-His-PKR in E. coli expression strain BL21 (DE3)-RIL (kindly provided by C. Müller, Heidelberg) cultured at 37 °C in autoinduction medium (1 % (w/v) tryptone, 0,5 % (w/v) yeast extracts, 1 mM MgSO_4_, 0.5 % glycerol, 0.05 % glucose, 0.2 % lactose monohydrate, 25 mM (NH_4_)_2_SO_4_, 50 mM KH_2_PO_4_, 50 mM Na_2_HPO_4_, 34 μg/ml chloramphenicol and 50 μg/ml kanamycin) until reaching OD_600_ of 2.0 and then transferred to 18 °C for 20 h. Bacteria were pelleted at 6,000 x g at 4 °C for 15 min and stored at −80 °C. The bacteria were resuspended in 5 ml/g bacteria lysis buffer (50 mM Na_2_HPO_4_, 300 mM NaCl, 20 mM imidazole, 1 mM phenylmethylsulfonyl fluoride (PMSF), cOmplete EDTA-free protease inhibitor cocktail (Roche); pH 8.0) and lysed in three homogenization cycles using EmulsiFlex C3 (Avestin). The lysate was centrifuged at 15,000 x g for 1 h at 4 °C and the supernatant was incubated with Ni-NTA agarose beads (1 ml of 50 % slurry; Macherey & Nagel) for 18 h at 4 °C while tumbling. Beads were pelleted by centrifugation for 10 min 500 x g at 4 °C and washed twice with 10 ml wash buffer (50 mM Na_2_HPO_4_, 300 mM NaCl, 20 mM imidazole).

Protein was eluted with 10 ml elution buffer (50 mM Na_2_HPO_4_, 300 mM NaCl, 250 mM imidazole) in 1.5 ml fractions. Eluate was dialyzed for 16 h in storage buffer (10 mM Tris pH 7.5, 50 mM KCl, 2 mM MgCl_2_, 10 % glycerol, 7 mM β-mercaptoethanol) using dialysis tubes with a 12,000-14,000 MWC. His-PKR was further concentrated to at least 10 mg/ml using Vivaspin centrifugal concentrator 30 K (Sartorius), snap-frozen in liquid nitrogen and stored at −80 °C.

His-eIF2α was expressed as described above with the following modifications: The protein pellet was resuspended in modified lysis buffer (20 mM Tris, 500 mM NaCl, 2 mM β-mercaptoethanol, 20 mM Imidazol, 1 mM PMSF, cOmplete EDTA-free protease inhibitor cocktail (Roche), pH 8.0). Beads were washed in modified wash buffer (20 mM Tris, 500 mM NaCl, 2 mM β-mercaptoethanol, 20 mM imidazol). Protein was eluted using modified elution buffer (20 mM Tris, 500 mM NaCl, 2 mM β-mercaptoethanol, 500 mM imidazol). Fractions containing the protein were identified by SDS-PAGE using Coomassie stain or Bio-Rad TGX Stain-Free FastCast gels (12 %). The fractions containing His-eIF2α were pooled and dialyzed into ion exchange buffer A (20 mM Bis-Tris pH 6.0, 2 mM β-mercaptoethanol) for 16 h, at 4 °C. Precipitate was separated by centrifugation at 15,000 x g, 5 min at 4 °C followed by an anion exchange chromatography loading the supernatant on a HiTrap Q HP 5 ml column (GE Healthcare) and intensive wash step of 20 column volumes with ion exchange buffer A. Protein was eluted by a gradient of 0-100 % ion exchange buffer B (20 mM Bis-Tris pH 6.0, 500 mM NaCl, 2 mM β-mercaptoethanol). Clean fractions were pooled and concentrated to at least 10 mg/ml using Vivaspin centrifugal concentrator 10 K (Sartorius), snap frozen in liquid nitrogen and stored at −80 °C. To remove hyperphosphorylation from His-PKR expressed in E. coli, the purified protein was subjected to dephosphorylation (adapted from (78)). 100 μg of His-PKR was dephosphorylated using 3,200 units of λ-PPase (New England Biolabs) in 200 μl reaction buffer (50 mM HEPES, 100 mM NaCl, 2 mM DTT, 0.01 % Brij 35 pH 7.5, 1 mM MnCl_2_) for 2 h at 37 °C. The dephosphorylation reaction was stopped by addition of 2 μl of 200 mM sodium orthovanadate and further incubation for 5 min at 37 °C. Protein aggregates were removed by centrifugation for 30 min at 20,000 x g at 4 °C. Protein concentration was determined by subjecting samples and dilution series of GST-PKR kinase domain (Abcam) to SDS-PAGE, followed by protein visualization by silver staining and band intensity quantification.

### *In vitro* PKR kinase assay

The protocol was adapted from (78). The reaction mix was prepared on ice in a final volume of 12 μl containing 150 ng of dephosphorylated His-PKR and 1 μg of His-eIF2α in activation buffer (20 mM HEPES pH 7.5, 4 mM MgCl_2_, 100 mM KCl) supplemented with 1 mM Ultra-Pure ATP (Promega). 3 μl of dsRNA dilutions or water were added to the mix and reactions were immediately incubated at 30 °C for 20 min. Potential phosphatase activity was inhibited by the addition of 2 μl of 20 mM sodium orthovanadate. 3.5 μl of 6x Laemmli buffer was added to the reaction mix and samples were boiled at 95 °C for 5 min. Half of the reaction (75 ng PKR, 500 ng eIF2α) was analyzed by SDS-PAGE and total protein was visualized by Silver stain (see below). 1.67 % of the reaction (2.5 ng PKR, 16.7 ng eIF2α) was analyzed by either SDS-PAGE for p-PKR and p-eIF2α levels analysis or by Phos-tag polyacrylamide gel to determine the percentage of p-eIF2α as described above.

### Visualization of PKR and eIF2α recombinant proteins in polyacrylamide gels by Silver staining

Proteins were visualized using the Silver Stain Plus Kit (Bio-Rad) following the manufacturer’s instructions. In short, protein samples were analyzed by SDS-PAGE. After electrophoresis, polyacrylamide gels were incubated in fixing buffer (50 % methanol, 10 % acetic acid, 10 % Fixative Enhancer) for 20 min at room temperature and washed twice for 10 min in double-distilled H_2_O. Proteins were stained in Staining and Developing Solution (5 % Silver Complex Solution, 5 % Reduction Moderator Solution, 5 % Image Developer Solution, 50 % Development Accelerator Solution). When bands became clearly visible, gels were incubated in Stop Solution (5 % acetic acid) for 15 min und subsequently washed with double-distilled H_2_O for at least 2 h.

### Quantitative RT-PCR of GADD34 mRNA species

8 x 10^4^ Huh7 cells were seeded in a 12-well plate and treated with 2 μM thapsigargin for up to 12 h. Cell were harvested at time 0, 1, 3, 6 and 12 h post treatment and total RNA was extracted as described above. Levels of mature GADD34 mRNA were determined using the qPCRBIO Probe 1-Step Go Lo-ROX, PCR Biosystems kit (Nippon Genetics). In brief, 15 μl reaction mixes contained 7.5 μl 2 x qPCRBIO mix, 0.75 μl 20 x RTase, 400 nM per primer, 200 nM per probe and 3 μl of extracted total RNA. For absolute quantification, serial 1:10 dilution series (10^3^ to 10^9^ copies) of GADD34 and GAPDH transcripts were processed on each plate. Each sample was measured in triplicate wells. qRT-PCR was performed on a C1000 Touch Thermal Cycler with a CFX96 Real-Time System (Bio-Rad) using the following settings: 50 °C for 10 min, 95 °C for 1 min and 40 cycles as follows: 95 °C for 10 sec and 60 °C for 1 min. The following primers and probes were used to detect cellular transcripts: GAPDH_For: 5’ – GAA GGT GAA GGT CGG AGT C – 3’; GAPDH_Rev: 5’ – GAA GAT GGT GAT GGG ATT TC – 3’; GAPDH_Probe: 5’ – VIC – CAA GCT TCC CGT TCT CAG CCT – TAMRA – 3’; GADD34mRNA_For: 5’ – CAG AAA CCC CTA CTC ATG ATC C – 3’, GADD34mRNA_Rev: 5’ – AAA TGG ACA GTG ACC TTC TCG – 3’, GADD34_Probe: 5’ – FAM – CCC CTA AAG GCC AGA AAG GTG CGC – TAMRA – 3’. GAPDH copy number was used for normalization. For the detection of GADD34 pre-mRNA, 30 μl of total RNA were incubated with 1 μl TURBO DNAse and TURBO DNase buffer (all from Thermo Fisher) in a total volume of 50 μl at 37 °C for 30 min to remove residual genomic DNA contaminations. Following RNA purification, 3 μl of DNAse-digested RNA were reverse transcribed into cDNA using the Applied Biosystems High Capacity cDNA Reverse Transcription kit (Thermo Fisher) as recommended by the manufacturer (25 °C for 10 min, 37°C for 2 h and 85°C for 5 min). The cDNA was diluted 1:20 prior to detection of GADD34 pre-mRNA transcripts using iTaq Universal SYBR Green (Bio-Rad). In short, 15 μl mix contained 7.5 μl 2x iTaq mix, 500 nM per primer and 3 μl of pre-diluted cDNA. Each sample was measured in triplicate wells. qRT-PCR was performed on a C1000 Touch Thermal Cycler with a CFX96 Real-Time System (Bio-Rad) using the following settings: 95 °C for 3 min and 45 cycles as follows: 95 °C for 10 sec and 60 °C for 30 sec. The following primers were used: GADD34prem_For: 5’ – ACA GTG ACA GGC AAG TGA CTA G – 3’, and GADD34prem_Rev: 5’ – GGA AGA GAG AGA GAG AAG CAA AC – 3’. GAPDH_For: 5’ – GAA GGT GAA GGT CGG AGT C – 3’; GAPDH_Rev: 5’ – GAA GAT GGT GAT GGG ATT TC – 3’. GAPDH mRNA was used for normalization of input RNA. RT-PCR data were analyzed by using the ΔΔCT method described previously (79).

### Statistical analysis

Statistical analysis was performed by using the GraphPad Prism software (version 7.04). Statistical significance for Western blot, Phos-tag gel and SG immunofluorescence analyses was calculated by performing two-way ANOVA with Dunnett’s multiple comparisons test. Statistical significance for FISH analyses was calculated by performing two-tailed unpaired t-test with Welch’s correction. *p<0.05; **p<0.01; ***p<0.001; ****p<0.0001.

## Supporting information

Klein, Kallenberger_Supplementary_Materials

## Acknowledgments

We thank Monika Langlotz from the ZMBH Flow Cytometry & FACS Core Facility (Heidelberg University) for her support in cell sorting, Vibor Laketa from the Infectious Diseases Imaging Platform (IDIP) at the Center for Integrative Infectious Disease Research (Heidelberg), Dirk Scholz and Klaus Nettesheim from Nikon Instruments Germany for their assistance and support with live-cell microscopy. We thank Gunter Stier and Irmgard Sinning (BZH, Heidelberg University) for kindly providing the pET His 1a vector. We thank Hanspeter Herzel, Marco Binder and Jens Timmer for critical discussions.

## Funding

This study was supported by:

Deutsche Forschungsgemeinschaft (DFG, German Research Foundation) – Project number 240245660 (TP13 to AR, TP8 to OTF, TP11 to RB and TH, Z4 to FAH and KRoh)

DFG – Project number 278001972 (A14 to AR and GS, A21N to TH)

DFG – Project number 272983813 (TP12 to AR and TP9 to RB)

Equipment program of Cellnetworks Cluster of Excellence (EXOC 81) (funds for MILO single cell Western system to OTF).

Bundesministerium Bildung und Forschung (BMBF) – Project number 031L0270 to SMK (Computational Life Sciences program).

## Author contributions

AR conceived the study, designed and interpreted the experiments, and wrote the manuscript with input from all authors. PK additionally designed, performed, analyzed the experiments, and edited the manuscript. SMK designed and performed all computational analyses and mathematical modeling, supported image analysis, and edited the manuscript. HR, VM performed experiments. TBNLH generated recombinant His-tagged PKR and eIF2α proteins. PK, KRot, JA, YQ, SW performed the image analysis under the guidance of FAH and KRoh. OO provided technical expertise with FISH analysis. RK and BDV developed a preliminary version of the deterministic model. RE, RB and OTF provided guidance and infrastructure. SRT, OTF, TH and GS reviewed and edited the manuscript. Competing interests: Authors declare that they have no competing interests.

## Data and materials availability

All data needed to evaluate the conclusions in the paper are present in the paper and/or the Supplementary Materials. Cell lines and plasmids are available upon materials transfer agreements. Additional data related to the image processing pipeline and mathematical model script may be requested to the authors.

