## Supplementary material for "Temporal control of the integrated stress response by a stochastic molecular switch": Klein, Kallenberger_Supplementary_Materials

**This file includes:**

Supplementary Text: Developments of mathematical models

Figs. S1 to S14

Tables S1 to S4

Movies S1 to S3

References (1 to 12)

**Other Supplementary Materials for this manuscript include the following:**

Movies S1 to S3

### Supplementary Text: Developments of mathematical models

#### 1. Descriptive models of stress granule fluctuations

For drawing parallels to processes that might be explanatory for the observed frequency spectra and autocovariances of stress granule (SG) appearance in single cells, we implemented three different models for transitions between an 'On' and an 'Off' phase – a random telegraph process, an oscillator and a process with transitions according to two gamma distributions (Fig. 1G). A random telegraph process was simulated by uniform sampling of switching times between an 'On' and an 'Off' phase simulated for 50 days with 4 time points per hour. Average Fourier transforms and autocovariances were calculated from n=500 simulations. An oscillator was simulated with two switching events per day for 3 days. Histograms of log10-scaled SG phase and break lengths in hours were phenomenologically described by a weighted sum of two gamma distributions,

$$f(x) = \frac{s_1}{b_1\Gamma(a_1)} x^{a_1-1} e^{-x/b_1} + \frac{s_2}{b_2\Gamma(a_2)} x^{a_2-1} e^{-x/b_2}. \quad (1.1)$$

By fitting histograms of untreated or IFN-α-treated HCV-infected cells, we obtained the following parameter estimates:

| | $s_1$ | $a_1$ | $b_1$ | $s_2$ | $a_2$ | $b_2$ |
|---|---|---|---|---|---|---|
| HCV, peaks | 45.93 | 2.848 | 0.5949 | 44.07 | 21.70 | 0.2104 |
| HCV, breaks | 40.03 | 1.645 | 1.611 | 22.65 | 37.03 | 0.1069 |
| HCV+IFN-α, peaks | 86.61 | 5.819 | 0.5053 | 14.48 | 54.14 | 0.01645 |
| HCV+IFN-α, breaks | 69.33 | 0.7055 | 1.501 | 57.52 | 15.76 | 0.2144 |

To simulate analogous random processes with transitions reflecting short or long gamma-distributed peaks or breaks, we sampled from

$$h(u, g_1, g_2) = ug_1 + (1-u)g_2 \quad (1.2)$$

with gamma distributed $g_1 \sim \frac{1}{b_1\Gamma(a_1)} x^{a_1-1} e^{-x/b_1}$, $g_2 \sim \frac{1}{b_2\Gamma(a_2)} x^{a_2-1} e^{-x/b_2}$ and $u$ being uniformly distributed in the interval [0,1]. Again, average Fourier transforms and autocovariances were calculated from n=500 simulations for untreated or IFN-α-treated HCV-infected cells.

Experimentally observed frequency spectra and autocovariances could only reproduced by sampling gamma-distributed peaks and breaks. Observed frequency spectra of SG fluctuations were continuously decreasing to higher frequencies, as in case of the telegraph model. However, autocovariances with negative values around lags of one day could be only reproduced by sampling from gamma-distributions. An interesting analogy can be drawn to a theoretical study of neurite growth, a process that continuously switches between growth and shortening phases, however, on the time scale of minutes (*1*). In this study, a gamma distribution of phase times was derived from a model accounting for a series of intermediate steps between 'On' and 'Off' phases.

#### 2. PKR activation model

It was described that, in cells, PKR binds to dsRNA, forms dimers and is activated by auto-phosphorylation (*2*). We observed that the amount of phosphorylated PKR (p-PKR) in Huh7 cells 8 hours after transfection with dsRNA increased up to an amount of 1 µg. However, lower levels of p-PKR resulted when transfecting with higher amounts of dsRNA up to 6 µg (Fig. 3, A and B). Similarly, we observed a bell-shaped dependency of the PKR activity on the dsRNA concentrations in experiments of PKR activation *in vitro* by dsRNA of three different fragment lengths (40 bp, 100 bp, 200 bp; Fig. 4A, fig. S5). Of note, in these experiments, peak levels of PKR activity increased with dsRNA fragment lengths (Fig. 4). Our observations are consistent with previous findings of a bell-shaped dependency of PKR activity under *in vitro* conditions on the concentration of (*3*).

The bell-shaped dependency of PKR activity on the dsRNA concentration can be explained by taking into account the ratio between binding sites in dsRNA and PKR molecules. Starting at low ratios between dsRNA and PKR, all possible binding sites will be engaged with PKR oligomers. An increase in dsRNA concentration will first result in an increased amount of oligomerized PKR bound at dsRNA. However, in case the ratio of binding sites to PKR molecules is strongly increased, the largest number of binding sites can be occupied by binding of PKR monomers. Therefore, the number of oligomers, and thus PKR activity, will be reduced in presence of large dsRNA concentrations.

We developed an optimal mechanistic model of PKR activation, consisting of coupled ordinary differential equations (ODEs), for our dataset of *in vivo* and *in vitro* experiments (Fig. 4, B to D, fig. S6, fig. S7). Measurements of p-PKR were taken at a time when PKR phosphorylation was at steady state. Therefore, experimental data was fitted with model species at steady state. To this end, model simulations were performed for an integration time of 3 days, parameters for binding were estimated, whereas parameters for unbinding were fixed to large values ($1min^{-1}$) to enforce convergence of model variables to steady state. Thereby, we determined ratios between parameters describing binding of PKR to dsRNA, $k_{d,on}$ and $k_{d,off}$, as well as parameters describing formation of oligomerized PKR, $k_{p,on}$ and $k_{p,off}$.

The stepwise model refinement and underlying mechanistic assumptions are visualized in fig. S6A. A total of 28 model variants with 4 to 14 species that contained between 15 and 20 estimated parameters (fig. S6) were calibrated with our experimental dataset of p-PKR measurements. The dataset comprised measurements from cells transfected with 200-bp dsRNA fragments using 6 different amounts (0 to 6 μg) and *in vitro* assay measurements at 12 dsRNA concentrations (0 to 1 μM) per fragment length in 40 bp, 100 bp and 200 bp dsRNA fragments resulting in a total number of 45 data points. For parameter estimations based on maximum likelihood estimation the MATLAB toolbox PottersWheel was used (*4*). A total of 500 multi-start local optimizations were conducted for each model variant. The corrected Akaike information criterion was used for model selection. Equations and observable definitions for selected model variants are reported in Table S1. Parameter estimates and parameter confidence intervals for the optimal model variant as part of a model of the cellular stress response are reported in Table S4.

Our first model 'Variant 1' took into account independent reactions for PKR dimerization and binding to dsRNA, different PKR affinities and binding capacities depending on the dsRNA fragment length (fig. S6). We observed that extending the model by reactions of higher PKR oligomers (n=2…6) improved the model fit indicated by a reduction in the $\chi^2$ measure as well as the corrected Akaike information criterion (ΔAICc, fig. S6). Further extending 'Variant 1' by reactions for PKR activation by cis and trans mechanisms did not result in an improvement ('Variant 1.1'), whereas, excluding oligomerization reactions of PKR without dsRNA (consistent with large $K_d$ values for free PKR) improved the model fit ('Variant 2'). Having observed that the decline of PKR activity at high dsRNA concentrations could be better described by including reactions to higher oligomers of PKR, we formulated a model accounting for cooperative binding of PKR to PKR:dsRNA complexes ('Variant 3') which resulted in a strong improvement of the model fit. Cooperative recruitment implies that binding of PKR to dsRNA facilitates binding of additional PKR molecules, resembled by a sigmoidal dependency of PKR binding on the PKR concentration, but is not indicative of the exact stoichiometry of PKR complexes.

Further *in vitro* studies investigated possible mechanisms of PKR activation by *cis* (intramolecular) or *trans* (intermolecular) reactions (*5-7*). Therefore, we additionally tested model variants ('Variant 4' to 'Variant 5.1') with or without cooperative PKR binding and optional *cis* or *cis* and *trans* reactions for PKR activation that, however, did not result in an improvement relative to the optimal model 'Variant 3'. Of note, the observation that it was not necessary to include *cis*/*trans* reactions for PKR activation in our model could be due to the fact that our dataset did not include time-resolved measurements of PKR activation but rather represent PKR activities at steady state.

Taken together, our dataset was optimally explained by a model that describes recruitment of PKR to dsRNA and PKR oligomer formation at dsRNA with cooperativity and takes into account different affinities of PKR for dsRNA of varying fragment lengths (40 bp, 100 bp, 200 bp), as well as for PKR oligomerization at dsRNA fragments of varying fragment lengths. Model equations and definitions of observables for the optimal variant are listed in Table S1. A recent study based on a mutational analysis of PKR and molecular dynamics simulations proposed that PKR protomers interact via front-to-front as well as back-to-back interfaces (*7*). Notably, these different modes of interactions between PKR protomers might provide an explanation for our observation that the bell-shaped dependency of PKR activity on the dsRNA concentration is optimally described by cooperative PKR recruitment.

### 3. GADD34 degradation model

To determine, whether GADD34 degradation is reflected by Michaelis-Menten kinetics or by an exponential decay, we fitted two alternative ODE models to measurements of GADD34 concentrations in Huh7 cells stably over-expressing GADD34 after inhibiting new synthesis by cycloheximide,

$$\frac{d[GADD34]}{dt} = -k_{deg,GADD34}[GADD34] \quad (3.1)$$

and

$$\frac{d[GADD34]}{dt} = -k_{deg,GADD34}\frac{[GADD34]}{K_{deg,GADD34}+[GADD34]}, \quad (3.2)$$

with degradation rate constant $k_{deg,GADD34}$ and Michaelis-Menten constant $K_{deg,GADD34}$. To fit the experimental dataset, observables were defined by $y_{GADD34} = s_{GADD34}[GADD34] + y_{bg,GADD34}$ with the scaling factor $s_{GADD34}$ and the background value $y_{bg,GADD34}$ to account for the background signal in Western blot measurements. To limit the weight of single data points with negligible SEM values, we estimated a lower limit of residual weights. To this end, we fitted a linear error model $\varepsilon_i = ax_i + b\max(x_i)$, describing the experimental uncertainty $\varepsilon_i$ depending on measurements $x_i$ for data points with index $i$ and coefficients $a$ and $b$, to standard errors of the mean depending on average GADD34 concentrations. Thereafter, the linear coefficient $a = 2.62nM$ was used as lower bound to weight residuals for model fitting. For each of the two models, a total of 100 multi-start local optimizations were conducted followed by profile likelihood estimation to determine parameter confidence intervals (Supplementary Text Fig. 1). Model comparison showed that Eq. (1) was superior to Eq. (2), indicated by an increase in the corrected Akaike information criterion of $\Delta AIC_{corr} = 18.4$. It can be concluded that the Michaelis-Menten constant $K_{deg,GADD34}$ for GADD34 degradation exceeds experimental GADD34 concentrations. We estimated a degradation rate constant of $k_{deg,GADD34} = 0.01854$ (1σ-C.I., [0.01516, 0.02289]) reflecting a GADD34 half-life of about 37 minutes.

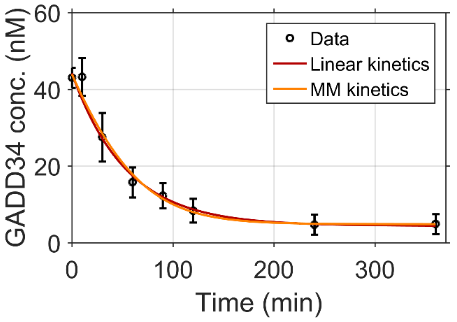


**Supplementary Text Fig. 1. GADD34 degradation.** A comparison between linear or Michaelis-Menten (MM) kinetics indicates that GADD34 degradation can be sufficiently well described by an exponential decay Shown are mean of n=4 replicates ± SEM. Lines, best-fits of GADD34 degradation models.

### 4. Deterministic model of the integrated stress response

To quantitatively study the integrated stress response, we constructed an ODE model accounting for cellular stress mediated by PKR (dsRNA stress), HRI (oxidative stress) and PERK (ER stress). The core model of dsRNA, oxidative and ER stress (Fig. 2A) is described by equations listed in Table S2. It contains 18 species and a total of 26 parameters. In the following, we will describe the model implementation and underlying assumptions.

In the model, cellular stress is either evoked by presence of dsRNA resulting in PKR activation, PERK activation by thapsigargin or HRI activation by arsenite. Stress kinase activation results in phosphorylation of eIF2α. We assumed that stress kinase activation is fast compared to eIF2α phosphorylation and GADD34 expression. Therefore, stress kinase activity, depending on dsRNA, thapsigargin or arsenite, is described at steady state in Eqs. 5–10 as further detailed below (Table S2). In the model, phosphorylated eIF2α is denoted by $eIF2\alpha^*$. Moreover, SG formation depending on the concentration of phosphorylated eIF2α and translational inhibition by cellular stress are described as fast and cooperative processes, reflected by Hill-type functions ($f_{SG,PKR}$, $f_{SG,A}$ and $f_{SG,T}$ in Eqs. 6, 8 and 10; Table S2). Depending on these functions, the *ppp1r15a* promoter $Pr$ becomes activate. Promoter activation is modeled as a process with a temporal delay to account for intermediate steps not accessed by experiments (reactions involving ATF4 and CHOP, formation of DNA-protein complexes) based on a linear chain of reactions ("linear chain trick"; Eqs. 11–16 in Table S2) (*8*). The active promoter $Pr^*$ causes expression of GADD34 mRNA. GADD34 is only expressed dependent on GADD34 mRNA in presence of cellular stress due to regulation by an upstream open reading frame (*9*). This was accounted for by the dependency of GADD34 translation on the Hill-type function for SG formation and translational repression $f_{SG}$ (Eq. 18 in Table S2). GADD34 expression results in dephosphorylation of $eIF2\alpha^*$. In a first model implementation, this GADD34-dependent dephosphorylation was described by Michaelis-Menten kinetics. Parameter estimations, however, resulted in large estimates of the $K_M$ value for GADD34 that exceeded the average cellular concentration of total $eIF2\alpha$ ($\overline{[eIF2\alpha]} = 2244\text{nM}$). Therefore, GADD34-dependent dephosphorylation was described with linear kinetics, which did not result in reduced fit quality.

In all Western blot measurements, we observed that, in absence of stressors, a small fraction of eIF2α was phosphorylated, which can be attributed to a background phosphorylation rate. In the model, background phosphorylation is described by the kinetic parameter $k_{ph,basal}$ (Fig. 3, A and D, fig. S4E, fig. S10, B, D and E). Basal dephosphorylation by CReP (constitutive repressor of eIF2α phosphorylation) (*10*) was accounted for by the parameter $k_{CReP}$ (Eqs. 5–10, Table S2).

The integrated stress response was experimentally characterized in cells infected with HCV, transfected with dsRNA, treated with arsenite or thapsigargin. To combine complementary information from different experiments and improve parameter identifiability, we performed multi-experiment fitting to a combination of experimental datasets. To this end, the core model had to be adapted to boundary conditions of each experimental dataset. In the following, we describe the eight model parts with partially shared kinetic parameters that were simultaneously fitted to experimental datasets. An overview of observables, assignments, algebraic equations, scaling factors, background offsets of all model parts is given in Table S3.

In most experimental datasets, measurements were only taken at single time points for different concentrations of the stress stimulus (arsenite, dsRNA, thapsigargin) or overexpressed GADD34. These measurements were either recorded at an early time point soon after applying the stressor or at a late time point after adaptation to a stressor. Time course measurements of eIF2α phosphorylation, SG percentage and GADD34 expression were recorded after thapsigargin stimulation (fig. S10F). These time-resolved measurements indicated that GADD34 expression was not induced after one-hour thapsigargin exposition, whereas GADD34 expression was maximal after 6 hours of thapsigargin exposition. Maximal levels of p-eIF2α were reached at 1 hour after thapsigargin stimulation. In general, measurements at single time points are insufficient for fitting a time-resolved function. For this reason, we applied steady state assumptions for model parts fitted to datasets containing measurements at single time points for varying stress stimuli or initial protein concentrations.

When fitting single measurements of p-eIF2α and SG percentage at 1 hour after applying a stressor, we assumed that these model species were at steady state. Because at that time GADD34 expression in response to a stressor is not induced, only equations for phosphorylation and dephosphorylation of eIF2α as well as SG formation were taken into account (Eqs. 5–10, Table S2). In combination with equations accounting for the effect of a stressor, adaptation of the system for basal eIF2α phosphorylation and dephosphorylation by CReP before application of a stressor was described by including an additional set of model Eqs. 7–18 (Table S2) with stressor concentrations set to zero for describing basal levels in model parts 4.3, 4.4 and 4.6 as described below and illustrated in Supplementary Text Fig. 3–10. Thereby, we simulated the effect of a stressor starting at basal enzyme activities for (de)phosphorylation of eIF2α but could simultaneously apply the steady state assumption for eIF2α phosphorylation and SG formation in response to the applied stressor. For model fitting of steady state concentrations, these model parts were simulated for an integration time of 3 days for different stress stimuli or initial protein concentrations, which was sufficient to enforce steady states of model variables in case of multi-experiment fitting.

When fitting single measurements at 6 hours (model part 4.4), 8 hours (model part 4.2) or later (model parts 4.7, 4.8), the whole set of equations describing eIF2α (de)phosphorylation and GADD34 expression was simulated for an integration time of 3 days to enforce steady states. Additionally, the dynamics of eIF2α (de)phosphorylation and GADD34 expression could be inferred by combined fitting to steady state and time-resolved experimental data (model part 4.5).

In all model parts, the initial concentration of eIF2α was set to the measured average concentration $\overline{[eIF2\alpha]} = 2244\text{nM}$. The initial concentration of PKR was fixed to the experimentally determined average concentration of $\overline{[PKR]} = 25.24\text{nM}$ for untreated cells or $\overline{[PKR]}_{IFN} = 75.48\text{nM}$ in cells treated with IFN-α. The initial concentration of GADD34 mRNA was set to $\overline{[mGADD34]} = 0.00571\text{nM}$. This concentration was derived from an average GADD mRNA molecule number in untreated cells divided by the average cell volume. The initial value of the inactive promoter species $Pr$ was set to 1. Initial values of all other species were set to zero at the start of the integration time. Because GADD34 mRNA degradation was not measured, we fixed the kinetic parameter for GADD34 mRNA degradation to $k_{deg,m} = 0.0035min^{-1}$ as previously reported (*11*).

By fitting the combined model to a total of 241 data points, a total of 78 parameters (including scaling factors and auxiliary parameters) were estimated. A total of 500 multi-start local optimizations were conducted followed by profile likelihood estimation to determine parameter confidence intervals. Parameter estimates, parameter bounds and parameter confidence intervals are listed in Table S4. In the following, the implementation of model parts describing different experimental datasets will be explained.

#### 4.1 Model of PKR activation by *in vitro* assay

The model of PKR activation under *in vitro* conditions was fitted to measurements of phospho-PKR obtained from an experiment with purified PKR protein and dsRNA. It is equivalent to the optimal PKR activation model and consists of Eqs. 1–4 in Table S2. The model describes binding of PKR to dsRNA fragments of 40 bp, 100 bp and 200 bp length (with parameter index $i \in \{1,2,3\}$ of Eqs. 1–4 in Table S2) and cooperative formation of oligomerized PKR at dsRNA (Supplementary Text Fig. 2).

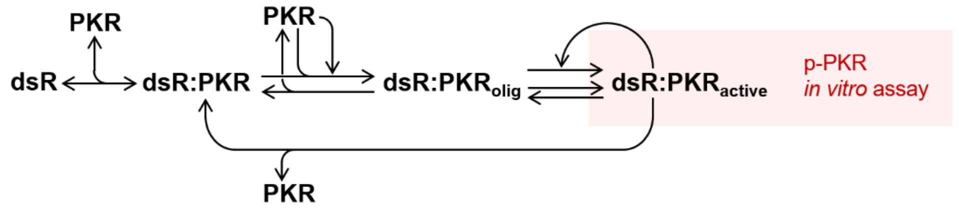


**Supplementary Text Fig. 2. Model for describing PKR *in vitro* assay.** Measurements of p-PKR were associated with the model species $dsR{:}PKR_{active}$ representing active PKR (red area).

We assumed different binding constants for binding of PKR monomers as well as oligomer formation. Further, we assumed different factors of binding sites in 100 bp and 200 bp fragments relative to the number of binding sites in 40-bp fragments in start value assignments of the model species $dsR$ of binding sites in dsRNA (Eqs. 1–3, Table S3). Observables contained a scaling factor independent of fragment length and background intensities (Eqs. 4–6, Table S3). The upper limit of background intensities was set to the minimal measured value of each dataset (Supplementary Table S4). Assuming steady states of PKR phosphorylation, the model was simulated for an integration time of 3 days to enforce convergence to steady states of model variables for parameter estimation. Parameter estimates indicated that the binding capacity for PKR molecules was about 11-fold higher in 100-bp fragments and about 23-fold higher in 200-bp fragments relative to the binding capacity of 40-bp fragments.

#### 4.2 Model of PKR activation by dsRNA transfection

To describe measurements of p-PKR, p-eIF2α (Western blot measurements of p-eIF2α, and % p-eIF2α measurements by Phos-tag gels) and SG percentage at 8 hours after transfection with 200-bp dsRNA, the PKR model was extended by reactions describing PKR-dependent and basal eIF2α phosphorylation, *ppp1r15a* promoter activation (with time delay), GADD34 mRNA and protein turnover as well as GADD34-dependent and basal (CReP-dependent) dephosphorylation of p-eIF2α (Supplementary Text Fig. 3).

The model part consists of Eqs. 1–6 and 11–18 in Table S2 (with parameter index $i = 4$ in Eqs. 1–4). Similar as in model part 4.1, the transfected dsRNA amount was related by a scaling factor to the initial number of PKR binding sites $dsR$. The start value assignment and for $dsR$ and implementations of observables for this model part are described by Eqs. 7–11 in Table S3.

Model simulations were performed for an integration time of 3 days to fit measurements by the model at steady state. As described in section '2. PKR activation model', we estimated ratios between parameters describing binding of PKR to dsRNA, $k_{d,on}$ and $k_{d,off}$, as well as parameters describing PKR oligomerization at dsRNA, $k_{p,on}$ and $k_{p,off}$. Thereby, however, we only could obtain an estimate of equilibrium dissociation constants for the binding of free PKR, $K_{d,PKR} = \frac{k_{p,off}}{k_{p,on}} = 10.9nM$, but not for PKR oligomerization, due to the non-linear dependency of PKR oligomerization on the PKR concentration (Eqs. 1–4, Table S2).

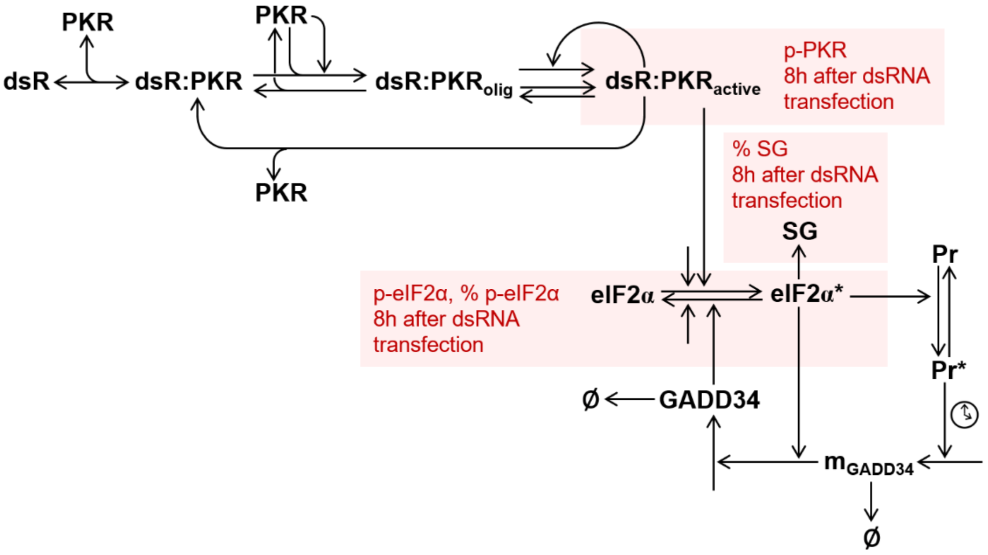


**Supplementary Text Fig. 3. Model for describing dsRNA-induced cellular stress.** Measurements of p-PKR, % SG, p-eIF2α and % p-eIF2α were associated with model species $dsR{:}PKR_{active}$, $eIF2\alpha$, $eIF2\alpha^*$, $SG$ as indicated by red areas.

Stress granule formation was described by the Hill-function

$$f_{SG,PKR} = \frac{[eIF2\alpha*]^l}{K_{SG,PKR}^l + [eIF2\alpha*]^l} \quad (4.1)$$

with Hill-coefficient $l$ and the parameter $K_{SG,PKR}$ equal to the $eIF2\alpha^*$ concentration at 50% SG presence, assuming that SG formation is a fast and cooperative process. Model fits of this part are shown in fig S10B.

#### 4.3 Arsenite titration model

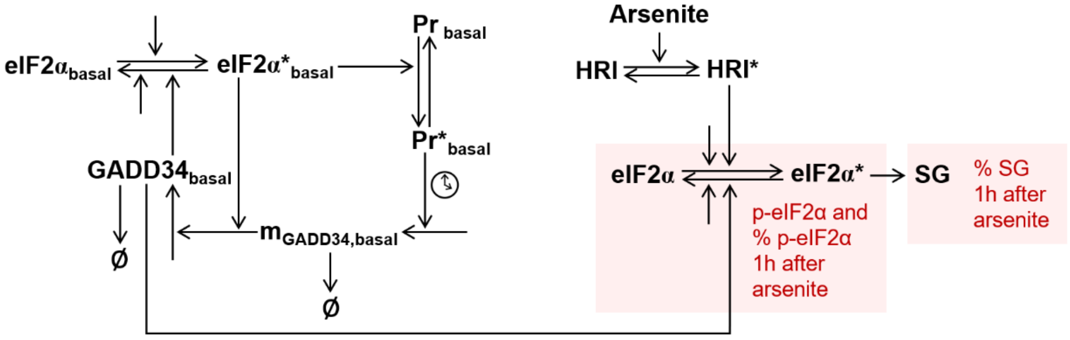


**Supplementary Text Fig. 4. Model for describing oxidative stress induced by arsenite.** Measurements of p-eIF2α, % p-eIF2α and % SG after 1-hour arsenite exposition were associated with model species $eIF2\alpha^*$ and $SG$ simulated at steady states for eIF2αphosphorylation.

To simulate eIF2α phosphorylation and SG formation 1 hour after exposition to arsenite for different arsenite concentrations (0 to 500µM), basal steady states of model species ($eIF2\alpha_{basal}$, $eIF2\alpha^{*}{}_{basal}$, $Pr_{basal}$, $Pr^{*}{}_{basal}$, $mGADD34_{basal}$, $GADD34_{basal}$) were simulated together with concentrations of eIF2α and phosphorylated eIF2α ($eIF2\alpha^{*}$) resulting from active HRI ($HRI^{*}$, Supplementary Text Fig. 4). We assumed that activation of HRI by arsenite was at steady state and that the concentration of active HRI could be described by

$$[HRI^{*}] = \frac{[HRI]_{tot}[A]}{K_A + [A]} \tag{4.2}$$

depending on the arsenite concentration $[A]$, the total HRI concentration $[HRI]_{tot}$ and the apparent dissociation constant $K_A$. In model reaction equations for $eIF2\alpha$ and $eIF2\alpha^{*}$ we summarized the kinetic parameter describing eIF2α phosphorylation by $HRI^{*}$ and $[HRI]_{tot}$ in the estimated parameter $k_{ph,A}$ with unit $nM/min$ describing the enzyme activity of HRI (Eqs. 7 and 8, Table S2). Implementations of model observables for this part are described by Eqs. 13–15 in Table S3. Stress granule formation was described by a Hill-function $f_{SG,A}$, similar as in model part 4.2, with Hill-coefficient $l$ and parameter $K_{SG,A}$ (Eq. 16 in Table S3). Model fits of this part are shown in Fig. 5B and fig. S10D.

#### 4.4 Thapsigargin titration model

The thapsigargin titration model was calibrated with measurements of eIF2α phosphorylation and SG formation at 1 hour and GADD34 expression at 6 hours after exposition to thapsigargin for different thapsigargin concentrations (0 to 2 µM). GADD34 protein was measured by normal and calibrated Western blotting (fig. S3C and fig. S10E). Average molecule numbers of GADD34 per cell obtained from calibrated Western blots were used to calculate average GADD34 concentrations according to the average cell volume of $V_c = 6.71pl$.

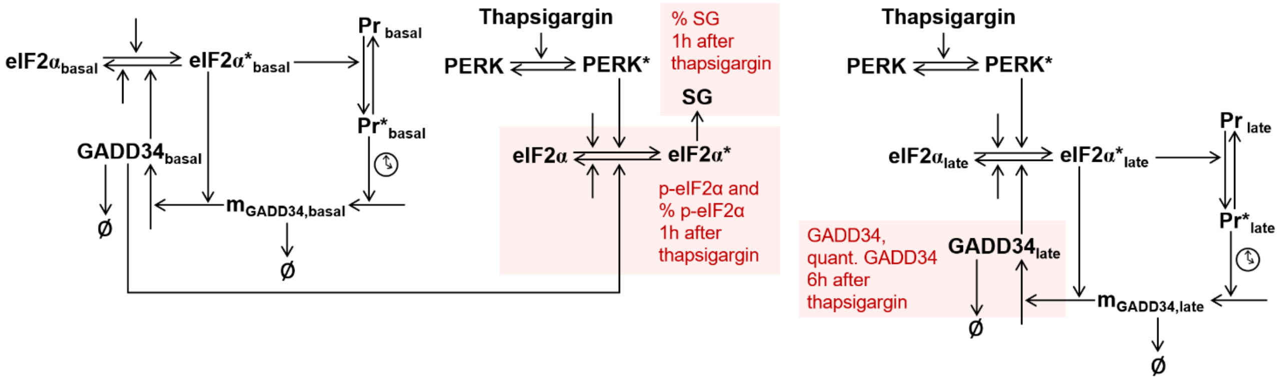


**Supplementary Text Fig. 5. Model for describing ER stress induced by thapsigargin.** Measurements of p-eIF2α, % p-eIF2α and % SG at 1h as well as GADD34 concentrations at 6h after exposition to thapsigargin (GADD34 measurements in arbitrary units from normal Western blot experiments and quantitative GADD34 measurements from calibrated Western blots) were associated with model species $eIF2\alpha^{*}$ and $SG$ (at steady states for eIF2α phosphorylation) and $GADD34_{late}$ (at steady states for eIF2α phosphorylation and GADD34 expression).

To fit measurements of p-eIF2α and SG percentages after 1-hour thapsigargin treatment, basal steady states of model species were simulated together with concentrations of eIF2α and phosphorylated eIF2α ($eIF2\alpha^{*}$) resulting from active PERK ($PERK^{*}$; Supplementary Text Fig. 5). Assuming that PERK activation was at steady state, the concentration of active PERK could be described by

$$[PERK^{*}] = \frac{[PERK]_{tot}[T]}{K_T + [T]} \tag{4.3}$$

depending on the thapsigargin concentration $[T]$, the total PERK concentration $[PERK]_{tot}$ and the apparent dissociation constant $K_T$. As in case of HRI, the kinetic parameter describing eIF2α phosphorylation by $PERK^{*}$ and $[PERK]_{tot}$ were summarized in the estimated parameter $k_{ph,T}$ (Eqs. 9 and 10, Table S2). Observable definitions for this model part are described by Eqs. 17–20 in Table S3. SG formation was described by a Hill-function $f_{SG,T}$, with Hill-coefficient $l$ and the parameter $K_{SG,T}$ (Eq. 21, Table S3). Model fits of this part are shown in Fig. 5B and fig. S10E.

### 4.5 Thapsigargin kinetics model

The thapsigargin kinetics model was used to describe time-resolved eIF2α phosphorylation and GADD34 expression after thapsigargin exposition at a concentration of 2 µM (Supplementary Text Fig. 6). It consists of Eqs. 9–18 in Table S2. Observable definitions for this model part are described by Eqs. 22–24 in Table S3.

It was used to estimate parameters for the dynamics of promoter activation, GADD34 mRNA and protein synthesis in combination with the other model parts. To fit measurements between 0 and 8 hours after adding thapsigargin, the model was first simulated for an integration time for 3 days to enforce convergence to basal steady states of model variables before simulating the response to thapsigargin-induced stress at time points between 0 and 8 hours (fig. S10F).

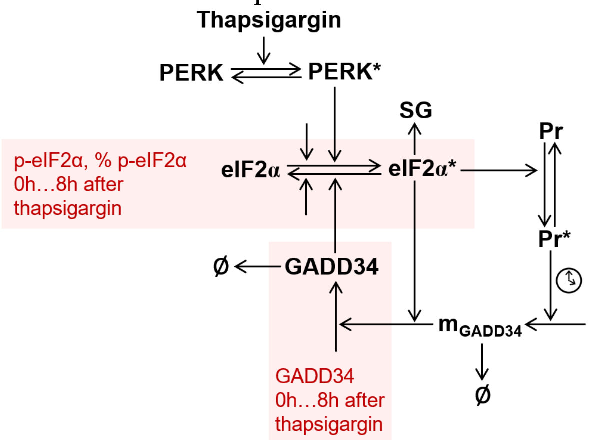


**Supplementary Text Fig. 6. Model for describing time-resolved ER stress response.** Measurements of p-eIF2α, % p-eIF2α and GADD34 were associated with model species $eIF2\alpha^*$ and $GADD34$.

### 4.6 GADD34 titration model

To study the dose-response of eIF2α dephosphorylation depending on the concentration of GADD34, we overexpressed GADD34. Huh7 cells were infected with different concentrations using different doses of a retrovirus encoding for GADD34 or stably transfected. In addition to normal Western blot measurements, calibrated Western blot were conducted to determine average molecule numbers per cell, and average concentrations were estimated based on the average cell volume (fig. S3B). Concentrations of exogenously expressed GADD34 varied between 17.1nM (lowest retrovirus dose) and 123.1nM (stable overexpression). In GADD34-overexpressing and control cells, p-eIF2α and SG percentage were measured at 1 hour after thapsigargin exposition, at a time when endogenously expressed GADD34 was not induced. As expected, we observed a sigmoidal decrease of p-eIF2α at higher GADD34 concentrations (fig. S10G).

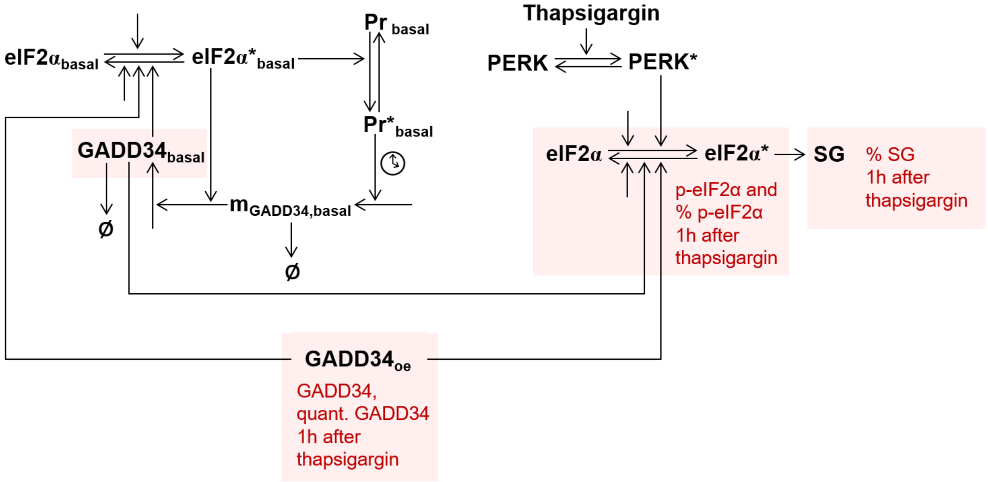


**Supplementary Text Fig. 7. Model for describing ER stress in case of GADD34 overexpression.** Measurements of p-eIF2α, % p-eIF2α and % SG at 1h after exposition to thapsigargin as well as GADD34 measurements from normal and quantitative Western blotting (quant. GADD34) were associated with model species $eIF2\alpha^*$ and $SG$ (at steady states for eIF2α phosphorylation), $GADD34_{basal}$ (control cells) or $GADD34_{oe}$ (overexpressing cells).

To fit measurements after 1h thapsigargin treatment originating from basal steady states, as in model parts 4.3 and 4.4, a set of model species describing concentrations at basal steady states described by Eqs. 9–18 (table S2) with a thapsigargin concentration set to zero was included in the model (Supplementary Text Fig. 7). Equations describing basal levels were combined with Eqs. 9 and 10 from table S2 to simulate phosphorylation of eIF2α after thapsigargin exposition. The equations for describing basal and thapsigargin-dependent concentrations of (un)phosphorylated eIF2α were modified by including the concentration of overexpressed expressed GADD34 ($[GADD34_{oe}]$):

$$\frac{d[eIF2\alpha_{basal}]}{dt} = -k_{ph,basal}[eIF2\alpha_{basal}] - k_{ph,T}\frac{[eIF2\alpha_{basal}]}{K_{ph,T} + [eIF2\alpha_{basal}]} +k_{CReP}[eIF2\alpha^{*}{}_{basal}] + k_{deph}[eIF2\alpha^{*}{}_{basal}]([GADD34_{basal}] + [GADD34_{oe}]) \tag{4.4}$$

$$\frac{d[eIF2\alpha^{*}{}_{basal}]}{dt} = k_{ph,basal}[eIF2\alpha_{basal}] + k_{ph,T}\frac{[eIF2\alpha_{basal}]}{K_{ph,T} + [eIF2\alpha_{basal}]} -k_{CReP}[eIF2\alpha^{*}{}_{basal}] - k_{deph}[eIF2\alpha^{*}{}_{basal}]([GADD34_{basal}] + [GADD34_{oe}]) \tag{4.5}$$

$$\frac{d[eIF2\alpha]}{dt} = -k_{ph,basal}[eIF2\alpha] - k_{ph,T}\frac{[T]}{K_T + [T]}\frac{[eIF2\alpha]}{K_{ph,T} + [eIF2\alpha]} +k_{CReP}[eIF2\alpha *] + k_{deph}[eIF2\alpha *]([GADD34_{basal}] + [GADD34_{oe}]) \tag{4.6}$$

$$\frac{d[eIF2\alpha^{*}]}{dt} = k_{ph,basal}[eIF2\alpha] + k_{ph,T}\frac{[T]}{K_T + [T]}\frac{[eIF2\alpha]}{K_{ph,T} + [eIF2\alpha]} -k_{CReP}[eIF2\alpha *] - k_{deph}[eIF2\alpha *]([GADD34_{basal}] + [GADD34_{oe}]). \tag{4.7}$$

In case of GADD34 overexpression, the variable $GADD34_{oe}$ was fitted to GADD34 measurements from normal and quantitative Western blot experiments (parameters $GADD34_{ex,1}$ to $GADD34_{ex,7}$ in Table S4). In control cells, the variable $GADD34_{basal}$ was fitted to GADD34 measurements. Assignments and model observables for this part are described by Eqs. 25–31 in Table S3. Model fits of this part are shown in fig. S10, G and H.

4.7 FISH model

In single cells, numbers of GADD34 mRNA molecules were detected using FISH probes in combination with labeled poly(dT) to indicate presence of cellular stress. To relate SG percentage with GADD34 mRNA expression, single-cell data were binned. To define edges of bins, cells were ranked according to FISH measurements of HCV positive-sense ssRNA copies. Thereby, cells with similar amount of viral RNA were grouped. Edges were determined by collecting equal cell numbers per bin. Average SG percentages were calculated, and standard errors of SG percentages were estimated by bootstrapping with $10^4$ samples per bin. For FISH experiments in HCV-infected untreated or IFN-α-treated cells, average concentrations of GADD34 mRNA in each bin were calculated relative to the average cell volume. Thereby, 19 data points of GADD34 concentrations and SG percentages were obtained (no cells were assigned to one bin interval for IFN-α-treated cells).

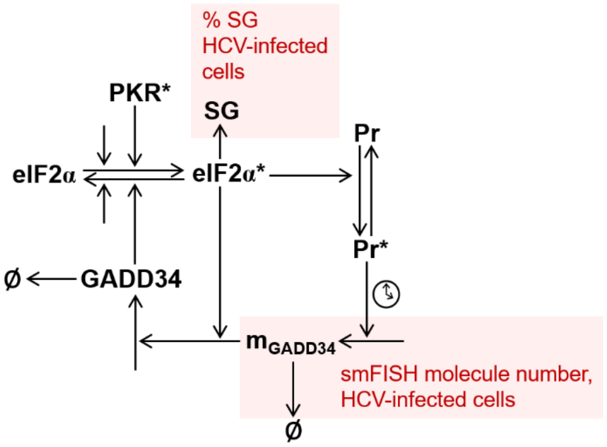


**Supplementary Text Fig. 8. Model for describing FISH measurements in HCV-infected cells.** Measurements of GADD34 mRNA and % SG were associated with model species $mGADD34$ and $SG$.

To describe the relation between SG percentage and GADD34 mRNA, this model part contained Eqs. 5–18 from Table S2. The species $PKR^*$ was described as an auxiliary parameter in this model part that was estimated for each data point (parameters $PKR\_act_1$ to $PKR\_act_{19}$ in Table S4). Assignments and observable definitions for this model part are defined by Eqs. 32–34 in Table S3. To fit the model at steady state to experimental data, the integration time was set to 3 days. Model fits of this part are shown in fig. S10A.

<u>4.8 Model of HCV-infected cells dependent on the infection marker</u>

For live-cell imaging experiments, we used a reporter construct that indicates virus replication (fig. S1, A and B). Assuming that the level of virus replication was proportional to the amount of dsRNA, which is formed as an intermediate during virus replication, we fitted the deterministic model of the cellular stress response to SG percentages depending on average infection marker levels. To this end, we calculated average infection marker intensities and percentages of SG presence in single cells recorded in live-cell microscopy experiments. Experimental measurements were collected in 10 bins defined by ranking cells to infection marker intensities. Edges of infection marker intensities were defined by collecting equal cell numbers per bin.

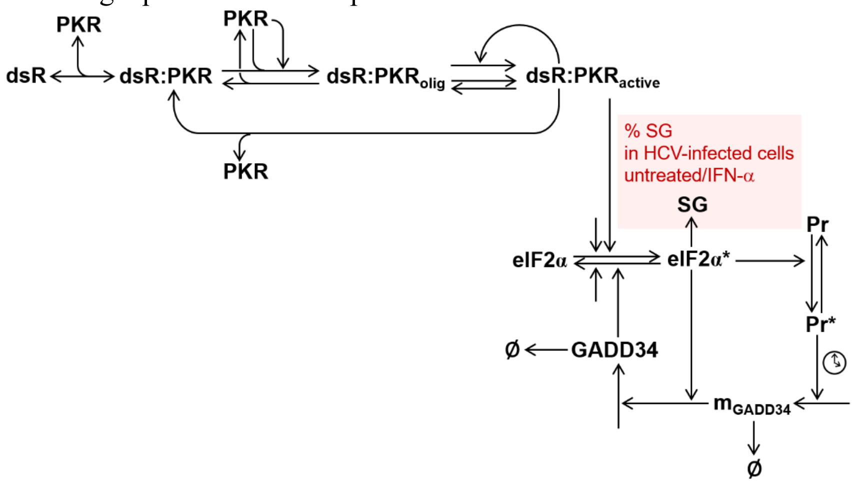


**Supplementary Text Fig. 9. Model for describing infection marker intensities in HCV-infected cells.** Infection marker intensities were related to initial values of PKR binding sites $dsR$ by a scaling factor. The model equation for SG presence was fitted to measurements of SG percentage.

The model describing infection marker intensities and SG percentage values contained Eqs. 1–6 and 11–18 from Table S2. Infection marker levels were assumed proportional to initial concentrations of dsRNA. This assumption was translated to a start value assignment for the model species $dsR$ (Eq. 36 in Table S3). We separately fitted datasets for HCV-infected cells that were untreated or treated with IFN-α. To this end, initial concentrations of PKR were set to experimentally measured average values in untreated ($\overline{[PKR]} = 25.24\text{nM}$) or IFN-α-treated cells ($\overline{[PKR]_{IFN}} = 75.48\text{nM}$). Assignments and observable definitions for this model part are defined by Eqs. 36–38 in Table S3. To fit the model at steady state to experimental data, the integration time was set to 3 days. Model fits of this part are shown in fig. S10I.

### 5. Stochastic model of the integrated stress response

The ODE model describing dsRNA-induced stress in HCV-infected cells (Table S4, Eqs. 5, 6, 11–18) was translated to a stochastic model version describing reactions downstream of active PKR. To this end, kinetic parameters were transformed to reaction propensities. According to the average cell volume of $V_c = 6.71$pl that was determined from confocal microscopic stack images, a concentration of 1nM was equivalent to a number of Nc molecules per cell. Accordingly, kinetic parameters with unit $nM$ or $nM^{-1}$ were multiplied with $N_c$ or $1/N_c$ for scaling to reaction propensities. The stochastic model contained a total of 10 species and 13 reaction propensities. Stochastic model simulations were performed using the matLeap toolbox that makes use of the tau-leaping algorithm (*12*).

To simulate heterogeneous cell populations, we determined the parameters μ and σ of initial log-normal distributions of PKR and eIF2α concentrations as well as viral infection marker, $\mathcal{LN}(\mu_{PKR}, \sigma_{PKR})$, $\mathcal{LN}(\mu_{eIF2\alpha}, \sigma_{eIF2\alpha})$ and $\mathcal{LN}(\mu_{IM}, \sigma_{IM})$. Average cellular concentrations of PKR and eIF2α were determined from calibrated Western blots using GST-tagged proteins (Supplementary Fig. 10E and F). Parameters μ and σ of log-normal distributions were derived by combining these average concentrations with distributions of single-cell Western blot measurements. From log-scaled single-cell Western blot measurements, we determined $\sigma_{PKR,HCV}$ for untreated HCV-infected cells, $\sigma_{PKR,HI}$ for HCV-infected cells treated with IFN-α, and $\sigma_{PKR,HP}$ for HCV-infected PKR-overexpressing cells. Untreated, IFN-α-treated and PKR-overexpressing cells did not differ in expression of eIF2α. Therefore, we assumed the same distribution parameters for these groups and determined only one parameter from single-cell Western blot measurements. Based on parameters σ and average concentrations $m$, parameters μ of log-normal distributions were calculated from $\mu = \log(m) - \sigma^2/2$. Parameters of the log-normal distribution for initial infection marker intensities were calculated form live-cell imaging data. Parameters used for sampling initial values are documented in the following table.

| Log-normal distributions of initial values | μ | σ |
|---|---|---|
| PKR, HCV (nM) | 3.1310 | 0.4416 |
| PKR, HCV+IFN-α (nM) | 4.2585 | 0.3615 |
| PKR, HCV+$PKR_{oe}$ (nM) | 5.0575 | 0.7412 |
| eIF2α (nM) | 7.5942 | 0.4934 |
| Infection marker, HCV (a.u.) | 7.1831 | 1.1809 |
| Infection marker, HCV+IFN-α (a.u.) | 7.6117 | 0.4813 |

To simulate initial conditions for the stochastic model, initial values were sampled for PKR, eIF2α and the infection marker. Then, steady states of active PKR species ($[dsR{:}PKR_{olig}]$) were simulated by the deterministic model of the integrated stress response (Eqs. 1–4 in Table S2) and used as input for stochastic model simulations. Initial values of dsRNA binding sites were calculated by multiplying the sampled initial infection marker values by the estimated scaling factor $f_{sites,HCV}$ (Eq. 36 in Table S3). To use time courses of viral dsRNA as input for stochastic model simulations (Fig. 7) we evaluated average infection marker intensities in untreated or IFN-α-treated HCV-infected cells (Supplementary Text Fig. 10).

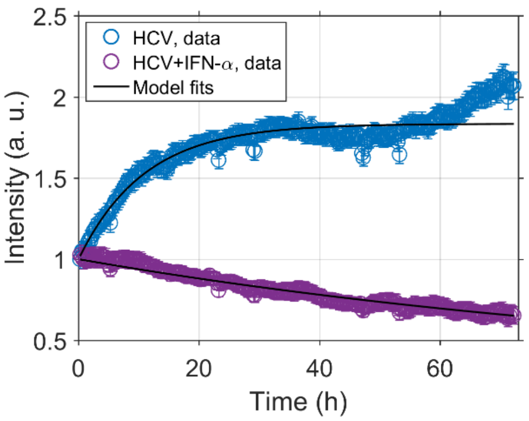


**Supplementary Text Fig. 10. Infection marker time courses.** Average time courses of the infection marker intensity, normalized to the average initial value, in untreated (blue circles) or IFN-a-treated (pink circles) HCV-infected cells measured in live-cell experiments (error bars: SEM). Time courses were fitted by an exponential function to determine turnover parameters.

An exponential function of the form

$$I(t) = I_0\left(1 - \frac{k_{I,syn}}{k_{I,deg}}\right)\exp\left(-k_{I,deg}t\right) + \frac{k_{I,syn}}{k_{I,deg}} \tag{5.1}$$

that describes synthesis and degradation of the infection marker $I$ with parameters $k_{I,syn}$ and $k_{I,deg}$, starting from an initial intensity $I_0$, and represents the solution of an ODE describing synthesis and degradation of the infection marker $dI/dt = k_{I,syn} - k_{I,deg}I$, was fitted to the normalized average trajectories. We obtained parameter estimates $k_{I,syn,HCV} = 0.168h^{-1}$ and $k_{I,deg,HCV} = 0.0917h^{-1}$ in untreated HCV-infected cells and $k_{I,syn,HI} = 0.00220h^{-1}$ and $k_{I,deg,HI} = 0.00869h^{-1}$ in IFN-α-treated HCV-infected cells.

To simulate time courses of PKR activity in a heterogenous cell population of $n$ cells with indices $i = 1 \dots n$, we sampled initial infection marker values $I_{0,i} \sim \mathcal{LN}(\mu_{IM}, \sigma_{IM})$ and simulated infection marker time courses in single cells

$$I_i(t) = I_{0,i}\left(1 - \frac{k_{I,syn,i}}{k_{I,deg}}\right)\exp\left(-k_{I,deg}t\right) + \frac{k_{I,syn,i}}{k_{I,deg}} \tag{5.2}$$

using single cell parameters for synthesis $k_{I,syn,i} = I_{0,i}k_{I,syn}$. Thereby, we assumed that each single cell infection marker intensity was changed during live-cell imaging experiments by the same factor $\frac{k_{I,syn}}{k_{I,deg}}$ as the population average ($f_{HCV} \approx 1.83$ in untreated and $f_{HI} \approx 0.253$ in IFN-α-treated cells). The concentration time course of dsRNA binding sites

$$[\mathrm{dsR}](t)_{HCV,i} = f_{sites,HCV}I_i(t) \tag{5.3}$$

was calculated using the estimated scaling factor $f_{sites,HCV}$. To simulate time resolved PKR activity from estimated dsRNA concentrations in single cells, we modified the model part describing PKR activation (Eqs. 1–4, Table S2) by including turnover of dsRNA binding sites described by parameters $k_{I,syn,i}$ and $k_{I,deg}$

$$\frac{d[PKR]_i}{dt} = -k_{d,on}\alpha_{HCV}[PKR]_i[dsR]_i + k_{d,off}[dsR{:}PKR]_i$$

$$-k_{p,on}[dsR{:}PKR]_i[PKR]_i\frac{[PKR]_i^{\,h}}{\left(K_{p,on}\gamma_{HCV}\right)^h + [PKR]_i^{\,h}} + k_{p,off}\left[dsR{:}PKR_{olig}\right]_i \tag{5.4}$$

$$+k_{I,deg}\left([\mathrm{dsR}]_i + [dsR{:}PKR]_i + 2\left[dsR{:}PKR_{olig}\right]_i\right)$$

$$\frac{d[dsR]_i}{dt} = -k_{d,on}\alpha_{HCV}[PKR]_i[dsR]_i + k_{d,off}[dsR{:}PKR]_i + k_{I,syn,i} - k_{I,deg}[\mathrm{dsR}]_i \qquad (5.5)$$

$$\begin{aligned}\frac{d[dsR{:}PKR]_i}{dt} &= k_{d,on}\alpha_{HCV}[PKR]_i[dsR]_i - k_{d,off}[dsR{:}PKR]_i \\ &-k_{p,on}[dsR{:}PKR]_i[PKR]_i\frac{[PKR]_i{}^h}{(K_{p,on}\gamma_{HCV})^h+[PKR]_i{}^h} + k_{p,off}\left[dsR{:}PKR_{olig}\right]_i \\ &-k_{I,deg}[dsR{:}PKR]_i\end{aligned} \qquad (5.6)$$

$$\begin{aligned}\frac{d\left[dsR{:}PKR_{olig}\right]_i}{dt} &= k_{p,on}[dsR{:}PKR]_i[PKR]_i\frac{[PKR]_i{}^h}{\left(K_{p,on}\gamma_{HCV}\right)^h + [PKR]_i{}^h} \\ &-k_{p,off}\left[dsR{:}PKR_{olig}\right]_i - k_{I,deg}\left[dsR{:}PKR_{olig}\right]_i.\end{aligned} \qquad (5.7)$$

Thereby, single-cell time courses of active PKR species $[dsR{:}PKR_{olig}]$ were simulated that served as time-dependent input for stochastic simulations. These were performed in subsequent 15-minute intervals with updated PKR activity values in each interval. To reproduce experimental data of SG presence in single cells, we simulated $N = 200$ single-cell trajectories per experimental condition over 3 days. Presence of SGs was defined based on the Hill-function for SG formation by $f_{SG} \geq 0.5$.

For an independent prediction of the live-cell imaging dataset recorded in PKR-overexpressing cells by stochastic model simulations (fig. S13E), we used the same initial conditions for eIF2α and the infection marker as in HCV-infected untreated cells ($\mu_{eIF2\alpha}$, $\sigma_{eIF2\alpha}$, $\mu_{IM}$, $\sigma_{IM}$, $k_{I,syn,HCV}$ and $k_{I,deg,HCV}$), and only adjusted the parameters of the log-normal distribution of initial PKR concentrations.

During evaluation of live-cell imaging data, short fluctuations of SG appearance or removal (peaks and breaks) of one 15-minute time point were regarded as noise and therefore removed assuming that SG formation and resolution takes place on a longer time scale. For comparison with experimental measurements of single-cell SG time series, short peaks and breaks of one 15-minute time point were removed from SG time series simulations. Comparisons of stochastic simulations and experimental measurements of single-cell SG time series are shown in Fig. 7.

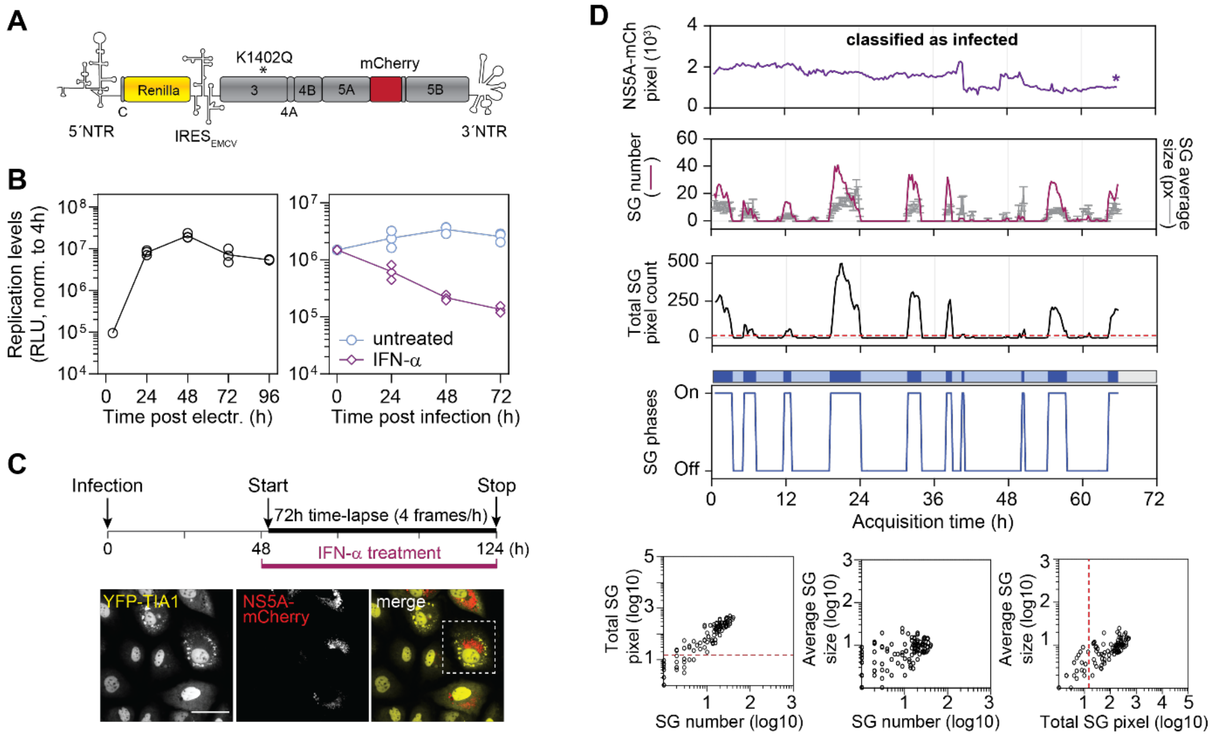


**Fig. S1. Characterization of $HCV_{TCP}$ system and live-cell time-lapse imaging.** (**A**) Schematic representation of HCV RLuc-mCherry bicistronic replicon used for the production of $HCV_{TCP}$. HCV 5' non-translated region (NTR) drives the translation of *Renilla* luciferase that is fused N-terminally to 16 codons of the Core protein of HCV. The internal ribosome entry site (IRES) of the encephalomyocarditis virus (EMCV) directs the translation of the HCV sequence NS3 to NS5B. K1402Q (indicated with a star) is a mutation in NS3, which increases assembly efficiency. The sequence of mCherry was inserted in fusion with NS5A. (**B**) Replication efficiency of HCV RLuc-mCherry replicon in Huh7 cells (n=3). Cells were lysed at the time points specified in the bottom to determine *Renilla* luciferase activity (Relative Light Units, RLU). Values were normalized to the 4 h time point post electroporation (left panel). Right panel: cells infected with $HCV_{TCP}$ for 48 h were treated with 100 IU/ml IFN-α for up to 72 h, or left untreated. *Renilla* luciferase activity was determined and values normalized to the time point 0 of IFN-α addition (n=3). (**C**) Experimental procedure for the time-lapse imaging of SG response dynamics in $HCV_{TCP}$-infected cells. Huh7 YFP-TIA1 cells were infected with $HCV_{TCP}$. Forty-eight hours post infection, cells were treated with 100 IU/ml IFN-α or left untreated and transferred to the microscope heating chamber for live-cell time-lapse microscopy. Images were acquired with an interval of 15 min for 72 h. Representative still images of the YFP-TIA1 and NS5A-mCherry channels are shown on the bottom. The white dotted square corresponds to the cell shown in Fig. 1A. Scale bar, 100 µm. (**D**) Example of complete single-cell time-lapse analysis output including for each time frame (i) the measurement of NS5A-mCherry signal intensity (total pixel count), (ii) the number and average size (± SD) of SGs and (iii) the total SG pixel count. The star indicates the end of a cell track. Cells with a NS5A-mCherry signal intensity above $10^3$ pixels were classified as "infected". Only cells tracked for at least 48 h were considered for the analysis. The red dotted line indicates the threshold of 15 total SG pixels below which cells were not considered to have stress. From these parameters, SG response time series reflecting the sequence of SG-On and SG-Off phases were deduced for each cell, and schematized in the top with dark blue regions for SG-On phases and clear blue regions for SG-Off phases.

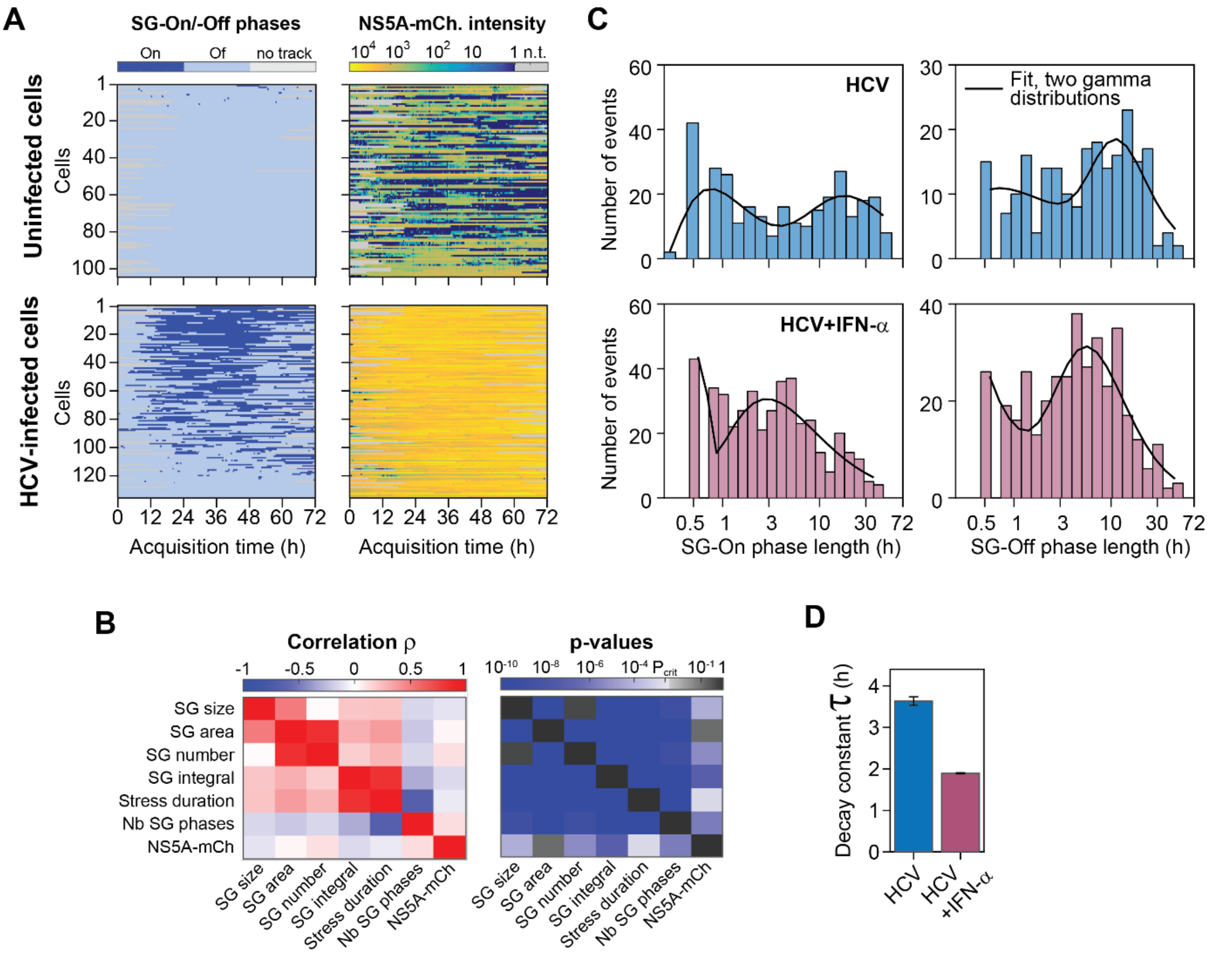


**Fig. S2. Single-cell SG response time series characteristics**. (**A**) Single-cell SG response time series. Shown are SG phases (left panels) and NS5A-mCherry intensity (right panels) from uninfected (n=105) and HCV-infected cells (n=135). Only cells tracked for more than 48 h were considered for further analysis. Single-cell SG response time series were sorted based on the total stress duration, with cell 1 on the top having more stress than cells on the bottom of the panel. (**B**) Significant correlations are visualized by blue colors ($p<p_{crit}=0.05/21$, Bonferroni correction for 21 comparisons). Spearman correlation coefficients (left panel) indicate positive (red) to negative (blue) correlations between SG size, area, number, SG integral and NS5A-mCherry intensity levels. The number of SG phases correlated negatively with all SG parameters except for the NS5A-mCherry intensity levels. Correlation of single-cell parameters (right panel; p-Values from Spearman rank correlation). (**C**) Histograms of experimentally measured SG-On phase lengths and SG-Off phase lengths together with fits of two joint-gamma distributions. (**D**) Exponential decay parameters from fits to the autocorrelation functions shown in (Fig. 1F).

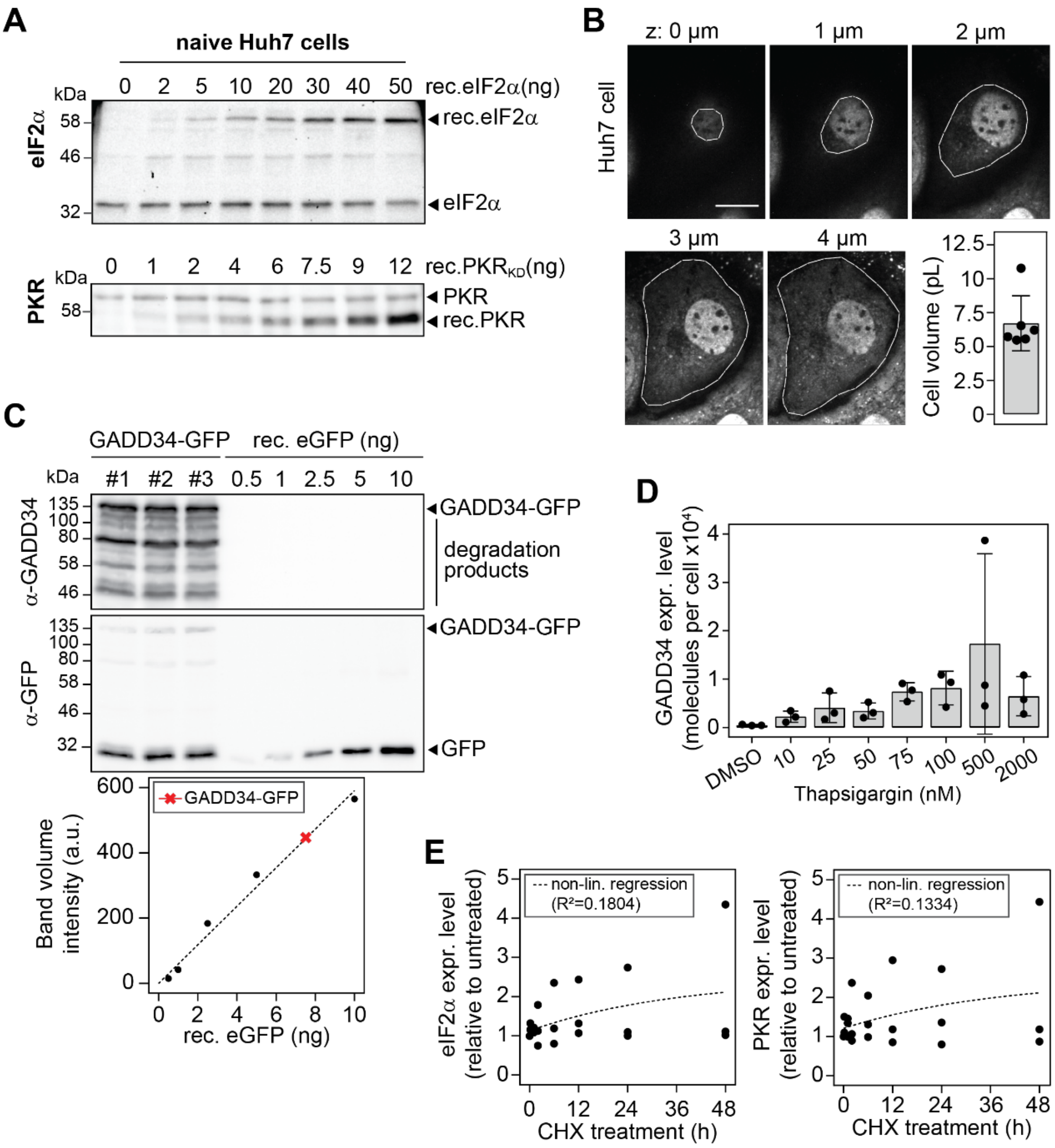


**Fig. S3. Quantification of model species.** (**A**) Absolute quantifications of eIF2α and PKR average molecules per cell. Lysates of defined numbers of Huh7 cells were spiked with increasing amounts of recombinant GST-tagged eIF2α (rec. eIF2α, top panel) or GST-tagged PKR kinase domain (rec. $PKR_{KD}$, bottom panel) and analyzed by quantitative Western blotting. (**B**) Determination of Huh7 YFP-TIA1 cells mean volume by confocal analysis. Cells were imaged in 3D using Z-stacks spaced by 1 µm. Shown are 5 Z-sections from a representative cell starting from the top (0 µm). For each slice, the YFP-TIA1 signal was used to manually outline the border of the cell (white line). Scale bar, 20 µm. The mean cell volume ± SD (n=6) was estimated to be approximately 6.71 pL. (**C**) Absolute quantifications of GADD34-GFP in Huh7 GADD34-GFP cell lysates (n=3). GADD34-GFP levels were compared to amounts of recombinant eGFP (rec. eGFP). Shown are representative Western blot analyses using GADD34-and eGFP-specific antibodies (top and bottom, respectively) and quantification. (**D**) Induction of GADD34 expression in Huh7 cells treated with increasing concentrations of thapsigargin. Cell lysates were analyzed by quantitative Western blotting using Huh7 GADD34-GFP as loading reference. Shown are mean GADD34 molecules per cell ± SD (n=3). (**E**) Analysis of PKR and eIF2α half-lives. Huh7 cells were treated with CHX for 48 h and analyzed by Western blotting. Shown is the quantification of PKR (left panel) and eIF2α (right panel) expression levels normalized to β-actin levels and relative to untreated cells. The dotted lines depict the best non-linear fit (one-phase decay).

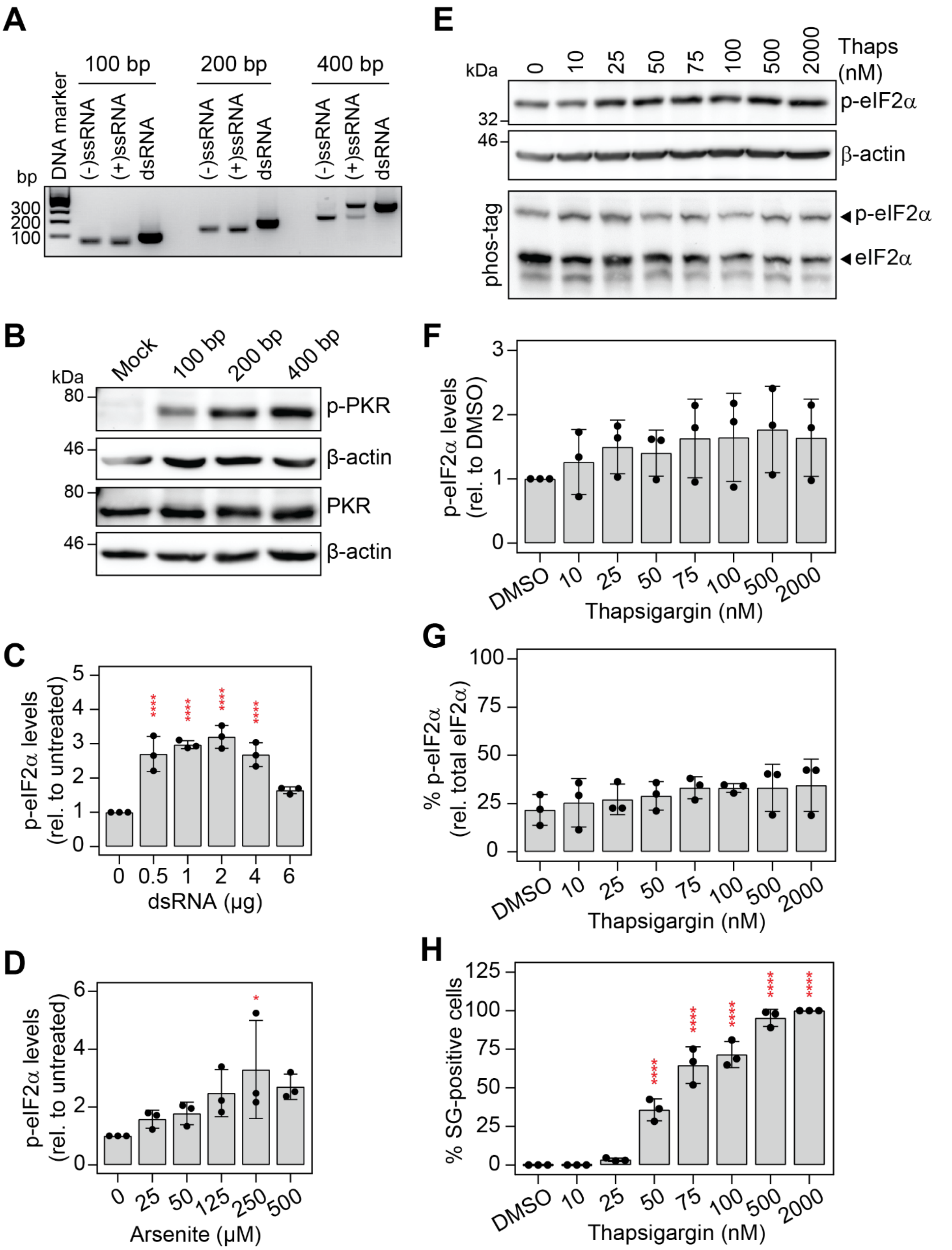


**Fig. S4. PKR activation by dsRNA.** (**A**) Analytical agarose gel of *in vitro* synthesized single-stranded RNAs of negative (-) and positive (+) polarity before and after hybridization (dsRNA). Different RNA lengths were analyzed (100, 200 and 400 bp). (**B**) Analysis of PKR activation in Huh7 cells transfected with dsRNAs of different lengths (15,5 nmol). Shown is a representative Western blot analysis of PKR and p-PKR expression levels. Expression level of β-actin was used as loading control. (**C**) Quantification of p-eIF2α levels in Huh7 cells transfected with increasing amount of 200-bp dsRNA (related to Fig. 3A), normalized to β-actin levels and relative to untreated cells (n=3). Statistical significance is indicated compared to untreated cells; ****p<0.0001. (**D**) Quantification of p-eIF2α levels in Huh7 cells treated with increasing arsenite concentrations (related to Fig. 3D), normalized to β-actin levels and relative to untreated cells (n=3). Statistical significance is given in the top; *p<0.05. (**E to H**) Huh7 cells were treated with increasing amounts of thapsigargin. Shown are representative Western blot and Phos-tag polyacrylamide gel analyses upon 1 h treatment (mean ± SD, n=3) (E), corresponding quantification of p-eIF2α levels normalized relative to β-actin levels and relative to cells treated with DMSO (F) and the percentage of p-eIF2α relative to total eIF2α. (H) The corresponding percentage of SG-positive cells was analyzed by fluorescence microscopy. Statistical significance is indicated compared to cells treated with DMSO; ****p<0.0001.

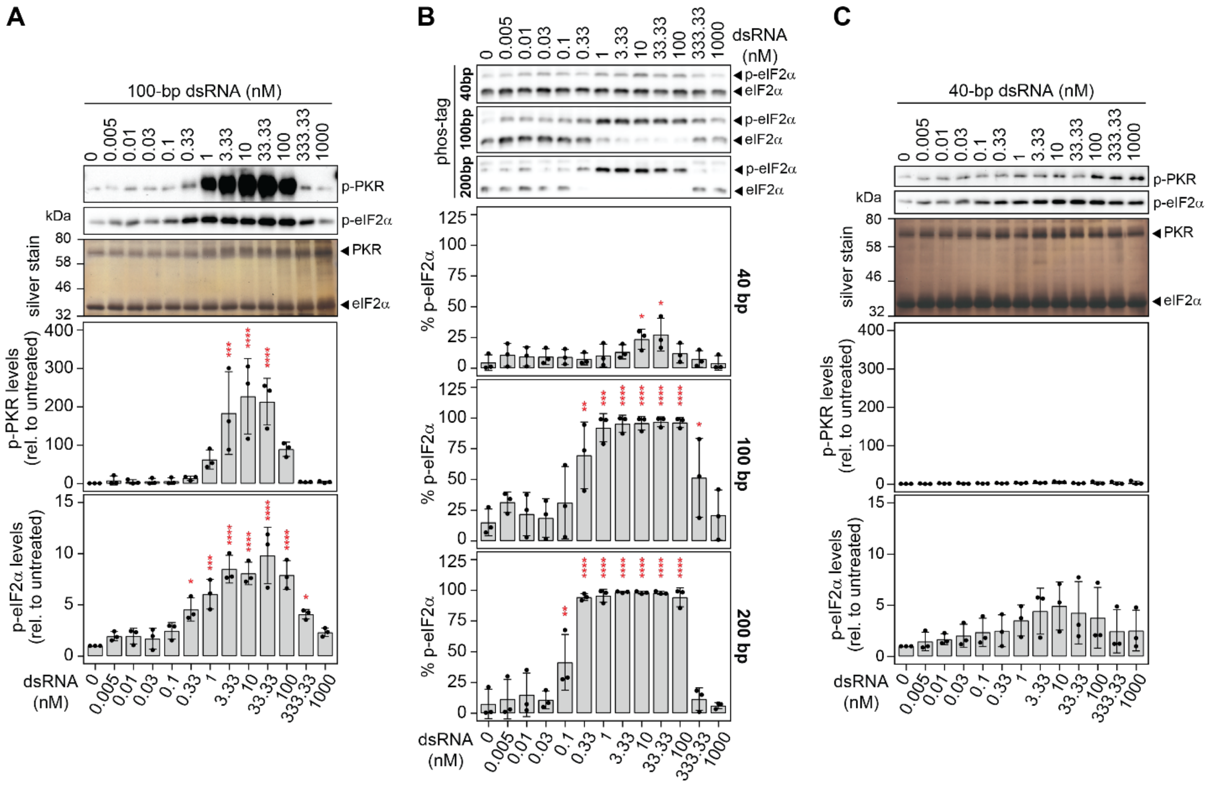


**Fig. S5. PKR *in vitro* kinase assay with dsRNA of varying lengths.** Increasing amounts of 40-bp, 100-bp and 200-bp dsRNA were incubated with recombinant His-tagged PKR and eIF2α (n=3). (**A**) Results for PKR *in vitro* activation by 100-bp dsRNA. Upper panel shows a representative Western blot analysis of p-PKR and p-eIF2α expression levels. Silver staining of the proteins on gel served as loading control. Quantification of mean levels ± SD are shown in the lower panels. Values were normalized to the untreated control. (**B**) Quantification of absolute p-eIF2α in PKR *in vitro* kinase assays. Upper panels show representative Phos-tag acrylamide gel analysis. Lower panels show quantification p-eIF2α. (**C**) Results for PKR *in vitro* activation by 40-bp dsRNA. Statistical significance is indicated compared to untreated; *p<0.05, **p<0.01, ***p<0.001, ****p<0.0001.

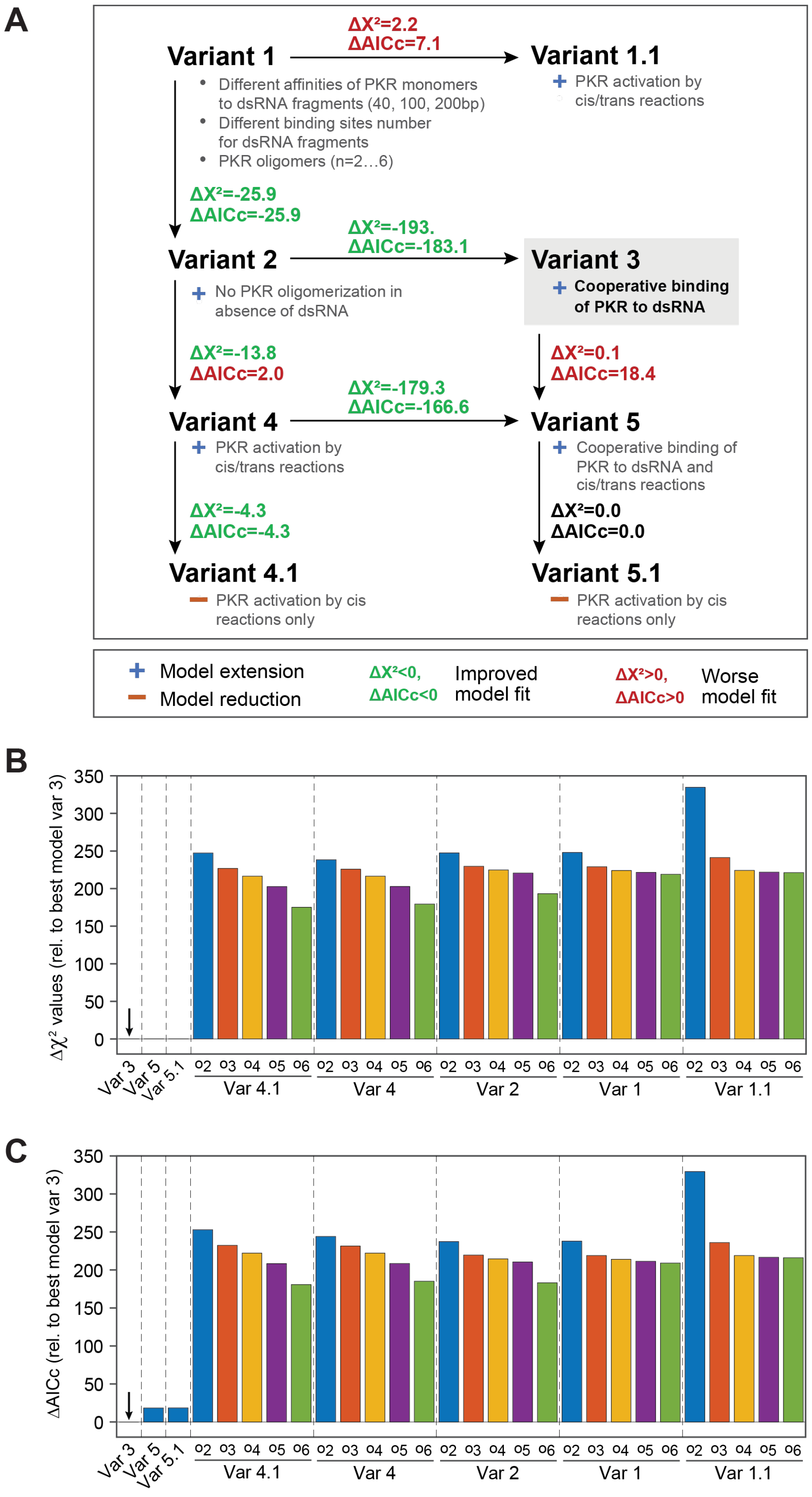


**Fig. S6. Selection of an optimal PKR activation model.** (**A**) The optimal PKR activation model variant was systematically identified by a sequence of model extensions (+) and reductions (–). Model features additionally included or excluded are described for each model variant. An improved fit was indicated by reduction in $\chi^2$ (**B**) or the corrected Akaike information criterion (**C**) ($\Delta\chi^2<0$, $\Delta$AICc<0), whereas, an increase in $\chi^2$ or AICc ($\Delta\chi^2>0$, $\Delta$AICc>0) indicated an inferior model fit. For model variants accounting for different PKR oligomerization states, differences are indicated for the best model variants. Based on AICc values, variant 3 was selected as an optimal model that accounts for (1) different affinities of PKR monomers to PKR bound at dsRNA fragments of 40-bp, 100-bp or 200-bp length, (2) different binding site numbers for dsRNA fragments, (3) no PKR oligomerization without dsRNA, and (4) cooperative binding of PKR at dsRNA. (**B**) Model variants 3, 5 and 5.1 ('Var3', 'Var 5', 'Var 5.1') accounting for cooperative binding of PKR to dsRNA were significantly better compared to variants 1 to 4.1 and variant 5 ($\Delta$X2>170). For model variants 1 to 4.1, different versions, in which PKR could form different types of oligomers (o2…o6: dimer to hexamer), were implemented and tested, resulting in a total number of 28 model versions. (**C**) Model selection based on differences in corrected Akaike information criterion ($\Delta$AICc). Variant 3 that did not account for *cis* and *trans* activation (Variant 5) or *cis* activation of PKR (Variant 5.1) was selected as the optimal model variant.

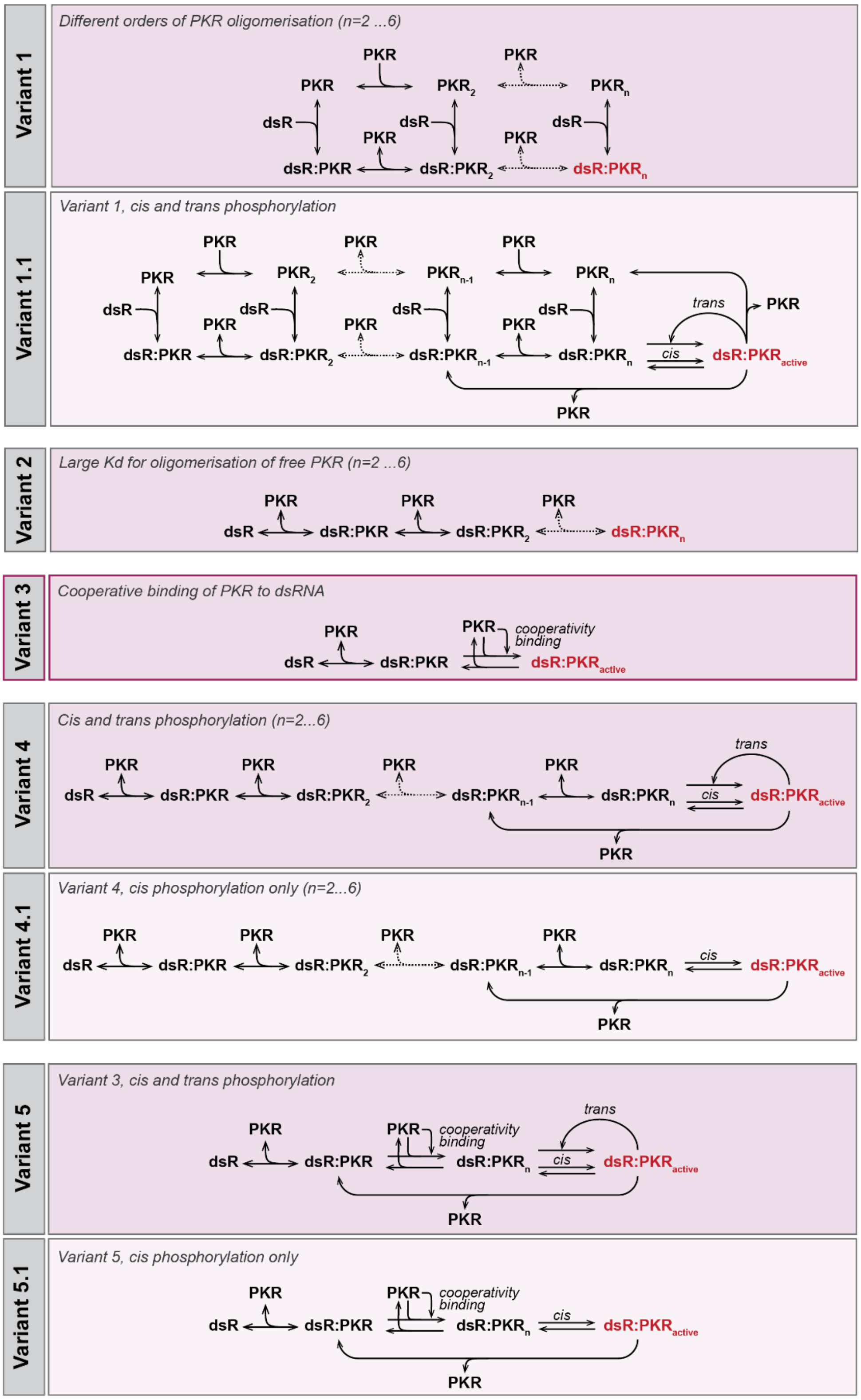


**Fig. S7. Visualizations of tested PKR activation model variants.** For model variants 1, 1.1, 2, 4 and 4.1, versions with different forms of PKR oligomerization (dimers to hexamers) were implemented and tested (dsR: binding sites for PKR at dsRNA; dsR:PKRn or dsR:PKRactive in red font: active PKR species). In these model versions that explicitly describe PKR oligomerization states, it was assumed that only the highest PKR oligomers could become active. In model variants 3, 5 and 5.1, PKR binding to PKR monomers at dsRNA was described as a cooperative process based on a Hill function. In model variants 1.1, 4, 4.1, 5 and 5.1, additional *cis* or *trans* reactions were included for describing PKR activation.

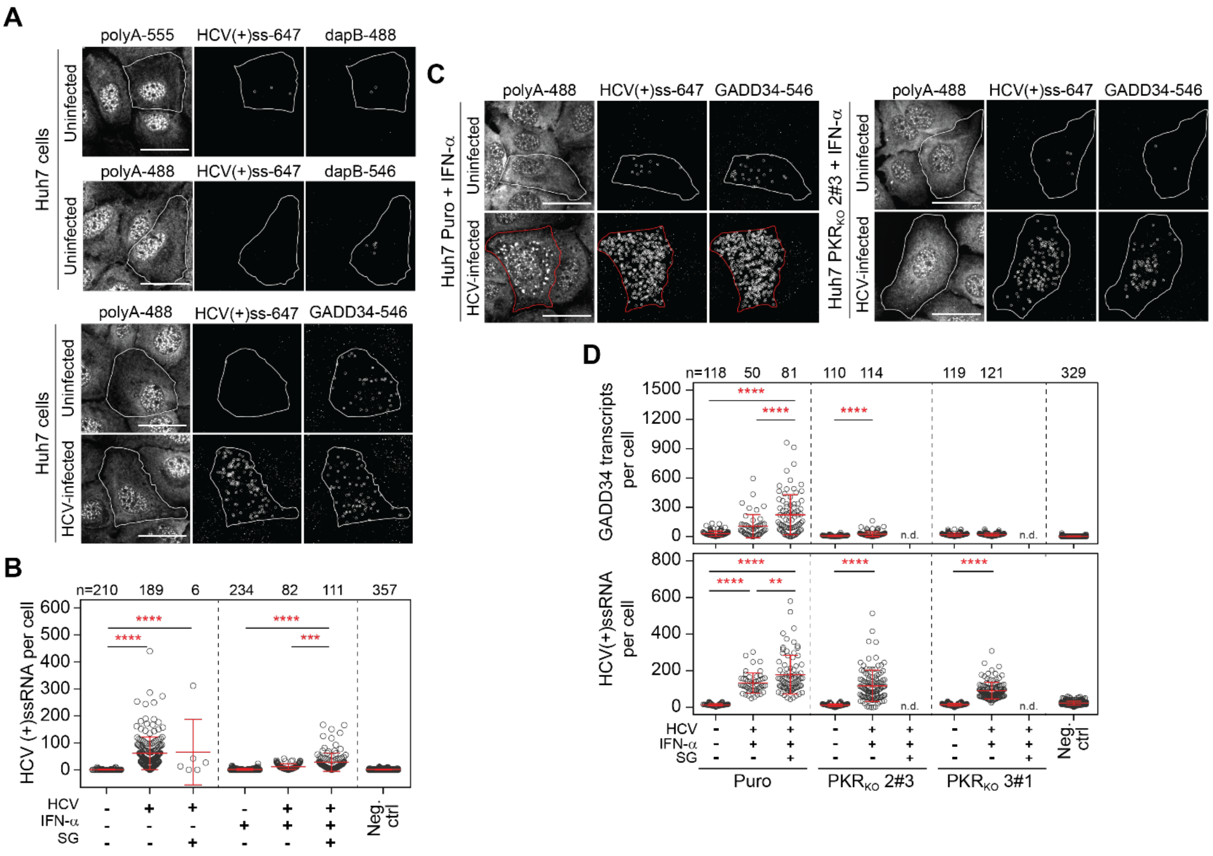


**Fig. S8. Transcriptional induction of GADD34 by HCV infection detected at single-cell level.** Huh7 cells were infected with HCV for 24 h and fixed for further FISH analysis. (**A**) Shown are representative still images of uninfected and HCV-infected cells. HCV (+) ssRNA genomes (HCV(+)ss), GADD34 transcripts and total polyadenylated mRNAs (polyA) were simultaneously detected using fluorescent probes. Additionally, uninfected cells were stained with probes targeting *E. coli* dapB transcripts and HCV (+)ss RNA genomes to determine background signals and unspecific binding of probes for the respective fluorophores. PolyA signal was used to outline cell boundaries. White circles indicate single particles detected after background subtraction. Scale bar, 20 µm. (**B**) Quantification of the mean number of HCV transcripts ± SD in Huh7 cells in presence and absence of IFN-α (related to Fig.5A). Statistical significance and the number of analyzed cells (n) cells from two independent biological repeats are indicated; ***p<0.001, ****p<0.0001. (**C and D**) FISH analysis of GADD34 transcriptional induction in PKR$_{KO}$ cells. Huh7 PKR$_{KO}$ Puro cells were left untreated or infected with HCV and treated with IFN-α. Huh7 Puro cells were used as a control. (C) Shown are representative still images. Scale bar, 20µm. (D) Quantification of mean GADD34 transcripts ± SD. Shown are pooled cells from two independent biological repeats. Statistical significance and the number of analyzed cells (n) cells are indicated; **p<0.01, ****p<0.0001.

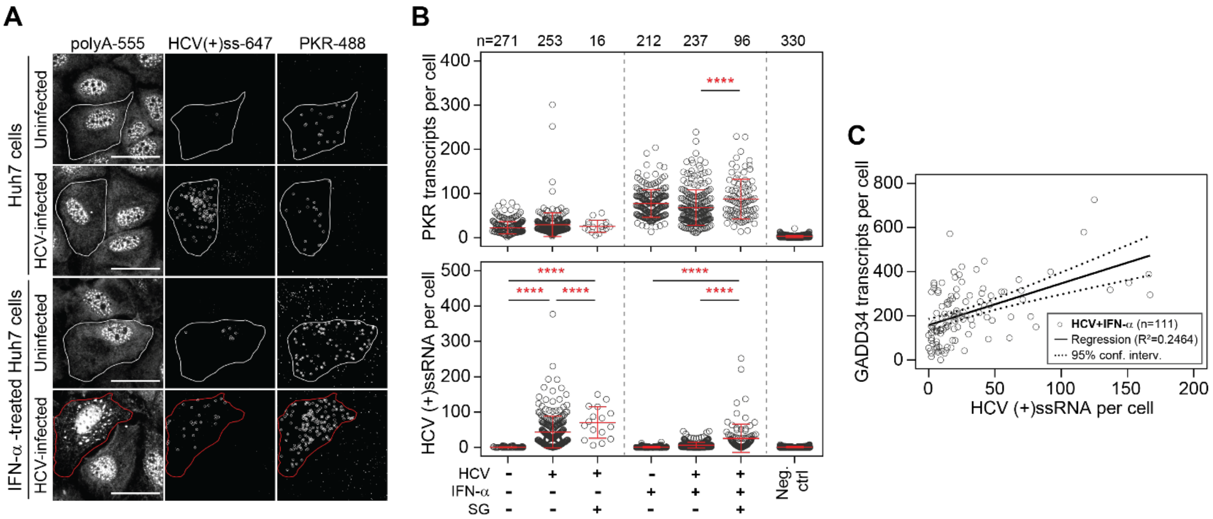


**Fig. S9. PKR transcriptional induction in HCV infected cells.** (**A and B**) Huh7 cells were infected with HCV for 24 h, subsequently treated with IFN-α for 24 h and fixed for further FISH analysis. (A) Shown are representative still images of uninfected and HCV-infected cells, left untreated (upper panel) or treated with IFN-α (bottom panel). HCV positive-sense (+) ssRNA genomes (HCV(+)ss), PKR transcripts and total polyA-tailed mRNAs (polyA) were simultaneously detected using fluorescent probes. PolyA signal was used to outline cell boundaries. White circles indicate single particles detected after background subtraction. Scale bar, 20 µm. (B) Quantification of the mean number of PKR (upper graph) and HCV (bottom graph) transcripts ± SD. Statistical significance and the number of analyzed cells (n) from two independent biological repeats are indicated; ****$p<0.0001$. (**C**) Scatter plot of the correlation between GADD34 transcripts and HCV (+)ssRNA genomes in HCV-infected cells treated with IFN-α. n, number of analyzed cells; $R^2$, coefficient of determination.

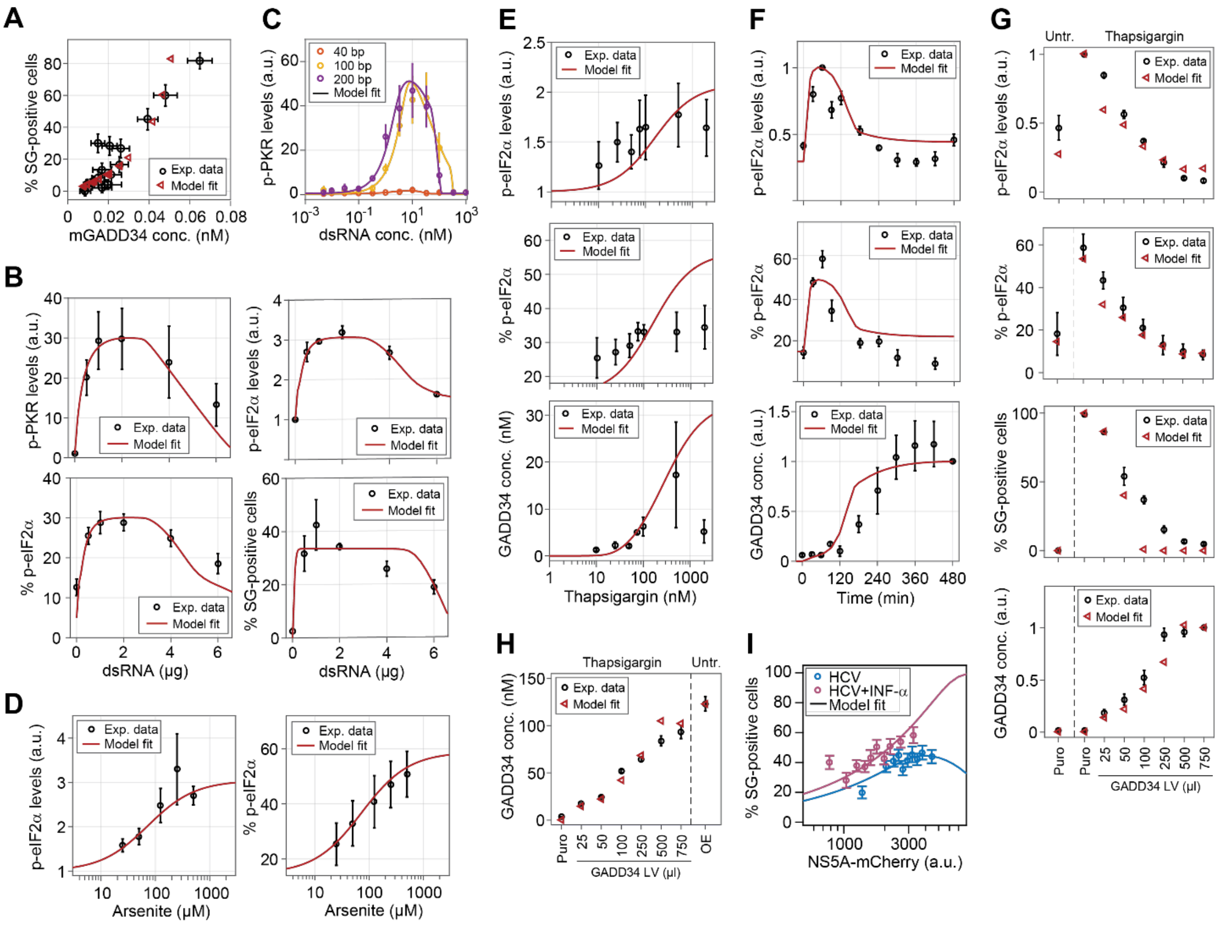


**Fig. S10. Best fits of the deterministic model of integrated stress response.** Multi-experiment fitting was performed by multi-starts local optimizations (n=2,500 runs). (**A**) Model fits to data from FISH analyses (mGADD34, mRNA of GADD34). Concentrations of GADD34 mRNA were estimated by dividing single-cell counts of detected particles by the average cell volume ($V_{cell}$=6.71pl). (**B**) Model fits to data from dsRNA transfection experiments (p-eIF2α in a. u. from Western blot analysis; % p-eIF2α from Phos-tag gel analysis; % SG-positive cells by fluorescence microscopy). (**C**) Model fits to data from *in vitro* PKR activation experiments using 40 bp, 100 bp or 200 bp dsRNA. (**D**) Model fits to data from arsenite titration experiments (the fit to % SG-positive cells is shown in Fig. 5D). (**E**) Model fits to data from thapsigargin titration experiments (the fit to % SG-positive cells shown in Fig. 5D). (**F**) Model fits to data from thapsigargin kinetics experiment. (**G** and **H**) Model fits to data from the GADD34 titration experiments. Cells were transduced with different amounts of lentivirus for GADD34 expression or a lentivirus for expression of a puromycin resistance gene as control (Puro) before treatment with thapsigargin (fits to p-eIF2α, % p-eIF2α, % SG-positive cells and GADD34 (F). (G) Fits to GADD34 concentration measurements in transduced cells compared to Huh7 GADD34 overexpressing cells (OE). (**I**) Model fits to the percentage of SG-positive cells as a function of NS5A-mCherry fluorescence intensity measured in time-lapse experiments.

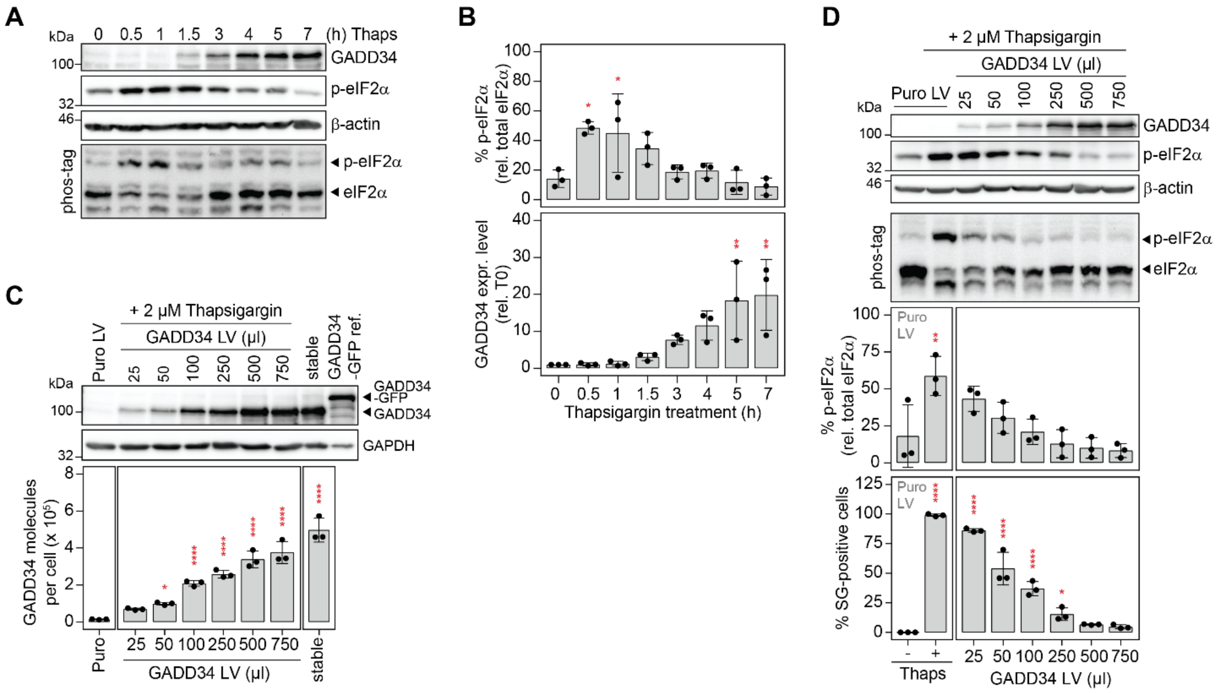


**Fig. S11. Characterization of GADD34 activity.** (**A and B**) Kinetics of GADD34 induction in Huh7 cells in response to treatment with 2 µM thapsigargin. (A) Shown are representative Western blot and Phos-tag gel analyses (n=3). (B) Quantification of mean percentage ± SD of p-eIF2α relative to total eIF2α (upper panel) and mean GADD34 expression levels ± SD normalized to the loading control β-actin and relative to untreated cells (lower panel). Statistical significance is indicated compared to untreated cells; *p<0.05, **p<0.01. (**C and D**) Ectopic expression of GADD34 using lentiviruses. Huh7 cells were transiently transduced by various amounts of lentivirus encoding GADD34 (GADD34 LV) for 30 hours. Lentivirus encoding for the puromycin resistance gene (Puro LV) was used as a control. (C) A reference lysate of Huh7 GADD34-GFP cells was loaded for quantification. Shown is a representative Western blot analysis of GADD34 expression. GAPDH was used as loading control (upper panel) (n=3). Quantification of the mean GADD34 molecules per cell ± SD is shown at the bottom. Statistical significance is indicated compared to cells transduced with control lentivirus (Puro); *p<0.05; ****p<0.0001. (D) Impact of GADD34 expression on eIF2α dephosphorylation (n=3). After transduction, Huh7 cells were treated with 2 µM thapsigargin for one hour and harvested for analysis of GADD34 and p-eIF2α by Western blot and Phos-tag gel analyses, respectively. SG formation was additionally analyzed in more than 100 cells for each condition by fluorescence microscopy. Shown in the top are representative Western blot and Phos-tag gel analyses. Quantifications of the mean percentages ± SD of p-eIF2α relative to total eIF2α and of SG-positive cells are shown at the bottom. Statistical significance is indicated compared untreated cells; *p<0.05; ****p<0.0001.

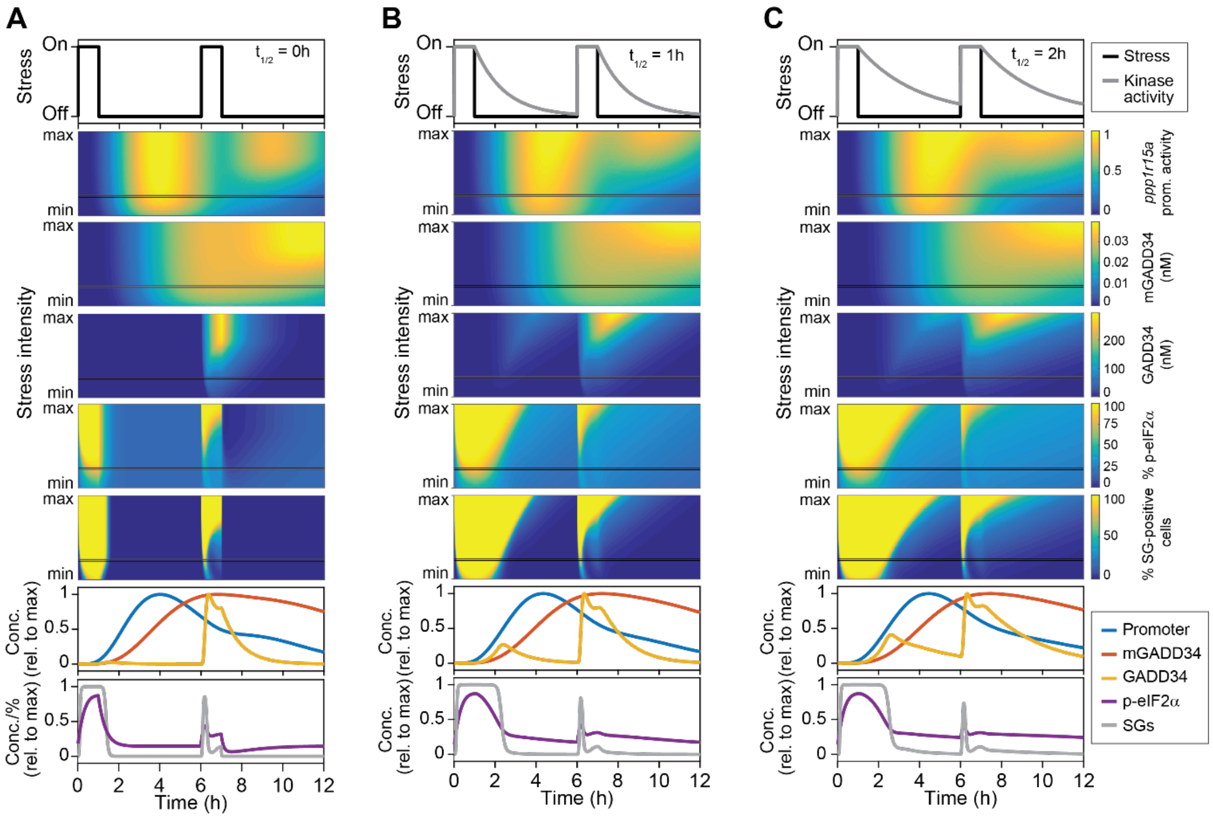


**Fig. S12. Adaptation to repeated stress pulses.** Computational simulations of two consecutive one-hour stress pulses interspaced by a five-hour recovery period. Shown is a range of stress kinase activities (stress intensities) varying between 10-fold lower (min) and 10-fold higher (max) than the reference kinase activity leading to 50% SG-positive cells. Color plots show the behavior of *ppp1r15a* promoter activity, concentrations of GADD34 mRNA and protein, percentages of p-eIF2α and SG-positive cells. Graphs at the bottom reflect the behavior of the above mentioned components for one chosen stress intensity (black line, moderate stress). Shown are the results for different kinase activity decays ($t_{1/2}$): (**A**) 0 h, (**B**) 1 h and (**C**) 2 h.

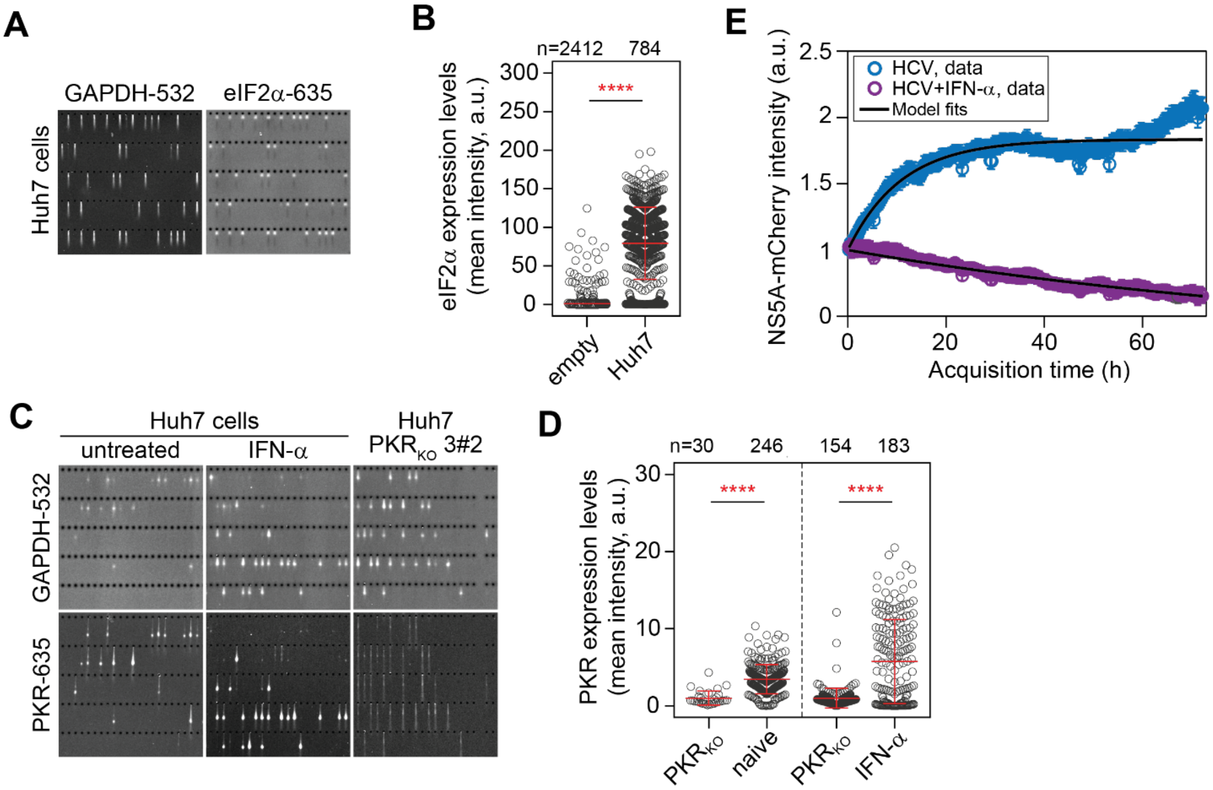


**Fig. S13. Single-cell Western blot analyses of PKR and eIF2α.** Quantification of the relative cell-to-cell variation in eIF2α and PKR expression levels by single-cell Western blot analysis. Single Huh7 cells were loaded into the micro-wells of array slides patterned in a photoactive polyacrylamide gel. After chemical lysis and electrophoresis, proteins were stained with primary and fluorescently-labeled secondary antibodies. The presence of cells in wells was confirmed by staining with an anti-GAPDH antibody. (**A and B**) Shown is a representative crop sections of a scanned micro-well array slide after staining. GADPH was detected using a secondary antibody labeled with AlexaFluor 532. Expression of eIF2α was detected using a secondary antibody labeled with AlexaFluor 635 (A). (B) Quantification of relative eIF2α levels per cell (a.u., arbitrary units). Mean signal intensity ± SD is indicated. The number of analyzed cells (n) and statistical significance compared to empty wells are indicated; ****$p<0.0001$. (**C and D**) Analysis of cell-to-cell variability of PKR expression levels in Huh7 cells left untreated or treated with IFN-α. Huh7 PKR$_{KO}$ cells were used as background control. (C) Representative crop sections of a scanned micro-well array slide after staining. GADPH was detected using a secondary antibody labeled with AlexaFluor 532. Expression of PKR was detected using a secondary antibody labeled with AlexaFluor 635. (D) Quantification of relative PKR levels per cell (a.u., arbitrary units). Mean signal intensity ± SD is indicated. The number of analyzed cells (n) and statistical significance compared to PKR$_{KO}$ cells are indicated; ****$p<0.0001$. (**E**) Analysis of NS5A-mCherry signal intensity in 72 h time-lapse experiments of HCV-infected cells in presence and absence of IFN-α. Shown are mean signal intensities (± SD) per cell for each time frame. Black lines represent best non-linear model fits.

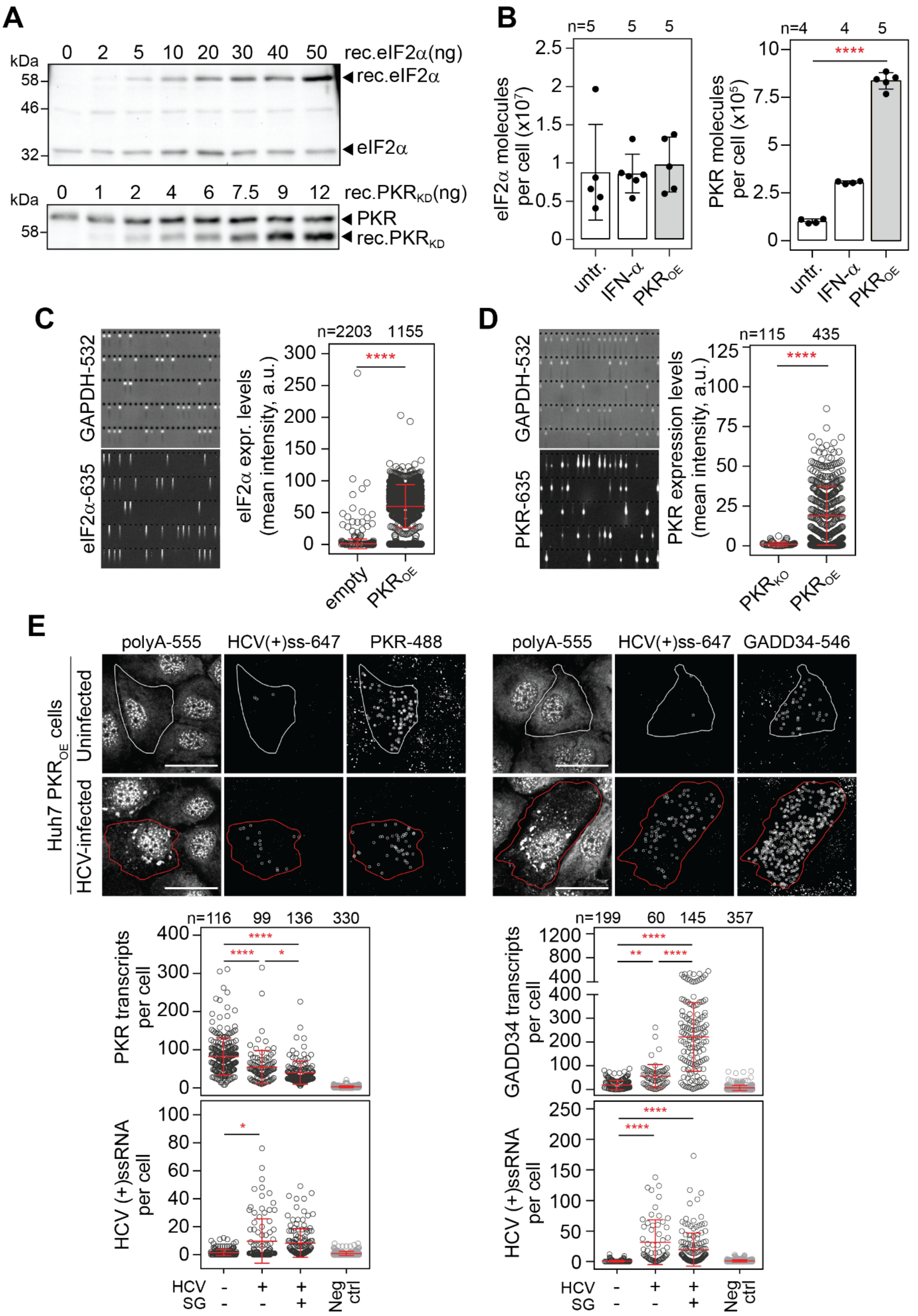


**Fig. S14. Analysis of PKR expression in Huh7 cells overexpressing PKR.** (**A and B**) Quantitative Western blot analysis of eIF2α and PKR expression levels in Huh7 $PKR_{OE}$ cells. Lysates of defined numbers of Huh7 $PKR_{OE}$ cells were spiked with increasing amounts of recombinant GST-tagged eIF2α (rec. eIF2α, top panel) or GST-tagged PKR kinase domain (rec. $PKR_{KD}$, bottom panel) and analyzed by quantitative Western blotting. (**B**) Estimated eIF2α and PKR mean molecule numbers per cell ± SD. Values for Huh7 cells untreated or treated with IFN-α (related to Fig. 2C) are shown for comparison. Number of repeats (n) and statistical significance compared to untreated cells are indicated; ****p<0.0001. (**C and D**) Cell-to-cell variability in eIF2α and PKR expression levels in Huh7 $PKR_{OE}$ cells. Empty wells or Huh7 $PKR_{KO}$ cells were used as background control. Shown are representative crop sections of a scanned micro-well array slide after staining. GADPH was detected using a secondary antibody labeled with AlexaFluor 532. Expression of eIF2α or PKR was detected using a secondary antibody labeled with AlexaFluor 635. (C) Quantification of relative eIF2α levels per cell (a.u., arbitrary units). Mean signal intensity ± SD is indicated. The number of analyzed cells (n) and statistical significance compared to empty wells are indicated; ****p<0.0001. (D) Quantification of PKR levels per cell (a.u., arbitrary units). Mean signal intensity ± SD is indicated. The number of analyzed cells (n) and statistical significance compared to $PKR_{KO}$ cells are indicated; ****p<0.0001. (**E**) Analysis of PKR and GADD34 transcription levels in HCV-infected Huh7 $PKR_{OE}$ cells by FISH. The upper panel shows representative still images of uninfected and HCV-infected cells. HCV (+) ssRNA genomes, GADD34 transcripts and total polyadenylated mRNAs (polyA) were simultaneously detected using fluorescent probes. Staining of polyA mRNAs allowed for the visualization of SGs and cell boundaries. Outlined in red is a SG-positive cell, outlined in white an unstressed cell. White circles indicate single particles detected after background subtraction. Scale bar, 20 µm. Scatter plots of GADD34 mean transcript levels ± SD per cell are shown on the bottom. Statistical significance and the number of analyzed cells (n) cells are indicated; *p<0.05, **p<0.01, ****p<0.0001.

**Table S1. Selection of PKR activation models visualized in Fig. S6.**

| | |
|---|---|
| **Variant 1** (Supplementary Fig. S6; n=2, oligomerization of PKR up to dimers) | |
| Model equations | Description |
| $\frac{d[PKR]}{dt} = -k_{p,on,i}[PKR]^2 + k_{p,off}[PKR_2] - k_{d,on,i}[PKR][dsR] + k_{d,off}[dsR{:}PKR] - k_{p,on,i}[PKR][dsR{:}PKR] + k_{p,off}[dsR{:}PKR_2]$ <br> with $k_{d,on,i} = \begin{cases} k_{d,on} & for \quad i = 1 \quad (40bp\text{,in-vitro}) \\ k_{d,on}\alpha_{100bp} & for \quad i = 2 \quad (100bp\text{,in-vitro}) \\ k_{d,on}\alpha_{200bp} & for \quad i \in \{3,4\} \quad (200bp\text{,in-vitro/transf.}) \end{cases}$ <br> and $k_{p,on,i} = \begin{cases} k_{p,on} & for \quad i = 1 \quad (40bp\text{,in-vitro}) \\ k_{p,on}\gamma_{100bp} & for \quad i = 2 \quad (100bp\text{,in-vitro}) \\ k_{p,on}\gamma_{200bp} & for \quad i \in \{3,4\} \quad (200bp\text{,in-vitro/transf.}) \end{cases}$ | Unbound PKR |
| $\frac{d[dsR]}{dt} = -k_{d,on,i}[PKR][dsR] + k_{d,off}[dsR{:}PKR] - k_{d,on,i}[PKR_2][dsR] + k_{d,off}[dsR{:}PKR_2]$ | Binding to dsRNA and unbinding |
| $\frac{d[dsR{:}PKR]}{dt} = k_{d,on,i}[PKR][dsR] - k_{d,off}[dsR{:}PKR] - k_{p,on,i}[PKR][dsR{:}PKR] + k_{p,off}[dsR{:}PKR_2]$ | Complexes dsRNA and PKR |
| $\frac{d[PKR_2]}{dt} = k_{p,on,i}[PKR]^2 - k_{p,off}[PKR_2] - k_{d,on,i}[PKR_2][dsR] + k_{d,off}[dsR{:}PKR_2]$ | PKR dimers |
| $\frac{d[dsR{:}PKR_2]}{dt} = k_{d,on,i}[PKR_2][dsR] - k_{d,off}[dsR{:}PKR_2] + k_{p,on,i}[PKR][dsR{:}PKR] - k_{p,off}[dsR{:}PKR_2]$ | Active PKR dimers |
| **Variant 3 (optimal model variant)** | |
| Model equations | Description |
| $\frac{d[PKR]}{dt} = -k_{d,on,i}[PKR][dsR] + k_{d,off}[dsR{:}PKR] - k_{p,on}[dsR{:}PKR][PKR]\frac{[PKR]^h}{K_{p,on,i}^h + [PKR]^h} + k_{p,off}[dsR{:}PKR_{olig}]$ <br> with $k_{d,on,i} = \begin{cases} k_{d,on} & for \quad i = 1 \quad (40bp\text{,in-vitro}) \\ k_{d,on}\alpha_{100bp} & for \quad i = 2 \quad (100bp\text{,in-vitro}) \\ k_{d,on}\alpha_{200bp} & for \quad i \in \{3,4\} \quad (200bp\text{,in-vitro/transf.}) \end{cases}$ <br> and $K_{p,on,i} = \begin{cases} K_{p,on} & for \quad i = 1 \quad (40bp\text{,in-vitro}) \\ K_{p,on}\gamma_{200bp} & for \quad i = 2 \quad (100bp\text{,in-vitro}) \\ K_{p,on}\gamma_{200bp} & for \quad i \in \{3,4\} \quad (200bp\text{,in-vitro/transf.}) \end{cases}$ | Binding of PKR to dsRNA and PKR oligomerization at dsRNA, different affinities for binding to dsRNA and oligomerization of dsRNA fragments |
| $\frac{d[dsR]}{dt} = -k_{d,on,i}[PKR][dsR] + k_{d,off}[dsR{:}PKR]$ | Binding to dsRNA and unbinding |

| | |
|---|---|
| $\frac{d[dsR:PKR]}{dt} = k_{d,on,i}[PKR][dsR] - k_{d,off}[dsR:PKR]$ <br> $-k_{p,on}[dsR:PKR][PKR]\frac{[PKR]^h}{K_{p,on,i}^h + [PKR]^h} + k_{p,off}[dsR:PKR_{olig}]$ | Complexes of dsRNA and PKR |
| $\frac{d[dsR:PKR_{olig}]}{dt} = k_{p,on}[dsR:PKR][PKR]\frac{[PKR]^h}{K_{p,on,i}^h + [PKR]^h}$ <br> $-k_{p,off}[dsR:PKR_{olig}]$ | Active oligomerized PKR bound to dsRNA |
| Start value assignments, observables, algebraic equations, scaling factors, background offsets* | Description |
| $[\mathrm{dsR}](t_0)_{40bp} = [dsRNA_{40bp}]$ <br> $[\mathrm{dsR}](t_0)_{100bp} = [dsRNA_{100bp}]f_{sites,100bp}$ <br> $[\mathrm{dsR}](t_0)_{200bp} = [dsRNA_{200bp}]f_{sites,200bp}$ <br> $[\mathrm{dsR}](t_0)_{200bp,transf} = [dsRNA_{200bp}]f_{sites,200bp}\beta_{transf}$ | Start value assignment of PKR binding sites at dsRNA (dsR) depending on the concentration of dsRNA fragments, factors for 100bp and 200bp fragments and a factor for transfection |
| $y_{PKR*,40bp} = s_{PKR*,iv}[dsR:PKR_{olig}] + b_{PKR*,iv,40bp}$ <br> $y_{PKR*,100bp} = s_{PKR*,iv}[dsR:PKR_{olig}] + b_{PKR*,iv,100bp}$ <br> $y_{PKR*,200bp} = s_{PKR*,iv}[dsR:PKR_{olig}] + b_{PKR*,iv,200bp}$ <br> $y_{PKR*,200bp,transf} = s_{PKR*,transf}[dsR:PKR_{olig}] + b_{PKR*,transf}$ | Observables for phosphorylated PKR at dsRNA fragments containing scaling factors and background intensities |

* Upper limits of background offsets were defined by minimal values of respective datasets (Supplementary Table S4).

**Table S2. Deterministic model of the integrated stress response visualized in Fig. 2A.**

| Model equations | Description |
|---|---|
| $\frac{d[PKR]}{dt} = -k_{d,on,i}[PKR][dsR] + k_{d,off}[dsR{:}PKR]$ <br> $-k_{p,on}[dsR{:}PKR][PKR]\frac{[PKR]^h}{K_{p,on,i}^h+[PKR]^h} + k_{p,off}[dsR{:}PKR_{olig}]$ <br> (1) <br> with <br> $k_{d,on,i} = \begin{cases} k_{d,on} & for \quad i=1 \quad (40bp\text{,in-vitro}) \\ k_{d,on}\alpha_{100bp} & for \quad i=2 \quad (100bp\text{,in-vitro}) \\ k_{d,on}\alpha_{200bp} & for \quad i\in\{3,4\} \quad (200bp\text{,in-vitro/transf.}) \\ k_{d,on}\alpha_{HCV} & for \quad i=5 \quad (HCV) \end{cases}$ <br> and <br> $K_{p,on,i} = \begin{cases} K_{p,on} & for \quad i=1 \quad (40bp\text{,in-vitro}) \\ K_{p,on}\gamma_{100bp} & for \quad i=2 \quad (100bp\text{,in-vitro}) \\ K_{p,on}\gamma_{200bp} & for \quad i\in\{3,4\} \quad (200bp\text{,in-vitro/transf.}) \\ K_{p,on}\gamma_{HCV} & for \quad i=5 \quad (HCV) \end{cases}$ | Binding of PKR to dsRNA and PKR oligomerization at dsRNA, different affinities for binding to dsRNA and oligomerization of dsRNA fragments and HCV |
| $\frac{d[dsR]}{dt} = -k_{d,on,i}[PKR][dsR] + k_{d,off}[dsR{:}PKR]$ (2) | Binding to dsRNA and unbinding |
| $\frac{d[dsR{:}PKR]}{dt} = k_{d,on,i}[PKR][dsR] - k_{d,off}[dsR{:}PKR]$ <br> $-k_{p,on}[dsR{:}PKR][PKR]\frac{[PKR]^h}{K_{p,on,i}^h+[PKR]^h} + k_{p,off}[dsR{:}PKR_{olig}]$ (3) | Complexes of dsRNA and PKR |
| $\frac{d[dsR{:}PKR_{olig}]}{dt} = k_{p,on}[dsR{:}PKR][PKR]\frac{[PKR]^h}{K_{p,on,i}^h+[PKR]^h} - k_{p,off}[dsR{:}PKR_{olig}]$ <br> (4) | Active oligomerized PKR bound to dsRNA |
| $\frac{d[eIF2\alpha]}{dt} = -k_{ph,basal}[eIF2\alpha] - k_{ph,PKR}[dsR{:}PKR_{olig}]\frac{[eIF2\alpha]}{K_{ph,PKR}+[eIF2\alpha]}$ <br> $+k_{CReP}[eIF2\alpha *] + k_{deph}[eIF2\alpha *][GADD34]$ (5) | Phosphorylation of eIF2α by PKR |
| $\frac{d[eIF2\alpha *]}{dt} = k_{ph,basal}[eIF2\alpha] + k_{ph,PKR}[dsR{:}PKR_{olig}]\frac{[eIF2\alpha]}{K_{ph,PKR}+[eIF2\alpha]}$ <br> $-k_{CReP}[eIF2\alpha *] - k_{deph}[eIF2\alpha *][GADD34]$ (6) <br> $f_{SG,PKR} = \frac{[eIF2\alpha *]^l}{K_{SG,PKR}^l + [eIF2\alpha *]^l}$ | Phosphorylated eIF2α, PKR; $f_{SG,PKR}$, function describing SG formation |
| $\frac{d[eIF2\alpha]}{dt} = -k_{ph,basal}[eIF2\alpha] - k_{ph,A}\frac{[A]}{K_A+[A]}\frac{[eIF2\alpha]}{K_{ph,A}+[eIF2\alpha]}$ <br> $+k_{CReP}[eIF2\alpha *] + k_{deph}[eIF2\alpha *][GADD34]$ (7) | Phosphorylation of eIF2α by HRI after activation of HRI by arsenite (A, arsenite) |
| $\frac{d[eIF2\alpha *]}{dt} = k_{ph,basal}[eIF2\alpha] + k_{ph,A}\frac{[A]}{K_A+[A]}\frac{[eIF2\alpha]}{K_{ph,A}+[eIF2\alpha]}$ <br> $-k_{CReP}[eIF2\alpha *] - k_{deph}[eIF2\alpha *][GADD34]$ (8) | Phosphorylated eIF2α, HRI |

| | |
|---|---|
| $$f_{SG,A} = \frac{[eIF2\alpha *]^l}{K_{SG,A}^l + [eIF2\alpha *]^l}$$ | |
| $$\frac{d[eIF2\alpha]}{dt} = -k_{ph,basal}[eIF2\alpha] - k_{ph,T}\frac{[T]}{K_T + [T]}\frac{[eIF2\alpha]}{K_{ph,T} + [eIF2\alpha]}$$ $$+k_{CReP}[eIF2\alpha *] + k_{deph}[eIF2\alpha *][GADD34] \quad (9)$$ | Phosphorylation of eIF2α by PERK after activation of PERK by thapsigargin (T, thapsigargin) |
| $$\frac{d[eIF2\alpha *]}{dt} = k_{ph,basal}[eIF2\alpha] + k_{ph,T}\frac{[T]}{K_T + [T]}\frac{[eIF2\alpha]}{K_{ph,T} + [eIF2\alpha]}$$ $$-k_{CReP}[eIF2\alpha *] - k_{deph}[eIF2\alpha *][GADD34] \quad (10)$$ $$f_{SG,T} = \frac{[eIF2\alpha *]^l}{K_{SG,T}^l + [eIF2\alpha *]^l}$$ | Phosphorylated eIF2α, PERK |
| $$\frac{d[Pr_{off}]}{dt} = -k_{Pr,on}[Pr_{off}]f_{SG}([eIF2\alpha *]) + k_{Pr,off}[Pr_{on}] \quad (11)$$ | PPP1R15A promoter activation |
| $$\frac{d[Pr_{delay,1}]}{dt} = k_{Pr,on}[Pr_{off}]f_{SG}([eIF2\alpha *]) - k_{Pr,on}[Pr_{delay,1}] \quad (12)$$ $$\frac{d[Pr_{delay,2}]}{dt} = k_{Pr,on}[Pr_{delay,1}] - k_{Pr,on}[Pr_{delay,2}] \quad (13)$$ $$\frac{d[Pr_{delay,3}]}{dt} = k_{Pr,on}[Pr_{delay,2}] - k_{Pr,on}[Pr_{delay,3}] \quad (14)$$ $$\frac{d[Pr_{delay,4}]}{dt} = k_{Pr,on}[Pr_{delay,3}] - k_{Pr,on}[Pr_{delay,4}] \quad (15)$$ $$\frac{d[Pr_{on}]}{dt} = k_{Pr,on}[Pr_{delay,4}] - k_{Pr,off}[Pr_{on}] \quad (16)$$ | Delay of transcriptional response described by reaction chain |
| $$\frac{d[mGADD34]}{dt} = k_{syn,m}[Pr_{on}] - k_{deg,m}[mGADD34] \quad (17)$$ | Turnover of GADD34 mRNA |
| $$\frac{d[GADD34]}{dt} = k_{syn,GADD34}[mGADD34]f_{SG}([eIF2\alpha *])$$ $$-k_{deg,GADD34}[GADD34]$$ (18) | Turnover of GADD34 |

741

**Table S3. Parts of deterministic model and model observables.**

| Model part | Start value assignments, observables, algebraic equations, scaling factors, background offsets* |
|---|---|
| **PKR activation**, in-vitro assay (40bp, 100bp and 200bp dsRNA) | $[\mathrm{dsR}](t_0)_{40bp} = [dsRNA_{40bp}]$ (1)<br>$[\mathrm{dsR}](t_0)_{100bp} = [dsRNA_{100bp}]f_{sites,100bp}$ (2)<br>$[\mathrm{dsR}](t_0)_{200bp} = [dsRNA_{200bp}]f_{sites,200bp}$ (3)<br>$y_{PKR*,40bp} = s_{PKR*,iv}[dsR:PKR_{olig}] + b_{PKR*,iv,40bp}$ (4)<br>$y_{PKR*,100bp} = s_{PKR*,iv}[dsR:PKR_{olig}] + b_{PKR*,iv,100bp}$ (5)<br>$y_{PKR*,200bp} = s_{PKR*,iv}[dsR:PKR_{olig}] + b_{PKR*,iv,200bp}$ (6) |
| **PKR activation**, transfection with 200bp dsRNA | $[\mathrm{dsR}](t_0)_{200bp,transf} = [dsRNA_{200bp}]f_{sites,200bp}\beta_{transf}$ (7)<br>$y_{PKR*} = s_{PKR*,transf}[dsR:PKR_{olig}] + b_{PKR*,transf}$ (8)<br>$y_{eIF2\alpha*} = s_{eIF2\alpha*,dsRNA}[eIF2\alpha *] + b_{eIF2\alpha*,dsRNA}$ (9)<br>$y_{phostag} = \varepsilon_{transf}\frac{[eIF2\alpha *]}{[eIF2\alpha] + [eIF2\alpha *]}$ (10)<br>$y_{SG} = \varepsilon_{transf}f_{SG,PKR}$ (11)<br>$f_{SG,PKR} = \frac{[eIF2\alpha *]^l}{K_{SG,PKR}^l + [eIF2\alpha *]^l}$ (12) |
| **Arsenite titration** | $y_{eIF2\alpha*} = s_{eIF2\alpha*,A}[eIF2\alpha *] + b_{eIF2\alpha*,A}$ (13)<br>$y_{phostag} = \frac{[eIF2\alpha *]}{[eIF2\alpha] + [eIF2\alpha *]}$ (14)<br>$y_{SG} = f_{SG,A}$ (15)<br>$f_{SG,A} = \frac{[eIF2\alpha *]^l}{K_{SG,A}^l + [eIF2\alpha *]^l}$ (16) |
| **Thapsigargin titration** | $y_{eIF2\alpha*} = s_{eIF2\alpha*,T}[eIF2\alpha *] + b_{eIF2\alpha*,T}$ (17) |

| | | |
|---|---|---|
| | $$y_{phostag} = \frac{[eIF2\alpha *]}{[eIF2\alpha] + [eIF2\alpha *]}$$ | (18) |
| | $$y_{GADD34,quant} = [GADD34]$$ | |
| | $$y_{SG} = f_{SG,T}$$ | (19) |
| | $$f_{SG,T} = \frac{[eIF2\alpha *]^l}{K_{SG,T}^l + [eIF2\alpha *]^l}$$ | (20) |
| | | (21) |
| **Thapsigargin kinetics** | $$y_{eIF2\alpha *} = s_{eIF2\alpha *,T,kin}[eIF2\alpha *] + b_{eIF2\alpha *,T,kin}$$ | (22) |
| | $$y_{phostag} = \frac{[eIF2\alpha *]}{[eIF2\alpha] + [eIF2\alpha *]}$$ | (23) |
| | $$y_{GADD34} = s_{GADD34,T,kin}[GADD34]$$ | (24) |
| **GADD34 titration** (thapsigargin treatment in GADD34-overexpressing and control cells) | $$[\mathrm{GADD34}_{oe,i}](t_0) = \begin{cases} 0 & for \quad i = 0 \\ \mathrm{GADD34}_{ex,i} & for \quad i \in \{1, \dots, 7\} \end{cases}$$ | (25) |
| | $$y_{eIF2\alpha *} = s_{eIF2\alpha *,G}[eIF2\alpha *] + b_{eIF2\alpha *,G}$$ | (26) |
| | $$y_{phostag} = \frac{[eIF2\alpha *]}{[eIF2\alpha] + [eIF2\alpha *]}$$ | (27) |
| | $$y_{GADD34,i} = s_{GADD34,G} \begin{cases} [GADD34] & for \quad i = 0 \\ [\mathrm{GADD34}_{ex}] & for \quad i \in \{1, \dots, 7\} \end{cases}$$ | (28) |
| | $$y_{GADD34,quant,i} = \begin{cases} [GADD34] & for \quad i = 0 \\ [\mathrm{GADD34}_{ex}] & for \quad i \in \{1, \dots, 7\} \end{cases}$$ | |
| | $$y_{SG} = f_{SG,T}$$ | (29) |
| | $$f_{SG,T} = \frac{[eIF2\alpha *]^l}{K_{SG,T}^l + [eIF2\alpha *]^l}$$ | (30) |
| | | (31) |
| **FISH** (Cells infected with HCV) | $$[\mathrm{PKR} *_{act,i}](t_0) = PKR\_act_i \quad for \quad i \in \{1, \dots, 19\}$$ | (32) |
| | $$y_{GADD34,mRNA} = [mGADD34]$$ | (33) |
| | $$y_{SG} = f_{SG,PKR}$$ | (34) |
| | $$f_{SG,PKR} = \frac{[eIF2\alpha *]^l}{K_{SG,PKR}^l + [eIF2\alpha *]^l}$$ | |

| | |
|---|---|
| | (35) |
| **Infection marker** (Cells infected with HCV; $I_{inf}$, infection marker intensity) | $[\mathrm{dsR}](t_0)_{HCV} = I_{inf} f_{sites,HCV}$ (36)<br><br>$y_{SG} = f_{SG,PKR}$ (37)<br><br>$f_{SG,PKR} = \frac{[eIF2\alpha *]^l}{K_{SG,PKR}^l + [eIF2\alpha *]^l}$ (38) |

* Upper limits of background offsets were defined by minimal values of respective datasets (Supplementary Table S4).

**Table S4. Parameter estimates and confidence intervals.**

| Parameter | Unit | Best fit value | Confidence interval obtained from profile likelihood estimation | | identifiable | Allowed parameter interval | |
|---|---|---|---|---|---|---|---|
| | | | Lower bound | Upper bound | | Lower bound | Upper bound |
| $k_{p,on}/k_{p,off}$* | $nM^{-1}$ | 0.09159 | 0.07153 | 0.1228 | yes | $10^{-6}$ | $10^{5}$ |
| $k_{d,on}/k_{d,off}$* | $nM^{-1}$ | 0.002426 | 0.002203 | 0.002954 | yes | $10^{-6}$ | $10^{5}$ |
| $\alpha_{100bp}$ | unitless | 17.63 | 8.339 | 34.45 | yes | 1 | $10^{3}$ |
| $\alpha_{200bp}$ | unitless | 3.881 | 1.951 | 6.66 | yes | 1 | $10^{3}$ |
| $\alpha_{HCV}$ | unitless | 19.14 | 10.48 | 41.79 | yes | $10^{-6}$ | $10^{3}$ |
| $K_{p,on}$ | nM | 147.4 | 146.8 | 148.1 | yes | $10^{-6}$ | $10^{5}$ |
| $h$ | unitless | 100 | 85.83 | Inf | no | 1 | $10^{2}$ |
| $\gamma_{100bp}$ | unitless | 0.006702 | 0.003701 | 0.007295 | yes | $10^{-6}$ | 1 |
| $\gamma_{200bp}$ | unitless | 0.04226 | 0.02792 | 0.07227 | yes | $10^{-6}$ | 1 |
| $\gamma_{HCV}$ | unitless | 16.98 | 0 | Inf | no | $10^{-6}$ | $10^{3}$ |
| $f_{sites,100bp}$ | unitless | 11.04 | 9.421 | 13.07 | yes | 1 | $10^{3}$ |
| $f_{sites,200bp}$ | unitless | 23.22 | 20.69 | 26.47 | yes | 1 | $10^{3}$ |
| $\beta_{transf}$ | unitless | 2.278 | 1.857 | 2.519 | yes | $10^{-6}$ | $10^{5}$ |
| $f_{sites,HCV}$ | nM | 0.0004136 | 0.0002634 | 0.0005161 | yes | $10^{-6}$ | $10^{3}$ |
| $k_{ph,basal}$ | $min^{-1}$ | 0.006799 | 0.005629 | 0.008403 | yes | $10^{-6}$ | $10^{5}$ |
| $k_{CReP}$ | $min^{-1}$ | 0.0395 | 0.03314 | 0.04858 | yes | $10^{-6}$ | $10^{5}$ |
| $k_{ph,PKR}$ | $min^{-1}$ | 1020 | 275.1 | 543.9 | yes | $10^{-6}$ | $10^{5}$ |
| $K_{ph,PKR}$ | nM | 60.52 | 0 | Inf | no | $10^{-6}$ | $10^{5}$ |
| $k_{deph}$ | $min^{-1}$ | 0.003571 | 0.003087 | 0.004288 | yes | $10^{-6}$ | $10^{5}$ |
| $K_{SG,PKR}$ | nM | 841.8 | 687.5 | 883.3 | yes | $10^{-6}$ | $10^{5}$ |
| $l$ | unitless | 11.25 | 9.911 | 12.34 | yes | 1 | 50 |
| $k_{ph,A}$ | $nM\cdot min^{-1}$ | 45.94 | 37.24 | 57.83 | yes | $10^{-6}$ | $10^{5}$ |
| $K_{ph,PKR}$ | nM | 0.003542 | 0 | Inf | no | $10^{-6}$ | $10^{5}$ |
| $K_{A}$ | nM | 72.71 | 61.79 | 88.17 | yes | $10^{-6}$ | $10^{5}$ |
| $K_{SG,A}$ | nM | 892.8 | 819 | 969.6 | yes | $10^{-6}$ | $10^{5}$ |
| $k_{ph,T}$ | $nM\cdot min^{-1}$ | 275.5 | 168.8 | Inf | no | $10^{-6}$ | $10^{5}$ |
| $K_{ph,T}$ | nM | 5195 | 2639 | Inf | no | $10^{-6}$ | $10^{5}$ |
| $K_{T}$ | nM | 275.9 | 250.7 | 305.6 | yes | $10^{-6}$ | $10^{5}$ |
| $K_{SG,T}$ | nM | 605.4 | 588.0 | 622.6 | yes | $10^{-6}$ | $10^{5}$ |
| $k_{Pr,on}$ | $min^{-1}$ | 0.01499 | 0.01339 | 0.01701 | yes | $10^{-6}$ | $10^{5}$ |
| $k_{Pr,off}$ | $min^{-1}$ | 5619 | 0 | Inf | no | $10^{-6}$ | $10^{5}$ |
| $k_{syn,m}$ | $nM\cdot min^{-1}$ | 345.6 | 0 | Inf | no | $10^{-6}$ | $10^{5}$ |
| $k_{deg,m}$ | $min^{-1}$ | 0.0035** | | | | | |
| $k_{syn,GADD34}$ | $min^{-1}$ | 200.2 | 137.2 | 285.4 | yes | $10^{-6}$ | $10^{5}$ |
| $k_{deg,GADD34}$ | $min^{-1}$ | 0.01854*** | 0.01516 | 0.02289 | yes | $10^{-4}$ | $10^{2}$ |
| $s_{PKR*,transf}$ | $nM^{-1}$ | 5.652 | 4.749 | 7.252 | yes | $10^{-4}$ | $10^{6}$ |
| $b_{PKR*,transf}$ | unitless | 1.000 | 0.9795 | 1.019 | yes | $10^{-4}$ | 1.11 |
| $\varepsilon_{transf}$ | unitless | 0.336 | 0.3264 | 0.3454 | yes | $10^{-2}$ | 1 |
| $s_{PKR*,iv}$ | $nM^{-1}$ | 1.002 | 0.8928 | 1.117 | yes | $10^{-4}$ | $10^{6}$ |
| $b_{PKR*,iv,40bp}$ | unitless | 0.3301 | 0.3270 | Inf | no | $10^{-4}$ | 0.3301 |
| $b_{PKR*,iv,100bp}$ | unitless | 0.2196 | 0.1850 | Inf | no | $10^{-4}$ | 0.2196 |
| $b_{PKR*,iv,200bp}$ | unitless | 0.4959 | 0.4704 | Inf | no | $10^{-4}$ | 0.4959 |

| Parameter | Unit | Best fit value | Confidence interval obtained from profile likelihood estimation | | identifiable | Allowed parameter interval | |
|---|---|---|---|---|---|---|---|
| | | | Lower bound | Upper bound | | Lower bound | Upper bound |
| $k_{p,on}/k_{p,off}$* | $nM^{-1}$ | 0.09159 | 0.07153 | 0.1228 | yes | $10^{-6}$ | $10^{5}$ |
| $s_{eIF2\alpha*,dsRNA}$ | $nM^{-1}$ | 0.00123 | 0.001145 | 0.001323 | yes | $10^{-3}$ | $10^{3}$ |
| $b_{eIF2\alpha*,dsRNA}$ | unitless | 0.5945 | 0.5545 | 0.6327 | yes | $10^{-6}$ | $10^{3}$ |
| $s_{eIF2\alpha*,A}$ | $nM^{-1}$ | 0.00206 | 0.001762 | 0.002422 | yes | $10^{-6}$ | $10^{5}$ |
| $b_{eIF2\alpha*,A}$ | unitless | 0.3213 | 0.2514 | 0.4307 | yes | $10^{-6}$ | $10^{5}$ |
| $s_{eIF2\alpha*,T}$ | $nM^{-1}$ | 0.001146 | 0.0009139 | 0.001381 | yes | $10^{-6}$ | $10^{5}$ |
| $b_{eIF2\alpha*,T}$ | unitless | 0.6228 | 0.5403 | 0.702 | yes | $10^{-6}$ | $10^{5}$ |
| $s_{eIF2\alpha*,G}$ | $nM^{-1}$ | 0.0008348 | 0.0008114 | 0.0008587 | yes | $10^{-6}$ | $10^{5}$ |
| $b_{eIF2\alpha*,G}$ | unitless | $1.002 \cdot 10^{-6}$ | 0 | 0.001217 | no | $10^{-6}$ | $10^{5}$ |
| $s_{GADD34,G}$ | $nM^{-1}$ | 0.009771 | 0.009426 | 0.01013 | yes | $10^{-6}$ | $10^{5}$ |
| $s_{GADD34,T,kin}$ | $nM^{-1}$ | 0.03304 | 0.0213 | 0.04735 | yes | $10^{-6}$ | $10^{5}$ |
| $s_{eIF2\alpha*,T,kin}$ | $nM^{-1}$ | 0.0008976 | 0.0008309 | 0.000921 | yes | $10^{-6}$ | $10^{5}$ |
| $b_{eIF2\alpha*,T,kin}$ | unitless | $1.002 \cdot 10^{-6}$ | 0 | 0.2009 | no | $10^{-6}$ | $10^{5}$ |
| $GADD34_{ex,1}$ | nM | 14.53 | 9.656 | 20.20 | yes | $10^{-2}$ | $10^{3}$ |
| $GADD34_{ex,2}$ | nM | 22.39 | 16.08 | 30.84 | yes | $10^{-2}$ | $10^{3}$ |
| $GADD34_{ex,3}$ | nM | 42.57 | 24.37 | 57.33 | yes | $10^{-2}$ | $10^{3}$ |
| $GADD34_{ex,4}$ | nM | 68.49 | 48.49 | 88.81 | yes | $10^{-2}$ | $10^{3}$ |
| $GADD34_{ex,5}$ | nM | 105.3 | 73.07 | 139.5 | yes | $10^{-2}$ | $10^{3}$ |
| $GADD34_{ex,6}$ | nM | 102.5 | 70.21 | 142.5 | yes | $10^{-2}$ | $10^{3}$ |
| $GADD34_{ex,7}$ | nM | 123.1 | 53.01 | 193.2 | yes | $10^{-2}$ | $10^{3}$ |
| $PKR\_act_{1}$ | $min^{-1}$ | 0.01173 | 0.007992 | 0.01728 | yes | $10^{-6}$ | $10^{5}$ |
| $PKR\_act_{2}$ | $min^{-1}$ | 0.01297 | 0.00972 | 0.01717 | yes | $10^{-6}$ | $10^{5}$ |
| $PKR\_act_{3}$ | $min^{-1}$ | 0.01468 | 0.01065 | 0.02031 | yes | $10^{-6}$ | $10^{5}$ |
| $PKR\_act_{4}$ | $min^{-1}$ | 0.01521 | 0.01082 | 0.02155 | yes | $10^{-6}$ | $10^{5}$ |
| $PKR\_act_{5}$ | $min^{-1}$ | 0.01529 | 0.01103 | 0.02127 | yes | $10^{-6}$ | $10^{5}$ |
| $PKR\_act_{6}$ | $min^{-1}$ | 0.02002 | 0.0124 | 0.03234 | yes | $10^{-6}$ | $10^{5}$ |
| $PKR\_act_{7}$ | $min^{-1}$ | 0.02137 | 0.01511 | 0.03026 | yes | $10^{-6}$ | $10^{5}$ |
| $PKR\_act_{8}$ | $min^{-1}$ | 0.02350 | 0.01374 | 0.03949 | yes | $10^{-6}$ | $10^{5}$ |
| $PKR\_act_{9}$ | $min^{-1}$ | 0.02924 | 0.01878 | 0.04582 | yes | $10^{-6}$ | $10^{5}$ |
| $PKR\_act_{10}$ | $min^{-1}$ | 0.03001 | 0.01961 | 0.04593 | yes | $10^{-6}$ | $10^{5}$ |
| $PKR\_act_{11}$ | $min^{-1}$ | 0.04581 | 0.02745 | 0.07559 | yes | $10^{-6}$ | $10^{5}$ |
| $PKR\_act_{12}$ | $min^{-1}$ | 0.04843 | 0.02905 | 0.0814 | yes | $10^{-6}$ | $10^{5}$ |
| $PKR\_act_{13}$ | $min^{-1}$ | 0.05189 | 0.03025 | 0.08741 | yes | $10^{-6}$ | $10^{5}$ |
| $PKR\_act_{14}$ | $min^{-1}$ | 0.08617 | 0.05138 | 0.143 | yes | $10^{-6}$ | $10^{5}$ |
| $PKR\_act_{15}$ | $min^{-1}$ | 0.08899 | 0.05422 | 0.1431 | yes | $10^{-6}$ | $10^{5}$ |
| $PKR\_act_{16}$ | $min^{-1}$ | 0.1344 | 0.08084 | 0.2176 | yes | $10^{-6}$ | $10^{5}$ |
| $PKR\_act_{17}$ | $min^{-1}$ | 0.4212 | 0.1064 | 0.6430 | yes | $10^{-6}$ | $10^{5}$ |
| $PKR\_act_{18}$ | $min^{-1}$ | 0.7110 | 0.1938 | 1.0627 | yes | $10^{-6}$ | $10^{5}$ |
| $PKR\_act_{19}$ | $min^{-1}$ | 1.256 | 0.3796 | 1.8443 | yes | $10^{-6}$ | $10^{5}$ |

* In absence of time-resolved p-PKR measurements, we only estimated parameters $k_{p,on}$ and and $k_{d,on}$, while keeping parameters $k_{d,off}$ and $k_{d,off}$ fixed to $1 min^{-1}$. By model simulations

for an integration time of 3 days, we enforced steady states and determined parameter ratios $k_{p,on}/k_{p,off}$ and $k_{d,on}/k_{d,off}$.
** The kinetic parameter for GADD34 mRNA degradation was fixed to $k_{deg,m} = 0.0035 min^{-1}$ as previously reported (*11*).
*** The kinetic parameter for GADD34 protein degradation, $k_{deg,GADD34} = 0.01854 min^{-1}$ was independently fitted (section 'GADD34 degradation model' in Supplementary Text) and fixed to this value for fitting the deterministic model of the integrated stress response.

**Movie S1.**

HCV-induced SG-On and SG-Off phases in Huh7 YFP-TIA1 cells treated with 100 IU/ml IFN-α.

**Movie S2.**

HCV-induced SG-On and SG-Off phases in Huh7 YFP-TIA1 cells.

**Movie S3.**

HCV-induced SG-On and SG-Off phases in Huh7 YFP-TIA1 cells overexpressing PKR.
